# Biologically Informed Variational Inference Enables Interpretable Cell Phenotyping and Discovery

**DOI:** 10.1101/2025.06.10.657924

**Authors:** Lucas Arnoldt, Julius Upmeier zu Belzen, Luis Herrmann, Khue Nguyen, Naveed Ishaque, Fabian J. Theis, Benjamin Wild, Roland Eils

## Abstract

Multi-omics technologies allow detailed characterization of cell types and states across omics layers as well as chemical and genetic perturbations. Variational autoencoders have become a cornerstone of single-cell data integration; however, they are often implemented as black-box models, requiring post hoc interpretation using known markers, pathways, or regulators to make sense of their latent representations. NetworkVI fundamentally flips this paradigm by incorporating biological knowledge directly into the model architecture. By embedding co-regulation networks derived from topologically associated domains and structured ontologies such as the Gene Ontology (GO), NetworkVI introduces a biologically-informed inductive bias that promotes the preservation of meaningful variation during integration while enforcing interpretability at both the gene and GO levels. NetworkVI achieves state-of-the-art data integration, modality imputation, and cell label transfer across bimodal and trimodal datasets. Beyond integration, here we show that NetworkVI facilitates ontology-guided hypothesis generation by exploiting established associations between genes, structured cellular programs, and regulatory domains to interpretably model cellular identities. Furthermore, decomposition of GO activation spaces resolves lineage-specific functional states within immune cell types, including quiescent, inflammatory, and transitional monocyte subpopulations, that are invisible to transcriptomic clustering. NetworkVI prioritizes GO-term programs associated with immunosenescence, consistent with age-associated immune dysregulation and reveals candidate immune evasion mechanisms consistent with CD58 loss in a Perturb-CITE-seq melanoma dataset.

## Introduction

Recent multimodal single-cell sequencing technologies have provided detailed insights into the complex cellular landscape (Baysoy *et al*. 2023, Hu *et al*. 2024). Technologies such as CITE-seq, 10x Multiome and emerging trimodal methods such as DOGMA-seq (Mimitou *et al*. 2021) and TEA-seq (Swanson *et al*. 2021), allow researchers to jointly profile the transcriptome, chromatin accessibility, and surface protein expression. Additionally, extensions such as Perturb-CITE-seq (Frangieh *et al*. 2021) incorporate CRISPR-based perturbations, enabling functional genomics screens alongside multimodal profiling. These rich datasets have uncovered regulatory mechanisms and cell states with unprecedented resolution (Baysoy *et al*. 2023, Lance *et al*. 2021). However, they present major challenges in interpretation, integration and the handling of technical noise and covariates.

Recent benchmarking studies (Hu *et al*. 2024, Fu *et al*. 2025) have reported strong performance of deep generative models such as TotalVI (Gayoso *et al*. 2021), MultiVI (Ashuach *et al*. 2023), and MIDAS (He *et al*. 2024), for denoising, imputation, and paired or mosaic integration. However, these models are typically black boxes: their latent spaces lack biological grounding, and biological insight must be recovered post hoc via clustering or gene set enrichment. At the same time, common modeling steps such as batch correction or harmonization are performed with little transparency: these procedures may remove unwanted technical variation, but they can also eliminate biologically meaningful signals such as those associated with age, patient identity, or subtle disease processes (Hu *et al*. 2024, Fu *et al*. 2025). Without interpretable architectures, it remains difficult to determine whether such covariates are nuisances or drivers of meaningful heterogeneity. Recent unimodal models such as VEGA (Seninge *et al*. 2021), expiMap (Lotfollahi *et al*. 2023), and scNET (Sheinin *et al*. 2025) incorporate domain knowledge but stop short of modeling structured regulatory hierarchies, offer limited interpretability for small-effect-size signals such as genetic perturbations, and do not extend to multimodal settings.

Incorporating biological priors directly into model architectures offers a path toward more interpretable and faithful representations of cellular states. Two underutilized yet powerful sources of prior knowledge are the Gene Ontology (GO), which encodes functional relationships between genes in a hierarchical structure, and topologically associated domains (TADs) - which capture gene co-regulation based on 3D chromatin architecture. While GO has been used for annotation or enrichment, its use as a neural inductive bias remains limited to genotype-to-phenotype prediction tasks (Ma *et al*. 2018) and unimodal RNA-seq perturbation analysis (Doncevic and Herrmann 2023). TAD-based co-regulation has not been integrated into model architectures, despite its importance for cell identity (Neems *et al*. 2016).

To address these limitations, we introduce NetworkVI, a biologically informed deep generative model that incorporates GO- or other hierarchies and TAD co-regulation networks directly into modality-specific encoders, enabling gene-, TAD-, and GO-term-level importance scores and covariate attribution through an attention mechanism, all while achieving competitive performance in established benchmarks.

## Results

### NetworkVI is a prior knowledge-driven variational autoencoder

NetworkVI is an explainable sparse deep generative model for multimodal single-cell integration (Figure 1a, Supplementary Figure 1), using modality-specific decoder modules from MultiVI (Ashuach *et al*. 2023). Each modality-specific encoder includes a TAD-informed gene co-regulation layer and a sparse GO-based neural network as introduced by (Ma *et al*. 2018) (Figure 1b), aligning the model’s internal structure with known biological hierarchies and enabling GO-term-, TAD-, and gene-level importance scores. Covariate effects are modelled via a GO-term-specific multi-head attention mechanism (Figure 1c). For query-to-reference mapping, NetworkVI adapts the scArches framework (Lotfollahi *et al*. 2022) (Figure 1d, see Methods).

**Figure 1:**
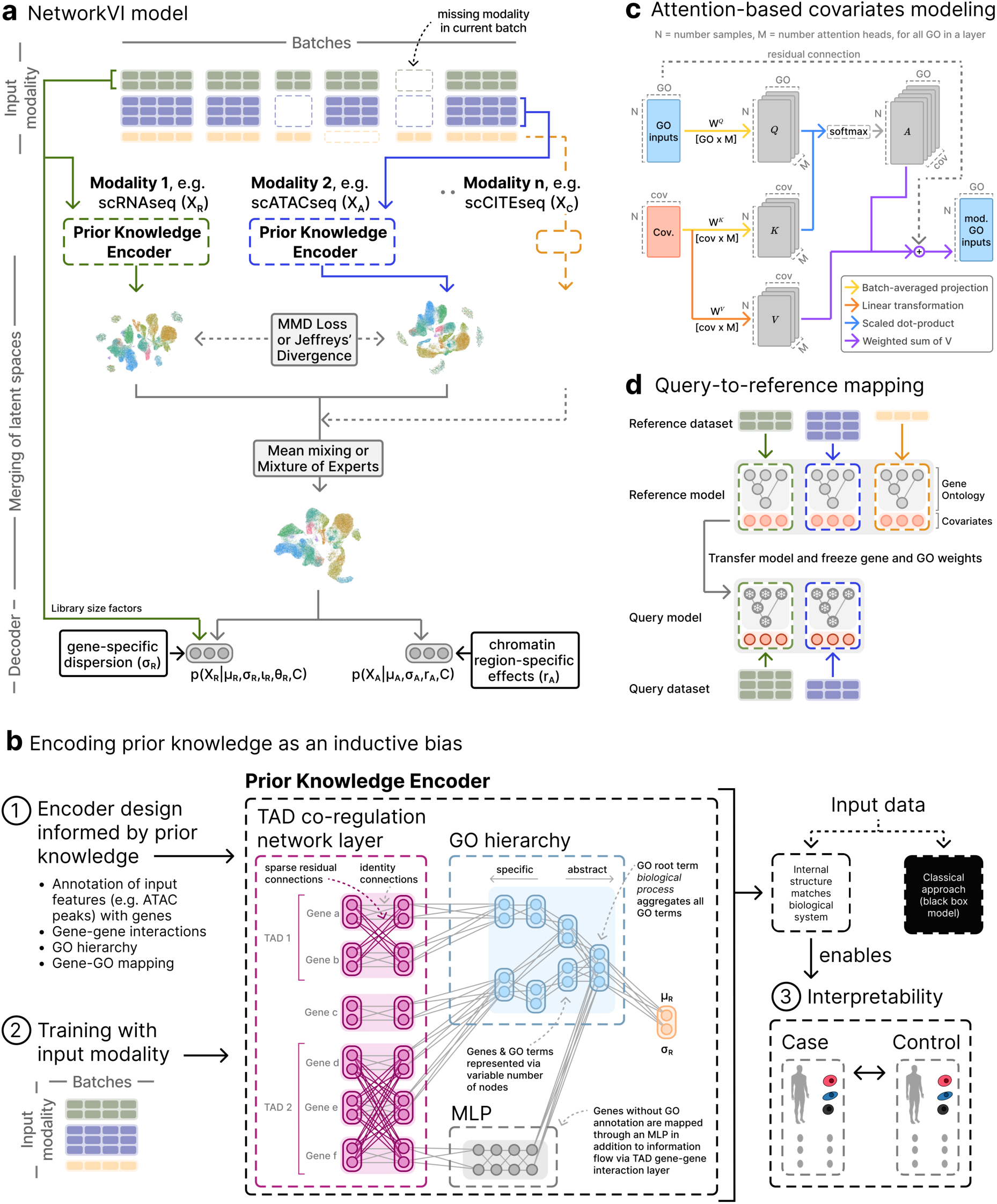
Overview of NetworkVI enabling multi-level interpretability via prior knowledge as an inductive bias. **a** Modality-specific representations are learned by the Prior Knowledge Encoder. A joint representation is generated by aggregation via mean mixing or Mixture of Experts. **b** The Prior Knowledge Encoder uses a gene TAD-informed residual connections linking co-localised genes, followed by a sparse Gene Ontology graph routing information from specific to abstract nodes. Unannotated features are mapped to the root GO biological process (GO:0008150) using a multilayer perceptron (MLP) (grey). NetworkVI offers in contrast to classic black box approaches multi-level interpretability across genes, regulatory domains (TADs), cellular programs (GO terms), and covariates. **c** GO term-specific multi-head attention mechanism for covariate modelling. Covariates serve as keys and values; GO term representations act as queries. The adjusted GO representations are returned via a residual connection. **d** NetworkVI can additionally be used for query-to-reference mapping by finetuning the GO term-specific covariate attention values while freezing all other weights in the GO graph.

For integration, we quantified performance using the scib score, which captures how well a method preserves biological signal, aligns data across modalities, and corrects for batch effects ((Luecken *et al*. 2022); see Methods). Across all tested datasets, NetworkVI achieves the highest biological variance conservation scores among benchmarked methods, with state-of-the-art integration performance in trimodal and CITE-seq settings and competitive performance for multiome data integration. In query-to-reference mapping and modality imputation, NetworkVI consistently outperforms MultiVI (Supplementary Note 1.1).

### GO Biological Process and lineage-matched TADs provide optimal integration performance and interpretability

NetworkVI is agnostic to the ontology and gene–gene interaction source used as prior knowledge: any directed acyclic graph or sparse interaction structure can be supplied at model-construction time (Figure 2a). We find that GO Biological Process and K562-derived TADs as reasonable defaults, outperforming alternatives quantitatively and providing high quality interpretability. Furthermore, randomizing the Gene-GO mapping as a negative control decreases the performance in scIB metrics, and completely removes the interpretability signal (Supplementary Note 1.2; Supplementary Figure 13).

**Figure 2:**
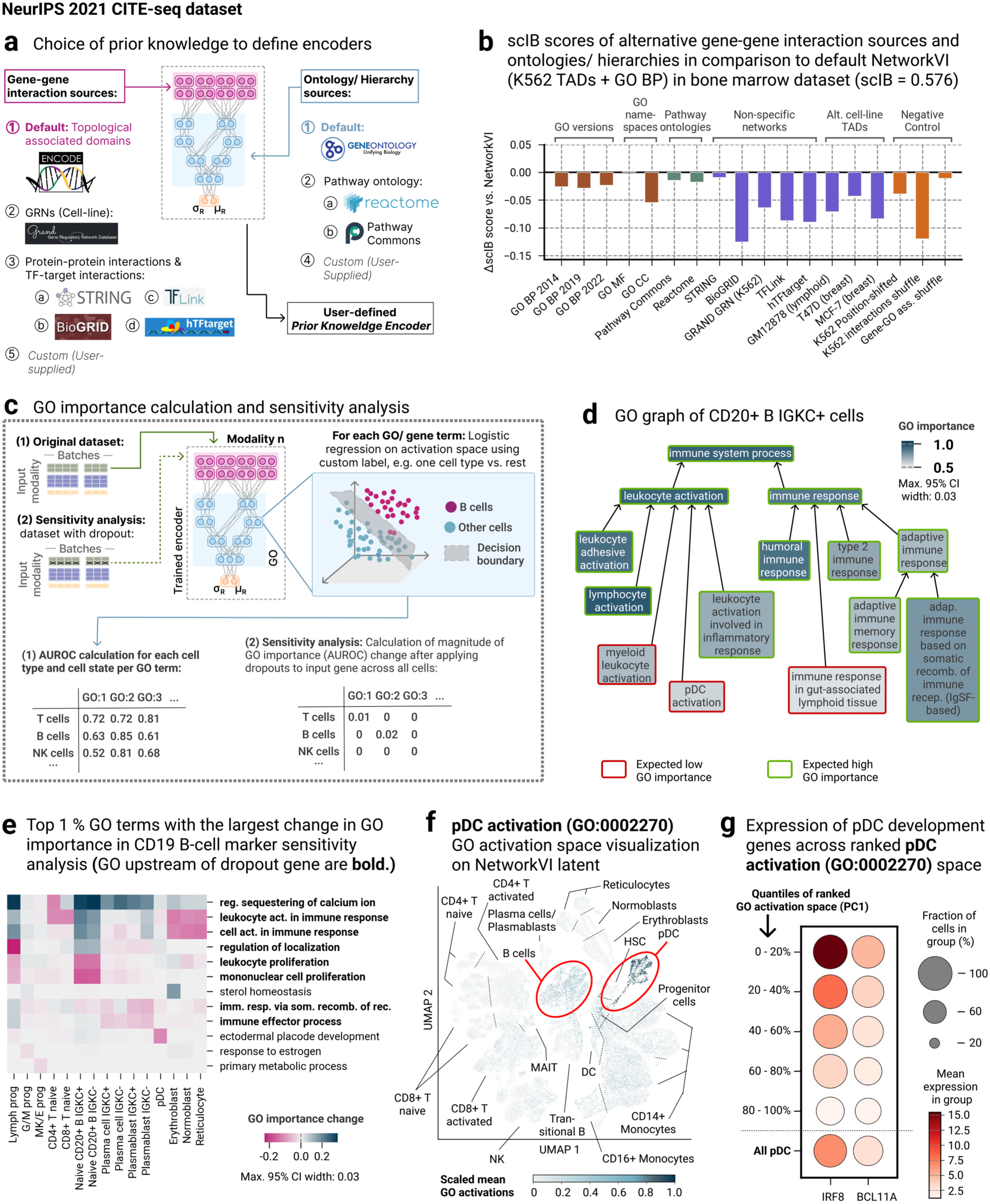
GO interpretability inferred from NetworkVI captures cell type and state characteristics. **a** Prior knowledge sources supported by NetworkVI: gene–gene interaction sources (TADs, PPIs, GRNs) and ontology/hierarchy sources (GO namespaces, Reactome, Pathway Commons). Custom sources and DAGs can be supplied. **b** ΔscIB integration score relative to the default NetworkVI configuration (K562 TADs + GO Biological Process, scIB = 0.576) for alternative prior knowledge sources on the NeurIPS 2021 CITE-seq BMMC dataset. GO versions (2014–2022) show progressively lower performance reflecting annotation growth over time. GO MF reaches near-parity; GO CC shows a larger penalty. Pathway ontologies (Pathway Commons, Reactome) show small penalties while retaining strong interpretability (≥82% of terms with AUROC > 0.55). Non-specific gene–gene interaction databases and tissue-mismatched TAD maps degrade integration more than any ontology alternative. The K562 Position-shifted TAD, a negative control, confirms that co-regulatory structure, not architectural sparsity, drives the TAD benefit. **c** Schematic of GO importance calculation (1) and sensitivity analysis (2). GO importance is computed via logistic regression on GO node activations (AUROC). Sensitivity analysis sets input features to zero and quantifies the resulting GO importance change. **d** GO interpretability was calculated for naive CD20⁺ B IGKC⁺ cells against cell types as labels, revealing the specific GO terms that allow NetworkVI to correctly identify the immunological role of naive B cells. For example, the GO term humoral immune response (GO:0006959) exhibits a high GO Importance, whereas pDC activation (GO:0002270) demonstrates a low GO Importance. GO importance scores are stable across bootstrap replicates; the maximum 95% CI width across all GO terms shown is 0.03. **e** NetworkVI correctly recapitulates known marker genes, as shown by a sensitivity analysis for the B-cell marker gene CD19, for which GO terms were selected by the magnitude of their GO importance change. A significant change in the importance of GO terms linked directly to the marker gene or via parent GO terms (upstream of GO terms associated with dropout gene) is observed. For instance, as expected, the absence of CD19 increases the importance of the GO term mononuclear cell proliferation (GO:0032943) for B cells and lymphoid progenitors. **f** UMAP visualization of the latent space generated by NetworkVI, color-coded by the averaged activations of the GO term pDC activation (GO:0002270). Specific relevant cell types, such as pDC and B cells can be clearly distinguished visually via the activations. **g** pDC cells were stratified into five quantiles based on their activation space score for the GO term pDC activation (GO:0002270). Shown are the average expression levels of two key transcription factors across these quantiles: IRF8, which directly feeds into the GO node, and BCL11A, which are functionally linked but not directly connected in the GO graph. IRF8 and BCL11A exhibit a strong gradient with activation level, suggesting activation levels capturing transient cell states.

### Enhancing biological interpretability via GO importance

NetworkVI is inherently interpretable due to the domain knowledge embedded in its modality-specific encoders. We use GO importance (ranging from 0.5 – 1.0) as a measure, which quantifies how well the representations of individual GO terms in NetworkVI predict cell-level properties, such as cell type or patient disease status (Figure 2c (1), Supplementary Figure 14a, see Methods). Since these activations come from modality-specific encoders, GO importance is computed separately for each modality, allowing an assessment of how different data modalities contribute to predicting cell properties. This level of interpretability is especially valuable for characterising systematic differences in gene expression and -regulation across and within cell types, where current black-box multimodal integration models do not provide meaningful biological insights between the gene and cell-level (Ribeiro *et al*. 2016, Lipton 2017). GO importance scores are stable across bootstrap replicates (max 95% CI width: 0.03) and robust to training stochasticity and dataset size (see Methods, Supplementary Note 2.1). Throughout, GO importance scores, TAD importance, and covariate attention values quantify learned associations between biological programs and cell states; they do not establish causal relationships and should be interpreted as structured hypotheses for downstream experimental validation.

We computed GO importance for naive CD20^+^ B IGKC^+^ cells in the CITE-seq bone marrow mononuclear cell (BMMC) dataset (Luecken *et al*. 2021) as a positive control, since the expected pattern (high: humoral/adaptive immunity terms; low: myeloid and pDC terms) is well established. NetworkVI recapitulates this pattern for naive B cells (Figure 2d): broad immune terms (*immune system proces*s (GO:0002376), *leukocyte activation* (GO:0045321)) show high importance, while terms specific to other lineages (*myeloid leukocyte activation* (GO:0002274), *pDC activation* (GO:0002270)) show low importance. Among more specific terms, *humoral immune response* (GO:0006959) and *adaptive immune response based on somatic recombination of immunoglobulin superfamily receptors* (GO:0002208) are particularly elevated, the latter distinguishing naive B cells from more differentiated subtypes in which somatic recombination has already occurred. Low importance of *gut-associated lymphoid tissue immune response* (GO:0002387) is consistent with the bone marrow origin of this dataset. GO importance also enables global comparison of cell types across modalities; each modality encoder learns distinct yet biologically concordant cell-type representations (Supplementary Note 2.2).

### Sensitivity analysis enables evaluation of Gene-GO associations

To evaluate gene–GO associations at single-gene resolution, we performed a sensitivity analysis by setting individual marker genes to zero across all cells and quantifying the resulting cell type-specific changes in GO importance (see Methods; Figure 2c(2), Supplementary Figure 17).

We first targeted CD19, a B-cell marker that is absent in plasma cells and used to distinguish lymphocyte progenitors, and identified the GO terms with the largest magnitude of importance change (Figure 2e, Supplementary Figure 17a). While most of these terms are upstream of the removed gene, NetworkVI’s interpretability does not prioritize all upstream terms equally, as their importance depends on their utility in determining cell identity. The absence of CD19 most significantly increased the GO importance for the term *regulation of sequestering of calcium ion* (GO:0051282; 0.32, 95% CI: 0.30–0.34), consistent with known differences in calcium signalling between lymphocyte progenitors and differentiated B cells (Oh-hora and Rao 2008, Scharenberg *et al*. 2007), followed by a decrease in *mononuclear cell proliferation* (GO:0032943; -0.17, 95% CI: 0.15–0.19). These findings illustrate how NetworkVI identifies relevant GO terms that are not necessarily immediately upstream of genes of interest but can also emerge from co-regulation networks or interactions within the model. Further results of the CD19 sensitivity analysis, covering erythroblast progenitors and a CD4⁺ T cell subset, are detailed in Supplementary Note 2.3. Similarly, removing further marker genes such as CD4, CD8 or CD38 (Supplementary Figure 17b-d), caused the most substantial GO importance changes in terms directly linked to the respective marker gene or its parent GO terms.

### GO term activations resolve activated pDC cell states

While sensitivity analysis highlights how individual genes influence functional representations through *in silico* perturbations, it offers only one perspective on interpretability. We therefore also examined unperturbed GO term activations directly, by color-mapping averaged activations for selected GO terms onto the UMAP embedding of the NetworkVI latent space (CITE-seq BMMC dataset; Figure 2f). GO terms show distinct activation patterns across cells, reflecting both discrete boundaries between cell types and subtle gradients within them. Cells close in latent space can show divergent GO activation profiles, reflecting biologically meaningful intra-type heterogeneity.

Although *pDC activation* (GO:0002270) has a low GO importance when distinguishing CD20⁺ B IGKC⁺ cells (Figure 2d), consistent with its limited discriminative role in this context, and with the known modest regulatory influence of pDCs on naïve B cells (Bekeredjian-Ding et al. 2005), it produces a *pDC activation*-specific pattern in the UMAP embedding (Figure 2f), partly explaining their separation from other cell types. Ranking pDCs by their *pDC activation* score (PC1) reveals a strong expression gradient for IRF8 (Spearman r = 0.90, 95% CI: 0.87– 0.93, p = 2.7×10⁻⁹; Sichien et al. 2016), one of only two genes directly connected to this GO node, and a moderate gradient for BCL11A (r = 0.43, 95% CI: 0.41–0.45, p = 1.8×10⁻⁶; (Ippolito *et al*. 2014)), which connects via shared parent terms (Figure 2g; Supplementary Note 2.3). To further validate the IRF8– *pDC activation* link *in silico*, we set IRF8 expression to zero across all cells and recomputed GO importance changes. *Plasmacytoid dendritic cell activation* (GO:0002270) appeared among the top 2% of GO terms with the largest importance change, consistent with IRF8 being a key contributor to this term’s discriminative power in pDCs (Supplementary Figure 17e). These results indicate that GO activation levels reflect continuous, biologically meaningful transcriptional states rather than discrete cell-type boundaries.

### GO-based decomposition maps functional substates across innate and adaptive immune populations

Screening all modeled GO terms per cell type on their 2D activation space (see Methods), most produce no distinguishable clusters, in line with prior expectation. In CD14^+^ monocytes, *T cell antigen processing and presentation* (GO:0002457) ranks highest on silhouette score, while simultaneously showing one of the lowest GO importance scores (0.47, 95% CI: 0.465–0.475), making it an ideal example of sub-state information invisible at the transcript level. HDBSCAN on the activation space identified three clusters within CD14⁺ monocytes: a quiescent majority (86%) and two transcriptionally distinct substates comprising 7% and 5% of cells respectively (Figure 3a). We labeled the 7% substate ICAM1-inflammatory because ICAM1 showed the largest fold change relative to the quiescent substate. The 5% substate was labeled transitional-inflammatory based on elevated FCGR3A (CD16a) and CX3CR1, consistent with a CD14^+^CD16^+^ transitional monocyte identity. The substates were not separable in the transcriptional latent space (NetworkVI, Figure 3b) and were robust across donors and batches (Supplemntary Figure 18a). Analogous substates for GO:0002457 emerged within CD16⁺ monocytes (Supplemntary Figure 18b), suggesting shared inflammatory programs, and underlining NetworkVIs robustness.

Recomputing GO importance for GO:0002457 per substate showed that NetworkVIs signal for antigen-presentation activity is concentrated in activated states: near-random importance for the quiescent subset (0.54, 95% CI: 0.52–0.56) versus near-maximum scores for ICAM1-inflammatory (0.99, 95% CI: 0.989–0.991) and transitional-inflammatory (0.94, 95% CI: 0.938–0.942) cells, with a similar pattern in CD16⁺ monocytes (Figure 3c, Supplementary Figure 18c). Both activated substates upregulated CX3CR1, CCL3, and IL1B, with transitional cells additionally expressing FCGR3A (CD16a), consistent with partial acquisition of non-classical identity (Figure 3d), notably without direct GO annotation to this term.

**Figure 3:**
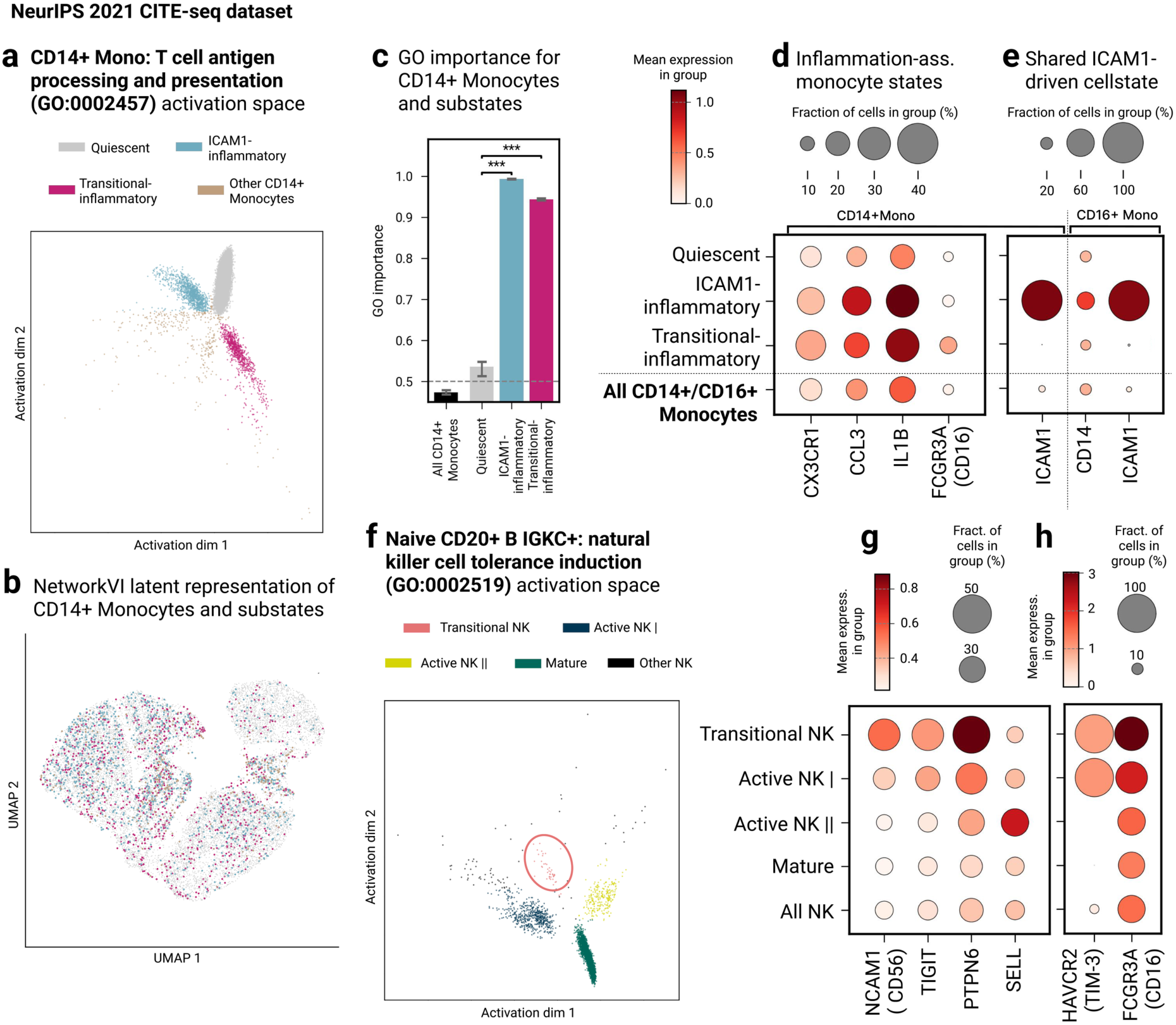
Functional GO decomposition reveals antigen presentation and tolerance induction substates across immune cell types. **a** GO activation space of T cell antigen processing and presentation (GO:0002457) of CD14+ monocytes, colored by substate: quiescent, ICAM1-inflammatory, Transitional-inflammatory. **b** UMAP of transcriptional latent space (NetworkVI) showing no spatial segregation of CD14+ Monocyte substates. **c** GO importance analysis reveals near-random signal for all CD14⁺ monocytes but high discriminability within GO-active subsets. Bootstrapped AUROC distributions (n=200 resamples) for each group. Bars indicate median AUROC; error bars denote 95% bootstrap confidence intervals. Statistical comparisons between groups were performed using two-sided DeLong test on bootstrap distributions (***=p<0.001). **d** Inflammatory substates show elevated expression of CX3CR1, CCL3, IL1B; transitional-inflammatory cells additionally express FCGR3A (CD16a). **e** Dot plot of substate-defining genes ICAM1, highlighting molecular distinctions between inflammatory and transitional substates. **f** GO activation space of NK cell tolerance induction (GO:0002519) of NK cells, colored by substate: Transitional NK, Active NK I, Active NK II, and Mature NK. **g** Dot plot of substate-defining genes NCAM1 (CD56), TIGIT, PTPN6, SELL, highlighting transitional versus differentiated NK phenotypes. **h** Dot plot of HAVCR2 (TIM-3), and FCGR3A defining Transitional NK and Active NK I substates.

The ICAM1-inflammatory state, present in both monocyte subsets (Figure 3e), likely reflects a stimulus-induced, reversible activation program. ICAM1 facilitates endothelial adhesion critical for MHC–TCR interactions (Lebedeva *et al*. 2004). A detailed comparison of the two activated substates is provided in Supplementary Note 2.4. Standard Louvain clustering and GRN/regulon-based workflows did not recapitulate these substates, nor did a shuffled gene–GO control (Supplementary Figure 13a), confirming they are not captured by transcription factor-centric programs (Supplementary Note 2.4; Supplementary Figure 18d,e).

The same screen identified *natural killer cell tolerance induction* (GO:0002519) as the top-ranked NK-specific GO term by silhouette score, resolving four substates (Transitional NK, Active NK I, Active NK II, and Mature NK), whose full marker characterisation and comparison with (Yang *et al*. 2019) is provided in Supplementary Note 2.5 (Figure 3f–h).

### Explicitly Modelling TAD Co-Regulation Enhances Interpretability and Functional Resolution in NetworkVI

NetworkVI’s TAD co-regulation layer operates through sparse residual connections among genomically co-localized genes. We provide a specific analysis of gene-pairs with the highest residual connection weights in Supplementary Note 2.6. We also compare the full NetworkVI model against an ablated version in which the TAD layer is removed (Figure 4a). Across all cell types in the transcriptomic modality, the full model exhibits a 4.5% higher average GO importance (95% bootstrap CI: 3.5–5.6%; Wilcoxon signed-rank test across phenotypes, p = 3.3×10⁻⁹), suggesting that enhanced graph connectivity from unannotated genes through TADs contributes to improved functional signal capture. This improvement is broadly distributed across the GO hierarchy: for CD14⁺ monocytes, 75.7% of GO terms (1,356 of 1,791) show a positive importance change upon addition of the TAD co-regulation layer (Supplementary Figure 19a). In the CITE-seq BMMC dataset (4,000 genes across 737 TADs), at least one unannotated gene (otherwise modelled through an MLP) is found in one of the 262 TADs. These residual pathways reduce the fraction of unannotated genes by 31%, re-integrating 279 previously unmapped genes (from an initial pool of 887) into the modeled hierarchy.

**Figure 4:**
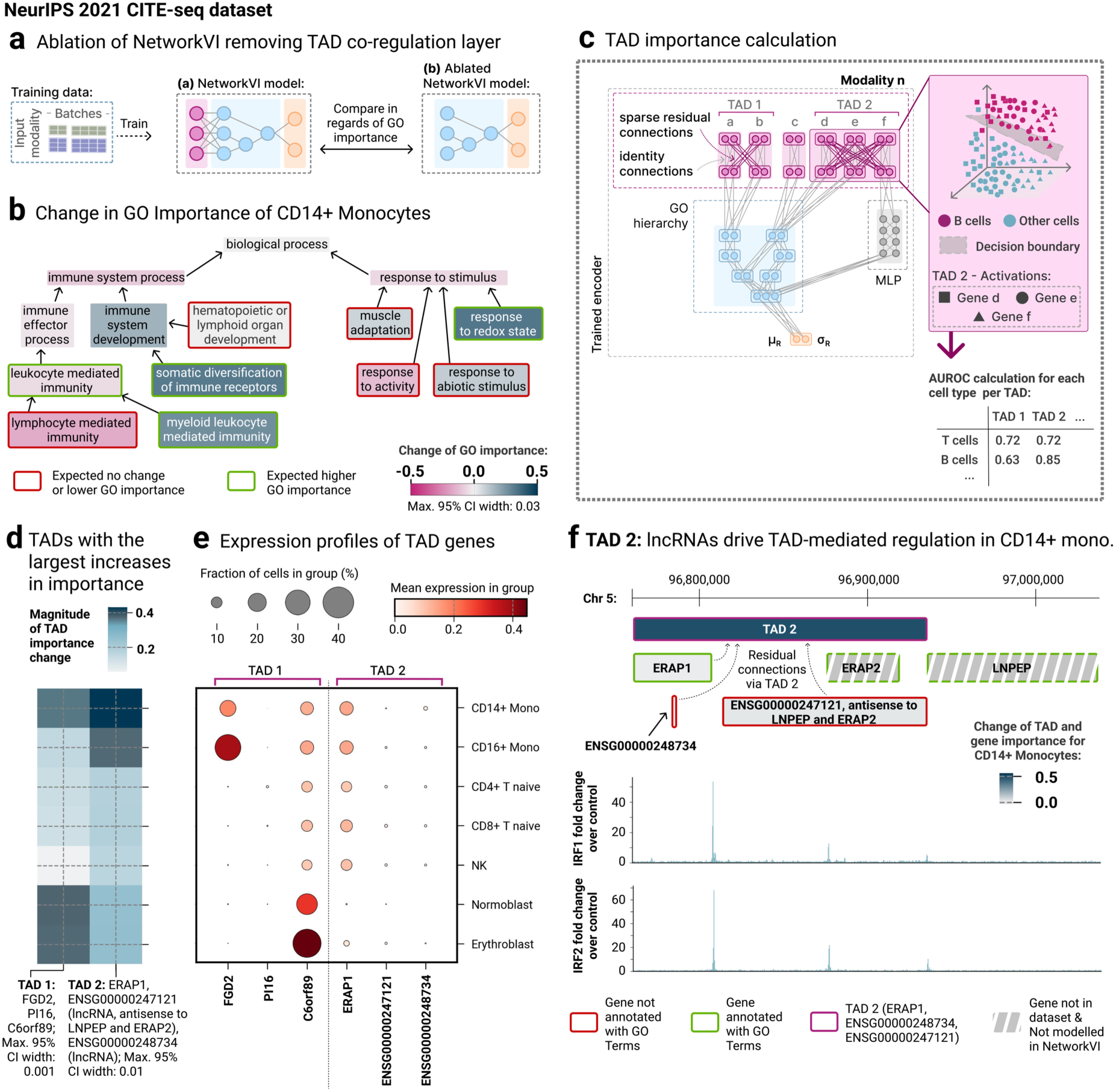
Modelling TAD co-regulation improves GO-level interpretability and recovers lncRNA regulation mechanisms. **a** We here perform a comparison of the full NetworkVI model, incorporating the TAD-informed co-regulation layer, with an ablated variant lacking this layer in regards of the GO importance, here for **b** CD14⁺ monocytes. Specific immune-related GO terms like myeloid leukocyte mediated immunity (GO:0002444) increase in importance, while generic terms like response to stimulus (GO:0050896) and non-specific GO terms decrease in importance. Overall GO importances increase by 16.9%. The stronger interpretability capabilities of NetworkVI with the TAD co-regulation layer can be traced back to the effective integration of 31% more previously unannotated genes into the GO hierarchy. **c** Schematic of residual connections within the co-regulation layer, enabling information flow between genes and thus through the GO graph. Via concatenating the representations of several genes and fitting a logistic regression, the importance of TADs can be inferred similarly to the calculation of GO importance. **d** Two exemplary TADs selected from the TADs with the 10% highest magnitude of changes in importance have been selected to demonstrate how the TADs capture cell-type-specific patterns. A TAD containing the genes FDG2, PI16 and C6orf90 show improvement of TAD importance for CD14+ Monocytes, Normoblast and Erythroblasts. Another TAD containing ERAP1 and two lncRNA genes without GO annotations shows a TAD improvement specific for Monocytes. TAD importance scores represent AUROC values computed from concatenated gene-level activations within each TAD. The maximum 95% CI width across all TAD importance values shown is 0.001 for TAD 1 and 0.01 for TAD2. **e** While FGD2 and PI16 are not highly expressed in Normoblasts and Erythroblasts, C6orf89 is, which indicates a role of C6orf89 where the co-regulatory context amplifies the TAD importance. In the second TAD, the expression profile does not hint at a specific co-regulatory role of the two unannotated genes ENSG00000247121 and ENSG00000248734. **f** However the gene importances of each gene of this TAD individually only increase slightly after the introduction the co-regulation layer and, **w**hile the TAD importance increases 20-fold, indicating co-regulatory interactions learnt here. The regulatory role of the unannotated gene ENSG00000247121 can be attributed to the gene being antisense to LNPEP and ERAP2 and co-location with IRF1 and IRF2. ENSG00000248734 is implicated to play a key role in CD14+ Monocytes, but not CD16+ Monocytes. The specificity of the TAD co-regulation layer is further established in Figure 2b: tissue-mismatched TAD maps from breast cancer cell lines reduce both integration performance and GO importance scores, while the position-shifted K562 control confirms that genuine co-regulatory groupings, not sparse architectural connectivity alone, drive the observed deltas.

GO importance deltas across all terms are particularly pronounced in myeloid cell types, with CD14⁺ and CD16⁺ monocytes showing increases of 16.9% (95% CI: 15.8–17.9%; Wilcoxon p = 8.0×10⁻¹³²) and 11.1% (95% CI: 10.1–12.1%; Wilcoxon p = 2.1×10⁻⁸⁶), respectively. By contrast, some hematopoietic progenitor cells exhibit smaller or negligible changes. We analyzed GO importance shifts in CD14⁺ monocytes (Figure 4b). Broad GO terms such as *immune system process* (GO:0002376) showed reduced importance, likely reflecting earlier combination of gene-level information in the GO graph. More specific, lineage-relevant GO terms such as *myeloid leukocyte mediated immunity* (GO:0002444) increase in importance. Conversely, unrelated terms such as *lymphocyte mediated immunity* (GO:0002449) showed diminished importance, enhancing the cell-type specificity of interpretations.

In parallel, NetworkVI enables quantification of individual gene-level and TAD-level importances (Figure 4c, see Methods). The gene importance increases by 3.2% (95% CI: 3.0–3.5%) across all cell types; the TAD-level importance by 1.5% (95% CI: 1.4–1.7%), with lymphoid cells exhibiting the largest increases. This is especially notable given that the TAD reference maps are derived from K562 (myeloid lineage), suggesting the model successfully adapts cross-lineage structural priors in a context-specific manner. Tissue-mismatched TAD maps from T47D and MCF7 reduce mean gene importance by 1.4% and 0.9% and TAD-group importance by 2.5% and 1.9% respectively, while the lineage-matched GM12878 map yields a modest increase (+1.3% gene, +1.9% TAD-group). The position-shifted K562 negative control shows no meaningful change (<0.03%), confirming that biologically coherent co-regulatory groupings, not architectural sparsity, drive the observed deltas (full CIs in Supplementary Note 2.7).

To identify the most impactful TADs, we compared their importance between models with and without the co-regulation layer. Genes annotated as immune cell markers from PanglaoDB (Franzén *et al*. 2019) have, on average, 8.7% higher importance scores than unannotated genes (95% CI: 8.1–9.2%; Wilcoxon p = 5.7×10⁻¹⁴). While TADs including marker genes show modest (on average 1%) higher importances compared to TADs excluding marker genes, their presence corresponds to a 25% average increase in TAD importance relative to the ablated model, indicating a central role for cell-type markers in shaping TAD-driven co-regulation. None of four potential confounders investigated (TAD size, Hi-C contact domain score, HVG normalised dispersion, CpG island density) showed a significant association (Supplementary Figure 19b–e). Orthogonal validation via ENCODE K562 ChIP-seq signals confirms the full model shows consistently higher correlations with the chromatin binding landscape than the ablated baseline (mean Spearman ρ = 0.022 vs −0.017; Wilcoxon p < 10⁻³⁰⁰; Supplementary Figure 21; Supplementary Note 2.8).

The TADs with the largest importance increases display distinct, cell-type-specific profiles, suggesting the model learns to exploit regulatory architecture in a lineage-aware fashion. Full characterization including two exemplary TADs, one containing protein-coding genes with partial GO annotation (FGD2/PI16/C6orf89) and one containing lncRNAs alongside a single annotated gene (ERAP1), is provided in Supplementary Note 2.9 and Figure 4d–f. Negative controls confirm the model does not uniformly inflate TAD importances: the ten TADs with the largest mean importance decreases include functionally unrelated gene pairs and one informative case (DYRK2/IFNG-AS1) where the lncRNA lacks co-regulatory signal because IFNG itself is absent from the dataset (Supplementary Note 2.10; Supplementary Figure 23).

### GO term-specific attention captures aging effects for T cells

We explored covariate attention values, which enable covariate attribution at the GO term level (Figure 5a), training NetworkVI on the CITE-seq BMMC dataset using collection site, smoking status, BMI, and age as covariates (Figure 5b, Supplementary Figure 24). Screening the top and bottom 1% of aggregated age-attention values across all GO terms identified clusters of similar cell types; notably, GO terms *regulation of metabolic process* (GO:0019222; mean attention 0.468, 95% CI: 0.462–0.474), *regulation of signaling* (GO:0023051; mean attention 0.514, 95% CI: 0.506–0.521), and *regulation of immune system process* (GO:0002682; mean attention 0.401, 95% CI: 0.391–0.410) showed strong attention to age in CD4+ activated integrin B7+ T cells, with similar patterns observed across other lymphocytic effector cells (ILC, MAIT, NK). This is consistent with thymic involution, reduced proliferative capacity, and immunosenescence (Mittelbrunn and Kroemer 2021).

To understand why *regulation of immune system process* (GO:0002682) strongly captures age effects in T and NK cells, we examined genes mapped to this term with age-correlated expression (Spearman’s ρ > 0.1, α < 0.05) showing lymphocyte-specific patterns. Notably, this set includes CCL4, CCL5, CST7, KLRB1, and KLRD1, genes previously linked to immunosenescence. CCL4 and CCL5 are upregulated during late T-cell activation (Mittelbrunn and Kroemer 2021), while age-related changes in KLRB1 and KLRD1 are well documented (Chen *et al*. 2013). These findings suggest that GO term-specific attention reflects molecular hallmarks of immune aging. Further discussion of additional cell types and covariates are detailed in Supplementary Note 2.11.

**Figure 5:**
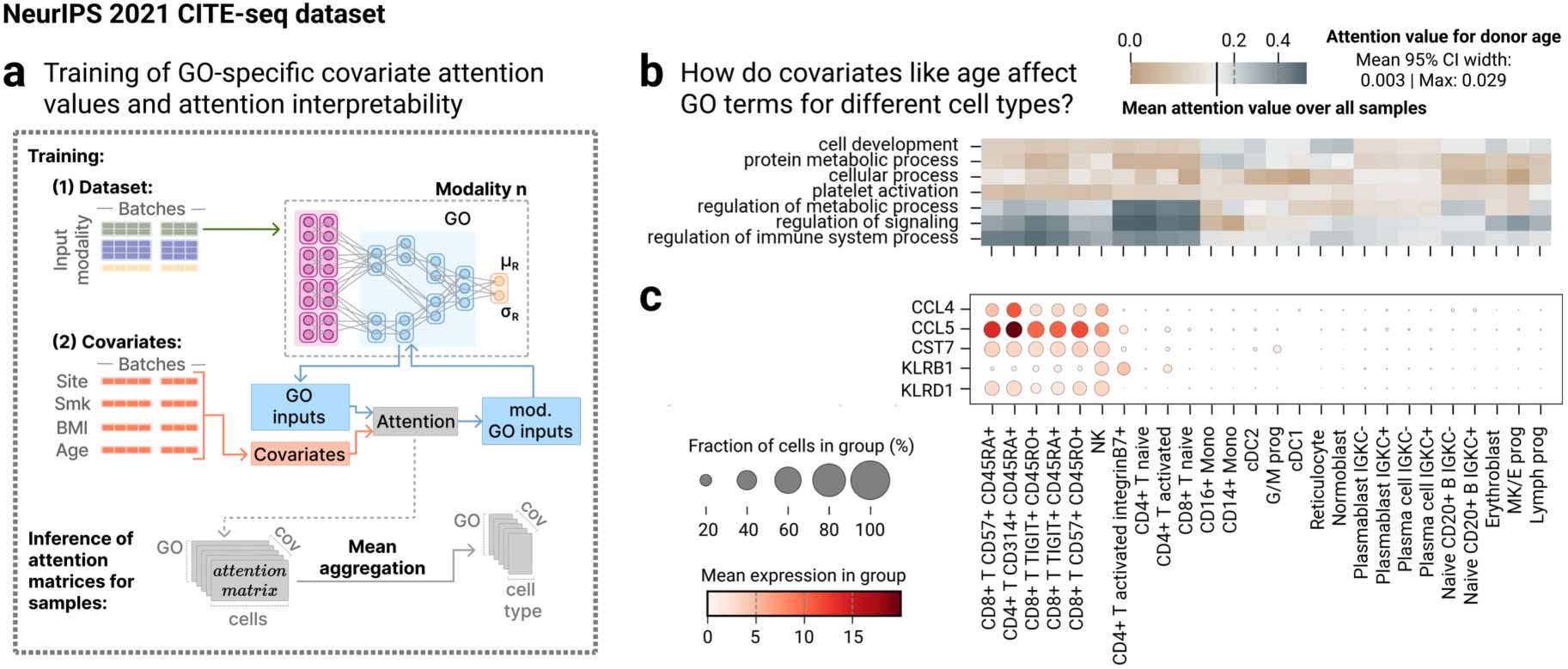
GO term-specific attention values allow for inferring aging effects in lymphocytic effector cells. **a** Schematic of GO-specific covariate attention mechanism during training. Covariates modulate GO term embeddings; original representations are preserved via residual connection. Attention values are averaged per cell-type label. **b** For the CITE-seq BMMC dataset high attention weights for the GO terms regulation of metabolic process (GO:0019222), regulation of signaling (GO:0023051), and regulation of immune system process (GO:0002682) indicate an association between the covariate donor age and lymphocytic effector cells. Covariate attention values represent means across cells within each cell type, with 95% confidence intervals computed via normal approximation. Attention values are stable across cells; the mean 95% CI width across all GO terms and cell types shown is 0.003 and the maximum 95% CI width is 0.029. **c** Age-correlated genes mapped to the GO term immune system process (GO:0002682) - including CCL4, CCL5, CST7, KLRB1, and KLRD1 - reflect known features of immunosenescence.

### GO importance provides insights into CRISPR-mediated perturbation effects

We trained NetworkVI on unperturbed cells from a melanoma Perturb-CITE-seq dataset (200,000 cells; (Frangieh *et al*. 2021)); as with other methods, perturbed and unperturbed cells are not separable in the latent space (Figure 6a), so GO importance was computed by comparing them within each screen for each perturbation gene separately (Figure 6b). We evaluated the perturbation of CD58, a glycoprotein involved in T cell adhesion, recognition and activation through CD2 receptors (Zhang *et al*. 2021). Analyzing cells pretreated with IFNγ and then cocultured with tumor-infiltrating lymphocytes (TILs), we observed high importances of several GO terms related to immune responses and antigen processing and presentation (Figure 6c, Supplementary Data 9). Notably, GO terms such as *T cell selection* (GO:0045058) and *leukocyte activation involved in immune response* (GO:0002366) were of high importance, consistent with the role of CD58 in T cell activation and cancer immune evasion (Zhang *et al*. 2021), which is MHC-independent and associated with CD58 loss (Frangieh *et al*. 2021, Ho *et al*. 2023). In the IFNγ-only and control screens, by contrast, high-importance terms were non-specific or broadly immune-related (e.g. *somatic diversification*).

**Figure 6:**
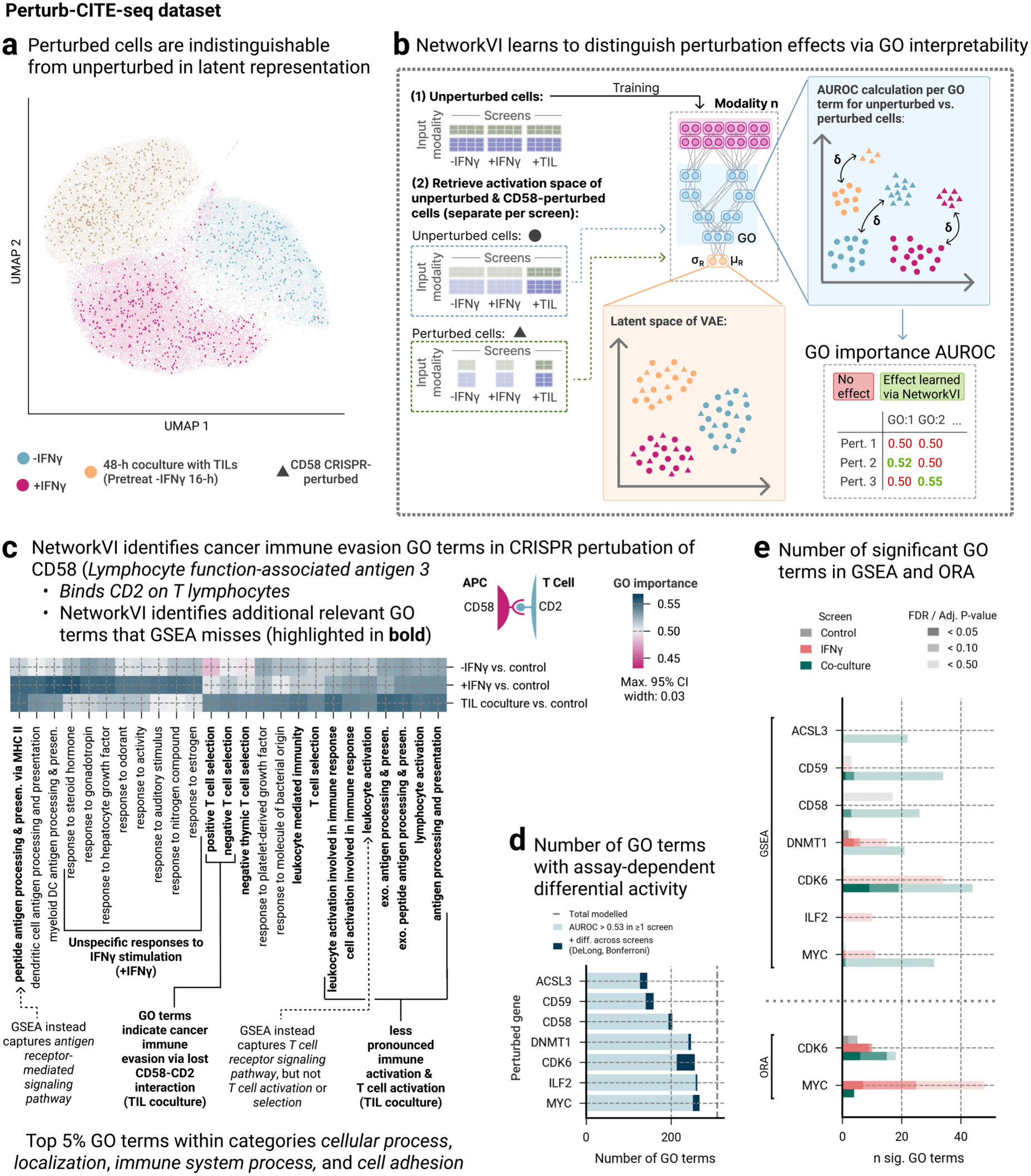
GO interpretability identifies plausible patterns of cancer immune evasion in CRISPR-mediated CD58-perturbed melanoma cells missed by GSEA. **a** UMAP visualization of the latent space from the Perturb-CITEseq dataset by (Frangieh et al. 2021) generated by NetworkVI shows that, similar to other methods, unperturbed and perturbed cells - such as those with a CD58 knockout - are not easily distinguishable in this space. **b** We trained NetworkVI exclusively on unperturbed cells and then calculated the GO importances by comparing perturbed and unperturbed cells for a single perturbation condition within a screen (-IFNγ, +IFNγ, and TIL coculture). **c** The highest GO importance scores for CRISPR-mediated CD58-perturbed cells in -IFNγ, +IFNγ and TIL coculture screens illustrate that NetworkVI highlights GO programs consistent with CD58’s role in T cell activation via GO interpretability. Immune-related GO terms, such as leukocyte activation involved in immune response (GO:0002366; 0.55, 95% CI 0.54-0.56) and T cell selection (GO:0045058; 0.54, 95% CI 0.53-0.55), together with terms describing antigen processing and presentation (GO:0019882; 0.55, 95% CI 0.54-0.55) and exogenous antigen processing and presentation (GO:0002478; 0.54, 95% CI 0.53-0.56), are of high importance in the TIL coculture screen exclusively, consistent with CD58’s role as a CD2 ligand on antigen-presenting cells. GSEA identifies 34 GO terms as significantly enriched, but these do not consistently exhibit screen-specific enrichment and often encompass unrelated biological processes; no GSEA-significant terms are absent from the model. The bold GO terms are missed by GSEA. No ORA terms reach significance for the CD58 TIL coculture condition. GO importance scores represent AUROC values comparing perturbed and unperturbed cells within each screen. The maximum 95% CI width across all GO importance values shown is 0.03. Bold GO term labels indicate terms not identified by GSEA. **d** Number of GO terms showing statistically significant (DeLong test) differential activity across the three assay conditions (−IFNγ vs. control, +IFNγ vs. control, TIL coculture vs. control) for each perturbation gene. A GO term is classified as differential if at least one pairwise z-test between conditions is significant at α = 0.05, using the 95% bootstrap CI to derive standard errors. The dotted line indicates the maximum number of modelled GO terms considered. **d** Number of GO terms showing signal across the three assay conditions (−IFNγ vs. control, +IFNγ vs. control, TIL coculture vs. control) for each perturbation gene, stratified by evidence tier: AUROC > 0.53 in any screen (light; above the upper bound of the detected 95% CIs, indicating detectable GO-level signal); differentially active across screens by CI non-overlap (dark, screen-specific signal). A GO term is classified as differentially active if at least one pairwise two-sample DeLong test between conditions is significant after Bonferroni correction for three comparisons (corrected α = 0.017), with standard errors derived from the 95% bootstrap CI half-widths. The dotted line indicates the maximum number of modelled GO terms considered. **e** Corresponding bars for GSEA (FDR thresholds 0.05/0.10/0.25) and ORA (adjusted p-value thresholds 0.001/0.01/0.05) across all three screens. The low absolute AUROC values (∼0.50–0.57) reflect the modest transcriptional effect of individual CRISPR perturbations on NetworkVI’s GO-level representations, and the small number of differential terms per gene reflects the conservative nature of significance testing at this effect size.

We conducted a separate GSEA enrichment analysis as a baseline (GO terms missed by GSEA but captured by NetworkVI are bold in Figure 6c, Supplementary Figure 25b), identifying similarly distinct functional groups across all screens, though several terms were non-specific. One of two significantly enriched GO terms linked to the cell-cell adhesion function of CD58 is *cell-matrix adhesion* (GO:0007160), which describes a mechanism that does not involve CD58 (Frangieh et al. 2021, Ho et al. 2023). GSEA distributes enrichment signals non-specifically across conditions and returns non-immune terms (e.g., *cellular response to hormone stimulus*) with comparable enrichment scores in screens lacking TIL exposure. Conversely, NetworkVI identifies immune activation and antigen presentation terms specifically concentrated in the TIL coculture condition. Meanwhile ORA does not identify any significantly enriched GO term within these four categories for the CD58 TIL coculture condition, illustrating a fundamental limitation of threshold-based enrichment in low-effect-size perturbation settings: when transcriptional effects are distributed across many genes at modest magnitude, no individual gene clears the significance threshold required by ORA, and GSEA permutation-based statistics lose power. This illustrates NetworkVIs superior capability to highlight concrete, condition-specific processes, even in low-powered, small effect-size settings.

To assess generality, we extended the GO importance analysis to six perturbation genes available in the dataset (Figure 6d; Supplementary Note 3, Supplementary Figure 26). A substantial number of GO terms are significant in at least one screen, most in CDK6 and CD58. By contrast, GSEA identifies few or no significant GO terms (FDR < 0.50) for the TIL coculture condition across most perturbation genes, and ORA reaches significance only for CDK6 and MYC (Figure 6e). In NetworkVI, CDK6 shows the clearest screen-specific signal, with antigen presentation and T cell selection terms concentrated in the TIL coculture condition, consistent with its documented role in repressing MHC class I genes (Jerby-Arnon et al. 2018), whereas GSEA distributes these terms more evenly across conditions and returns additional non-specific terms including wound healing and DNA damage processes. For genes with broader transcriptional effects (MYC, ACSL3, ILF2, DNMT1), neither method localises a sharp signal; NetworkVI reflects this transparently via near-baseline GO importance values rather than false-positive signals. While individual CRISPR perturbations produce distributed, low-magnitude transcriptional effects that are systematically underpowered for enrichment-based methods, they are detectable through NetworkVI’s nonlinear GO graph aggregation.

## Discussion

NetworkVI is an inherently interpretable state-of-the-art model for single-cell phenotyping and ontology-guided hypothesis generation. Unlike conventional approaches that apply biological interpretation only after model training, by overlaying known markers, pathways, or regulators, NetworkVI embeds the GO and TAD co-regulation networks directly into its architecture as inductive biases, ensuring that biological context shapes the training process itself and yields transparent, reproducible cell- and modality-specific GO importance scores.

While GO (biological process) serves as NetworkVI’s default, pathway collections, data-driven hierarchies, or other DAGs can readily be substituted. (Ribeiro *et al*. 2016, Lipton 2017).

GO activation space decomposition reveals functional immune cell substates, including ICAM1-inflammatory and transitional monocyte subpopulations and four NK tolerance-induction substates, that are invisible to transcriptional clustering or GRNs and emerge in NetworkVI unsupervised. The activation space’s continuous, graded structure makes the framework naturally suited to explore trajectory-like variation. GO importances prioritize programs consistent with immunosenescence and identify perturbation-specific immune activation patterns in the TIL coculture condition where GSEA and ORA find almost no signal, reflecting the advantage of nonlinear aggregation across the GO hierarchy for detecting distributed, low-magnitude transcriptional effects (Frangieh *et al*. 2021).

Because the model is informed by prior knowledge, it is more sensitive to annotation errors, which alter information flow in architecturally consequential ways. Like enrichment analysis, NetworkVI requires careful interpretation: importance scores and attention weights quantify associations rather than causality, and may reflect confounders such as library size or cell cycle state despite batch correction.

The highest GO importances, most crucial in analyses, are most robust across training variants (OR >16x for top 5 % terms), while correlation of all GO importances still reach r=0.41 – r=0.46 under cell subsampling and r=0.68 across training seeds, despite including many low-importance terms that are likely dominated by noise. We recommend caution for sparsely annotated terms, comparisons with fewer than ∼500 cells and experiments with top GO importances barely exceeding 0.5, and recommend treating bootstrap CI width as the primary reliability signal: wide CIs indicate rankings likely to shift under resampling. However, even in low-signal settings like CRISPR perturbations (∼0.50–0.57 AUROC at ∼100 cells per perturbation), meaningful associations remain detectable when CIs are examined. Practical guidance for reliable use of GO and TAD importance as well as attention interpretability is provided in Extended Data Figure 1.

These advantages come at the cost of approximately doubled training time relative to MultiVI (see Methods), a consequence of the structured encoder’s sparse matrix operations. Nonetheless, because the GO encoder scales as O(E) in ontology edges rather than O(G²) over genes, NetworkVI remains tractable at atlas scale.

While we focused mainly on immune benchmark datasets, we observe good generalisation to non-haematopoietic tissues (Supplementary Note 1.3). The current TAD co-regulation layer models only direct, one-step gene co-localisation. Multi-hop propagation, as in GNN message passing approaches (e.g., scNET (Sheinin et al. 2025)), can be beneficial for sparser, more specific interaction sources such as GRNs or TF-target networks, but deeper layers degrade performance for TAD maps (Supplementary Table 1). NetworkVI is inherently complementary to network inference approaches: context-specific inferred subnetworks can serve as inductive biases in its encoder, and the preprocessing pipeline already supports BioGRID, STRING, and TF motif sources. Possible extensions include data-driven ontologies (e.g., CliXO (Kramer *et al*. 2014)) or sparse-subnetwork discovery via lottery-ticket-style pruning (Liu *et al*. 2024), dedicated evaluation in very low-coverage settings such as low-input ATAC-seq, and extension to spatial transcriptomics and methylation modalities.

---

**Extended Data Figure 1:**
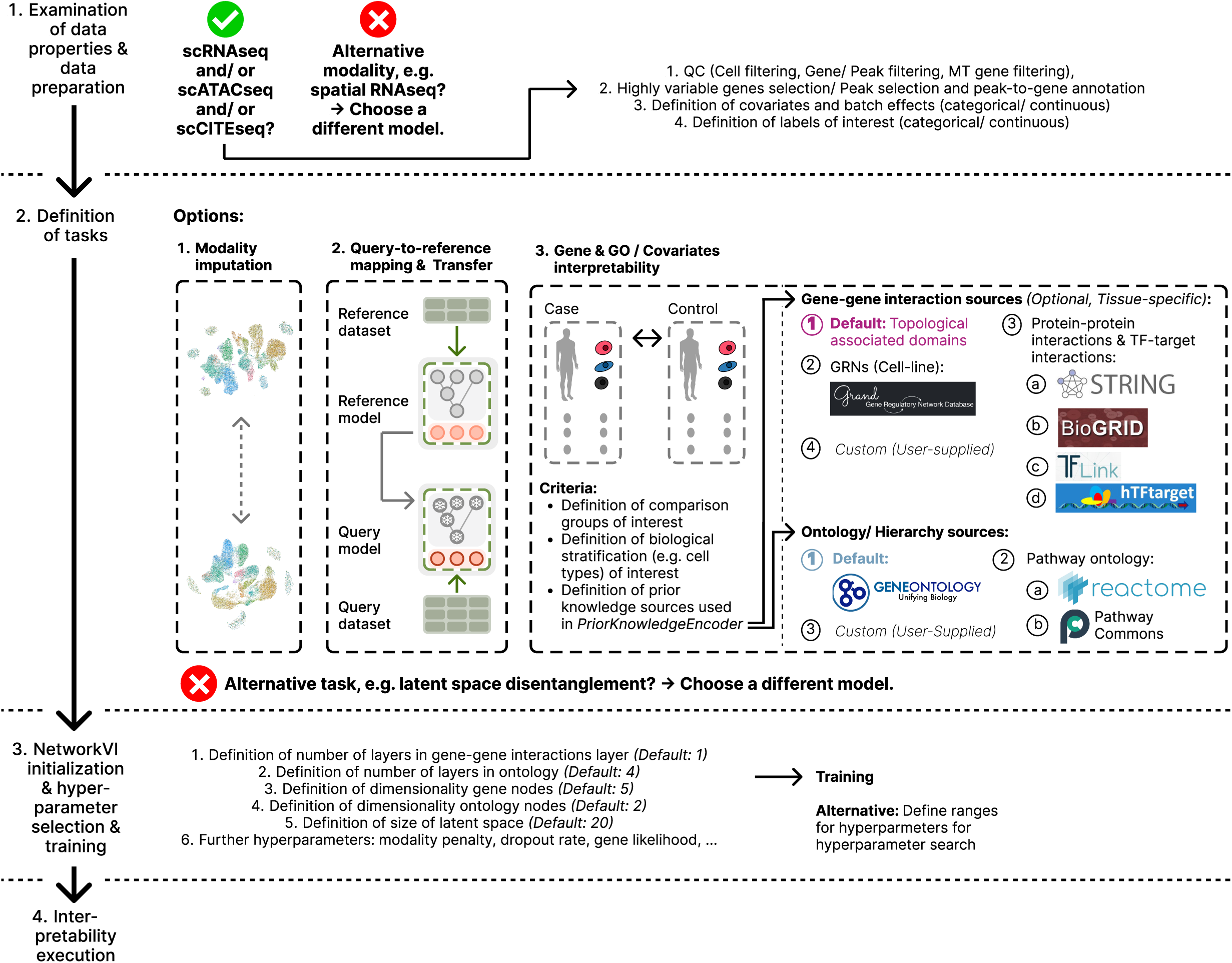
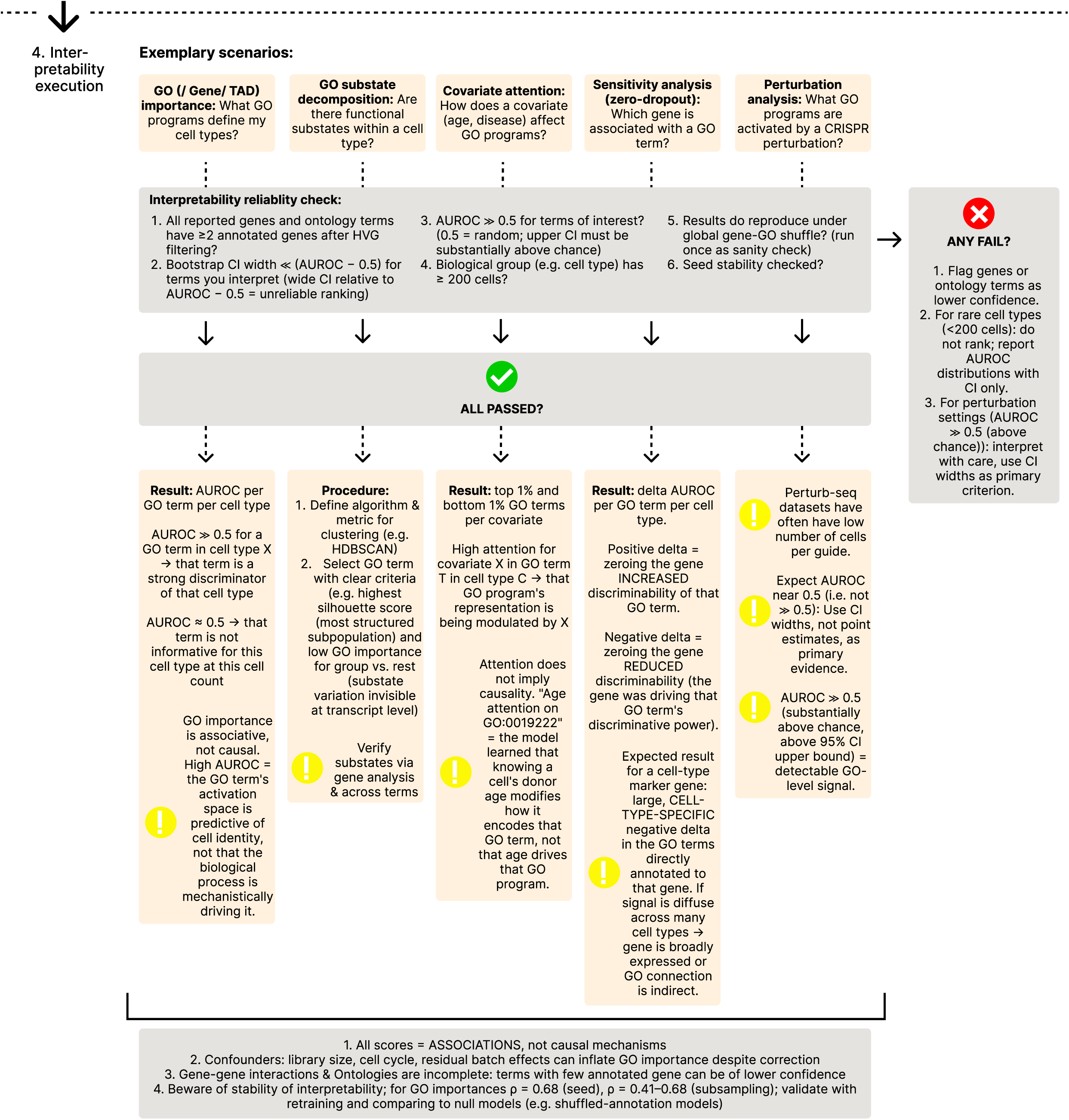
Flowchart for practical use of NetworkVI dataset processing, task definition, training and interpretability.

## Methods

### The NetworkVI architecture

NetworkVI generates a rich, batch-corrected low-dimensional representation of uni-, bi-, and trimodal single-cell datasets, utilizing normalized input data to estimate this representation (Figure 1a). Pairs of decoders and encoders power the integration of each modality. NetworkVI builds upon the multiVI model (Ashuach *et al*. 2023) and thus leverages modality-specific decoders; scVI for transcriptome (Lopez *et al*. 2018), PeakVI for chromatin accessibility (Ashuach *et al*. 2022), and totalVI for epitope data (Gayoso *et al*. 2021). Instead of fully-connected encoders, NetworkVI employs a prior knowledge-driven neural network architecture within the encoder.

### The NetworkVI model

NetworkVI builds on ideas from neural network designs presented by (Ma *et al*. 2018). Specifically, NetworkVI employs a sparse neural network architecture that integrates firstly, a gene layer to aggregate input features when multiple features correspond to the same gene, as is the case with chromatin accessibility data and secondly the GO, which aggregates information from all genes to a single representation per cell. Both genes and GO terms are represented as distinct groups of neural network nodes within the gene layer and GO graph, respectively.

The gene layer integrates co-regulation networks inferred from TADs downloaded from ENCODE (Supplementary Table 2) (Luo *et al*. 2020), which are modeled through a sparse layer with residual connections (Figure 1b). The TAD co-regulation layer is incorporated into the model architecture prior to training. Its inputs are the model inputs at the gene level, and its outputs have the same size as its inputs to enable their use as residuals on the inputs. The input of the lowest GO layer is thus the sum of the gene level inputs, and the residuals from the TAD layer. TAD boundary coordinates are used at model construction time to define the sparse residual connections in the gene layer: for each pair of genes co-localizing within the same TAD, a learnable weighted residual connection is instantiated between their corresponding gene nodes. This sparse connectivity pattern is fixed before any gradient updates occur. During each forward pass, the representation of each gene node is computed as the sum of its own transformed input and the weighted representations of its TAD co-localized partners via these residual connections; only after this gene-level aggregation step does information flow upward into the GO graph. The weights of the residual connections are learned end-to-end during training; the TAD prior therefore determines the *topology* of the gene layer, while the *strength* of each co-regulatory connection is data-driven. The TAD reference used for each dataset was selected to match the predominant lineage of that dataset (Supplementary Table 2); the TAD reference is a user-configurable input and can be replaced with any set of TAD boundary coordinates appropriate for the biological system under study. The TAD reference is supplied as a two-column CSV file in which each row encodes a pair of genes co-localised within the same TAD; any source of TAD calls or gene–gene co-localisation data (e.g. from Hi-C experiments) can be formatted in this way and passed directly to the model. Notably, this also allows the user to combine multiple TAD maps as a union by concatenating files, or use consensus TAD maps if biologically warranted.

The sparse connectivity in the GO graph results from the *is_a* descriptors through which relationships between GO terms are described (Ashburner *et al*. 2000). To connect gene nodes to GO nodes in the GO graph, we use gene-GO annotations which were downloaded from the GO consortium webpage (Release date: 06-17-2024) and filtered to include only GO terms within the biological_process namespace. For hyperparameter optimization, an additional Gene-GO mapping file was curated from the NPInter v5.0 database (Zheng *et al*. 2023). In this process, GO terms associated with ncRNA targets were also assigned directly to the ncRNA genes. By default, NetworkVI filters the GO terms to include only those with a hierarchy level below 5, with each GO term represented by two neural network nodes and each gene represented by five neural network nodes. These values were chosen based on initial tests, where they provided the best performance before performing hyperparameter optimization. Gene nodes are a lot more abundant, and each contain less information than the GO terms, which aggregate multiple gene nodes, hence the much smaller representation. A separate four-layer MLP aggregates all features that cannot be mapped to at least one GO term, including non-coding or poorly annotated genes lacking a recognized Ensembl identifier, as well as ATAC-seq peaks that fall outside the 10,000 bp annotation window and therefore cannot be assigned to any gene, and maps them directly to the root GO term, *biological process* (GO:0008150). This ensures that all input features contribute to the joint latent representation regardless of their annotation status, while keeping the interpretable GO/TAD encoder path restricted to features with valid gene assignments. The number of sparse connections for the transcriptome modality (Supplementary Table 3), chromatin accessibility modality (Supplementary Table 4), and epitope modality (Supplementary Table 5) is given as an example for the DOGMA-seq dataset.

For NetworkVI, a GO term-specific multi-head attention mechanism (Vaswani *et al*. 2017) is implemented to integrate covariates, such as batch effects and the modality (Figure 1c). The attention mechanism uses the covariates as keys *K* and values *V*, and GO term representations as queries *Q*. The original GO embeddings are incorporated through a residual connection. The modified representation is then passed on to the subsequent layers of the architecture.

### Training

NetworkVI can be trained on a dataset *D* with *n* cells with corresponding batch and covariate labels *c* as conditions. As an optional attribute, each dataset can contain cell type labels *l* and each cell may be assigned to a sample *s*_1_, *s*_2_, … , *s_d_*. Furthermore, we consider the dataset to be multimodal (*x*_1_, *x*_2_, … , *x_m_*), with a total of *m* modalities represented. As introduced in MultiVI (Ashuach *et al*. 2023), NetworkVI also models the probability of observing counts of a gene using a negative binomial distribution depending on a learned scaling factor capturing cell-specific biases (Lopez *et al*. 2018). Chromatin accessibility regions are modeled with a Bernoulli distribution (Ashuach *et al*. 2023), and the epitope is modeled with a mixture of negative binomial distributions representing both background and foreground protein expression (Gayoso *et al*. 2021).

Data from each single-cell modality is fed into a modality-specific encoder *e*_1_, *e*_2_, … , *e_m_*. Each encoder firstly includes a co-regulation networks layer, which is a fully connected or sparse layer with dropout, batch normalization, and a user-selectable nonlinear activation function, defaulting to leaky ReLU. Secondly, it includes the (GO-based) dimensionality reduction-module. Finally, the outputs form a normally distributed latent space *p*(*z*|*x*_1_), *p*(*z*|*x*_2_), … , *p*(*z*|*x_m_*) via variational inference from (µ_1, σ_1), … , (µ_*m*, σ_*m*), where µ*_i_*, σ*_i_* ∈ *R^n^*^×ℎ^. Modality-specific latent representations are merged by default via arithmetic averaging, applied separately to both μ and σ with equal weights. Alternatively, a Mixture of Experts (MoE) strategy, using a one-layer gating network with softmax activation, is evaluated during hyperparameter optimization. The gating network generates a weight vector for each modality based on the latent representations, enabling adaptive, dynamic weighting. The merged latent representation is then fed into the decoders. The library size factors l are retrieved via a separate encoder, implemented as a standard fully connected neural network.

### Inference

NetworkVI is trained and inferred using variational inference, similar to standard variational autoencoders (VAEs), but with key modifications, including multiple specialized encoders and decoders, as well as additional loss terms (Kingma and Welling 2014). Then distribution q, parametrized through the parameters η of the encoder networks, factorizes into the posterior distribution of the latent variable z given the inputs x, and the variational distribution of the latent variable l, denoting the library size factor, which is not specific to individual samples:

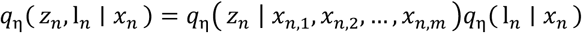

*z_n_* is computed as the mean of the modality-specific latent variables *z_n_*_,1_, *z_n_*_,2_, … , *z_n_*_,*m*_, each modeled as a Gaussian distribution based on the corresponding inputs *x_n_*_,1_, *x_n_*_,2_, … , *x_n_*_,*m*_.

The loss term for the model firstly consists of the ELBO which is made up of the reconstruction log-likelihood E and the Kullback-Leibler (KL) divergence *D_KL_*. Secondly, to align the latent spaces across modalities, an additional loss term - either the maximum mean discrepancy (MMD) loss or the Jeffreys divergence – is added with a scaling factor γ, minimizing the distance between their respective latent representations, as demonstrated in (Ashuach *et al*. 2023).

The loss is computed using variational approximation and is minimized by optimizing both model and variational parameters via stochastic gradient descent:

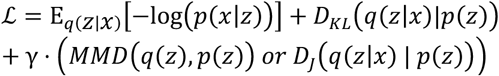

Here, q is the approximate posterior distribution learned by the model and p is the prior distribution, in this case a normal distribution N(0,1).

The MMD loss, which measures the discrepancy between the distributions *q*(*z*) and *p*(*z*), is calculated as defined here:

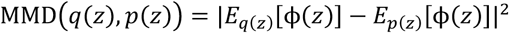

where **ϕ**(*z*) is a feature mapping to a reproducing kernel Hilbert space. Alternatively, the Jeffreys divergence - a symmetric version of the KL divergence, which considers the divergence in both directions between the posterior distribution *q*(*z*|*x*) and the prior distribution *p*(*z*) - is calculated as defined here:

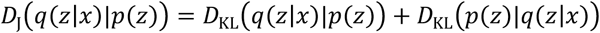

where the KL divergence is defined as:

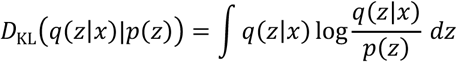

### Downstream tasks

The downstream analyses, including dimensionality reduction, normalization, denoising, and chromatin accessibility estimation, follow the methodologies introduced in MultiVI (Ashuach *et al*. 2023).

For modality-specific and joint latent representations, the mean of the approximate posterior q with the latent *z_m_* as the low-dimensional representation, *x_m_* as the single-cell modality, *s_m_* as covariates, such as batch information, and η representing the set of encoder parameters is returned:

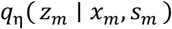

For normalization and denoising of the transcriptome modality, the expected value of the denoised gene expression ρ*_m_* (estimated using the neural network function *f*_Θ_) with l*_m_* as the library size factor (typically equal to 1) is returned as:

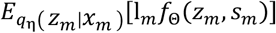

For the estimation of the chromatin accessibility, the expected accessibility *y_m_* is estimated by decoding the accessibility probability *p_r_* (estimated using the PeakVI architecture (Ashuach *et al*. 2022)) under the approximate posterior:

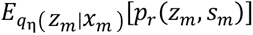

### Training and hyperparameter optimization

The training procedure of NetworkVI has been adapted from MultiVI (Ashuach *et al*. 2023). Briefly, NetworkVI has been trained on 90% of the data, while 10% of the data was used as a validation set, with splits generated at random. By default NetworkVI models have been trained for 500 epochs with early-stopping after at least 50 epochs without improvement of the validation loss using the AdamW optimizer (Loshchilov and Hutter) (η = 0.001 and ε = 0.1), a batch size of 128, and a latent dimension of 20. All experiments were conducted on a high-performance computing cluster. Model training was performed using a single NVIDIA A100 GPU (40 GB memory). Training times ranged from approximately 1 hour on the smallest dataset (DOGMA-seq) to 6 hours on the largest dataset (Perturb-CITE-seq), with early stopping applied. Interpretability analyses were carried out on CPUs, requiring between 2 hours for the smallest dataset and up to 24 hours for the largest dataset, depending on dataset size.

The default hyperparameters of NetworkVI (Supplementary Table 6) were manually determined before performing hyperparameter optimization on all datasets used in this study. Three architectural parameters govern the structure of the prior-knowledge encoder: (i) go_depth, the maximum hierarchy level included from the GO graph; (ii) standard_go_size, the number of neural-network nodes allocated to each GO term; and (iii) standard_gene_size, the number of nodes allocated to each gene in the gene layer. To select default values for these parameters, we performed a sequential optimisation of one parameter at a time on the NeurIPS 2021 CITE-seq BMMC dataset (fully paired), varying each parameter independently while holding the others at candidate defaults. The combined scib score was used as the selection criterion, with results reported in Supplementary Table 7. The larger allocation of gene nodes relative to GO nodes reflects an asymmetry in information density: individual gene nodes each carry sole responsibility for encoding a single input feature, whereas GO nodes aggregate across multiple genes and therefore benefit from richer inputs rather than a larger local hidden dimension. Throughout, we prefer lower parameter counts when performance is equivalent or near-equivalent, as sparser architectures reduce overfitting risk, improve training stability, and keep the model computationally tractable at atlas scale. For all datasets hyperparameter optimization was also performed for MultiVI (Supplementary Table 8), MultiMIL (Supplemntary Table 9) and TotalVI (Supplementary Table 10). The hyperparameter optimization was implemented using the ax library (Bakshy *et al*.) via the Sobol+BoTorch generation strategy (20 runs using the Sobol sampling and 30 runs using the BoTorch model).

### Dataset preprocessing

Standard preprocessing steps have been used to preprocess the transcriptome, chromatin accessibility, and epitope modalities (Heumos *et al*. 2023). For preprocessing of the transcriptome modality, *scanpy* (Wolf *et al*. 2018) was used: genes occurring in less than four cells were filtered out. In addition, cells for which less than 200 expressed genes or more than 20% mitochondrial genes are found were filtered out. For each cell, the counts were normalized to a target sum of 10,000, log-transformed, and scaled. The 4,000 most highly variable genes were determined from the log-transformed data.

For the preprocessing of the chromatin accessibility modality, *epiScanpy* (Danese *et al*. 2021) was used: Peaks occurring in less than 10 cells were filtered. In addition, cells with less than 10 peaks were filtered. For each cell, the counts were normalized to a target sum of 10,000, log-transformed, and scaled. The 20,000 most highly variable peaks were determined from the count data. For the preprocessing of the epitope modality, *muon* (Bredikhin *et al*. 2022) was used to apply centered log ratio normalization.

If the features were not already annotated with Ensembl IDs, they were annotated accordingly. For the annotation of genes in the transcriptome modality, we automated the mapping using Ensembl BioMart (Kinsella *et al*. 2011). Chromatin accessibility peaks were annotated with the nearest Ensembl gene within a maximum distance of 10,000 base pairs. For the genes within the epitope modality a manual annotation was performed.

### Datasets

The DOGMA-seq dataset from (Mimitou *et al*. 2021) includes paired transcriptome, chromatin accessibility, and epitope modalities and was downloaded including cell type annotations from the MultiVI zenodo repository (doi: 10.5281/zenodo.5762077). The original dataset is from GEO (accession no. GSE156478). We use the DIG_CTRL batch with the unperturbed datasets. After preprocessing, the dataset contains a total of 8,900 cells and 210 epitope markers. The CITE-seq dataset from (Hao *et al*. 2021) includes paired transcriptome and epitope modalities and was downloaded including cell type annotations from GEO (accession no. GSE164378). After preprocessing, the dataset comprises a total of 149,058 cells and 224 epitope markers. The NeurIPS 2021 CITE-seq BMMC dataset from (Luecken *et al*. 2021) includes paired transcriptome and epitope modalities and was downloaded including cell type annotations from GEO (accession no. GSE194122). After preprocessing, the dataset comprises a total of 83,290 cells and 134 epitope markers. The NeurIPS 2021 Multiome BMMC dataset from (Luecken *et al*. 2021) includes paired transcriptome and chromatin accessibility modalities and was downloaded including cell type annotations from GEO (accession no. GSE194122). After preprocessing, the dataset comprises a total of 69,110 cells. The Perturb-CITE-seq dataset from (Frangieh *et al*. 2021) includes paired transcriptome and epitope modalities and was downloaded via the zenodo collection (doi:10.5281/zenodo.13350497) of scPerturb (Peidli *et al*. 2024). After preprocessing, the dataset comprises a total of 218,331 cells and 20 epitope markers.

### Dataset limitations and potential biases

All human datasets used in this study are derived from donors of predominantly European ancestry; sex and age distributions are limited and may influence generalizability. While we demonstrate the applicability of NetworkVI across multiple single-cell modalities (transcriptome, chromatin accessibility, and epitope data), differences in annotation methods (manual vs automated) and cell type representation may introduce biases in downstream analyses. Standard batch correction mitigates technical and biological biases.

### Integration metrics

To evaluate the multimodal integration performance, we utilized a set of metrics introduced in the scIB package (Luecken *et al*. 2022). We have used a JAX-based implementation of these metrics provided on https://github.com/YosefLab/scib-metrics. These metrics were initially designed to assess batch correction and biological variance conservation in unimodal integration models. Thus, we assess integration performance using metrics calculated solely on the integrated embedding space. For assessment of the modality mixing, we utilize the metrics as they were defined for batch correction, essentially treating the modality axis as a batch axis. If no batch information is present, the scib score is the sum of 40% of the modality mixing score and 60% of the biological variance conservation score. If batch information is present, the scib score is the sum of 20% of the modality mixing score, 20% of the batch correction score and 60% of the biological variance score. A full breakdown of all individual scib sub-metrics for the NeurIPS 2021 CITE-seq BMMC dataset (fully paired) is provided in Supplementary Figure 6.

#### Batch correction and modality mixing

Batch correction was assessed using the graph connectivity, the integration local inverse simpson index (iLISI) (Korsunsky *et al*. 2019), the k-nearest-neighbor batch-effect test (kBET) (Büttner *et al*. 2019) and the average silhouette width (ASW). The graph connectivity evaluates how well cells of the same type are connected in a k-nearest neighbor graph, with higher scores indicating better batch or modality mixing. The iLISI measures mixing using a graph-based local inverse Simpson’s index, with values scaled from 0 (no mixing) to 1 (perfect mixing). kBET tests local neighborhoods for mixing consistency, where lower rejection rates signal effective correction. The ASW quantifies clustering quality by comparing the average intra-cluster distances to the average inter-cluster distances.

#### Biological variance conservation

Biological variance conservation was assessed using the cell type local inverse simpson index (cLISI) (Korsunsky *et al*. 2019), the adjusted rand index (ARI), the normalized mutual information (NMI), and the isolated label average silhouette width (ASW). The cLISI measures cell type clustering using a graph-based local inverse Simpson’s index, with values scaled from 0 (mixing) to 1 (no mixing). ARI and NMI compare integrated clustering with the ground truth labels. In contrast to the original implementation of scIB (Luecken *et al*. 2022), where k for k-means is inferred, the JAX-based metric implementation does not infer k but instead sets it to the given number of cell types. Isolated label ASW evaluates cluster compactness, distinguishing other cell types.

### Query-to-reference mapping

To evaluate the query-to-reference mapping performance, we adapt the scArches approach introduced by (Lotfollahi *et al*. 2022). This approach, termed architectural surgery, involves training a model on a reference dataset and finetuning it on a query dataset (Figure 1d). In the MultiVI model, batch and modality information are concatenated with the data before being fed into the first layers of both the encoder and decoder. During finetuning, all layers without additional covariate information are frozen. While it is generally desirable to freeze layers as this better utilizes the pretraining, layers with covariate inputs cannot be frozen, so that they can learn the dependencies of the new covariate distribution. In NetworkVI, only the GO term-specific covariate embeddings (roughly 1.3% of the trainable parameters in the encoder) and the first layer of the decoder are finetuned, while all other weights remain frozen. In contrast to MultiVI, this approach enables NetworkVI to capture covariate information across all layers of the encoder, which results in an increased number of parameters that require optimization.

For label transfer, a simple logistic regression classifier trained on the generated latent space was used; however, as with other latent space-based methods, NetworkVI is compatible with a range of classifiers. The query-to-reference mapping performance is assessed for three query-to-reference mapping scenarios: Training of NetworkVI and MultiVI on three batches (sites) out of four of the (a) CITE-seq BMMC dataset and the (b) Multiome BMMC dataset (site2, site3, and site4). Query mapping is then performed for the missing batch (site1) with either both modalities available, or unimodally after the removal of one of the modalities by setting all features to zero. For a cross-dataset and cross-tissue label transfer performance examination we (c) train on the full CITE-seq BMMC dataset and use the CITE-seq dataset generated by (Hao *et al*. 2021) as the query.

The micro F1 score is used to evaluate the accuracy of transferring cell type labels between two datasets. While optimizing the classifier choice and tuning hyperparameters could enhance performance, the experiments are run with the default settings of MultiVI, NetworkVI, and the logistic regression classifier as this is the more likely choice for users of NetworkVI. We calculate 95% confidence intervals via bootstrapping with 200 samples. The same bootstrapping procedure with 200 samples is used to compute 95% confidence intervals for all AUROC-based GO importance, gene importance, and TAD importance scores reported throughout the manuscript, as described in detail in the Gene and GO interpretability section.

### Transcriptome imputation

We compare the imputed transcriptome data with the raw count data and a denoised, smoothed version of the count data, as implemented in MultiVI (Ashuach *et al*. 2023). This smoothing is necessary due to the stochastic nature of single cell transcriptome sequencing, in which the observed transcript count for a single cell can deviate substantially around the true count of the cell population, especially for low-expression genes. When imputing the transcriptome, the estimate is closer to the true count, which in turn is more comparable to the smoothed data. Briefly, the top 30 principal components were used to construct a k-nearest neighbors (KNN) graph (k = 10), followed by averaging the expression values of neighboring cells.

### Benchmarking

For fair benchmarking, we selected the methods based on their reported integration performance in the respective publications and the benchmark results from (Hu *et al*. 2024). We configured all methods to generate embeddings with 20 latent dimensions to ensure consistency across paired and mosaic integration tasks. All methods were run once with their default hyperparameters for paired and mosaic integration. Additionally, for all datasets, we performed hyperparameter optimization as previously described for NetworkVI, MultiVI, MultiMIL, MOFA+, and TotalVI. We did not conduct hyperparameter optimization for MIDAS, as its training time exceeded that of the other benchmarked methods by up to an order of magnitude and was thus impractical, and likewise skipped optimization for WNN, as it lacks commonly tuned hyperparameters in practice.

For paired integration we benchmarked NetworkVI against WNN (Hao *et al*. 2021), MOFA+ (Argelaguet *et al*. 2020), MultiVI (Ashuach *et al*. 2023), MultiMIL (Litinetskaya *et al*. 2024) and MIDAS (He *et al*. 2024) on the DOGMA-seq dataset, the CITE-seq dataset by (Hao *et al*. 2021) and the BMMC datasets. For the CITE-seq datasets, we also benchmarked against TotalVI (Gayoso *et al*. 2021). For the DOGMA-seq dataset, we utilized the extension of WNN to an arbitrary number of modalities as presented by (Svensson *et al*. 2020) and implemented in the *muon* library (Bredikhin *et al*. 2022). The weighted-nearest neighbor (WNN) algorithm calculates cell-specific modality weights by identifying nearest neighbors, predicting affinities across modalities, and applying a softmax transformation, with modality weights summing to 1 per cell (Hao *et al*. 2021). MOFA+ is a linear factor model that is trained using variational Bayes to optimize the ELBO, decomposing the input data into latent factors and factor effects. MultiVI, MultiMIL, MIDAS, and TotalVI are VAE-based methods. MultiVI and TotalVI are trained on count data and thus fit modality-specific distributions. TotalVI employs a single encoder-decoder for CITE-seq data, while MultiVI aligns the modality-specific latent representations, which are generated via separate encoders, via a modality penalty. MultiMIL and MIDAS utilize the product of experts framework to generate a joint low-dimensional embedding (Lee and van der Schaar 2021).

For the mosaic integration task, we also benchmarked NetworkVI against the already introduced MultiVI, MultiMIL and MIDAS on the aforementioned datasets by artificially unpairing data as needed.

### Computational scalability

To assess the computational scalability of NetworkVI to atlas-scale datasets, we benchmarked training and interpretability runtimes on the Human Lung Cell Atlas (Sikkema *et al*. 2023), one of the largest publicly available human single-cell atlases, comprising 2,282,447 cells. Following the same preprocessing applied across all other datasets, genes were filtered to the 4,000 most highly variable. We trained NetworkVI on randomly subsampled fractions of the full HLCA (20%, 40%, 60%, 80%, 100%), corresponding to approximately 456,000 to 2,282,000 cells.

Training was performed on a single NVIDIA A100 GPU (40 GB). NetworkVI approximately doubles training time relative to MultiVI (∼3 hours vs. ∼1.5 hours on the NeurIPS 2021 CITE-seq BMMC dataset), a consequence of the structured encoder relying on slower sparse matrix operations. GPU memory allocation remained stable across all subsets at approximately 761 MB (reserved: ∼990 MB), reflecting that NetworkVI employs mini-batch training and GPU memory consumption is therefore independent of dataset size. Wall-clock training time scaled sublinearly with dataset size (from 2.5 hours at 20% to 10.4 hours at 100% of the full atlas) because early stopping was applied and larger datasets tend to converge in fewer epochs (56–68 epochs across subsets), with each epoch requiring proportionally more time. CPU memory use scaled approximately linearly with cell count, ranging from ∼36 GB at 20% to ∼250 GB at 100%. Because the sparse GO encoder scales as O(E) in ontology edges rather than O(G²) over genes, this computational overhead does not compound at scale.

GO importance analysis, which runs on CPU scales approximately linearly with the number of cells included in the comparison. For illustration, at the 20% subset the benchmark comparison comprised 22,696 CD8-positive alpha-beta T cells versus 16,562 CD4-positive alpha-beta T cells across 1,306 modeled GO terms, completing in 2.6 hours. This yields an approximate per-GO-term runtime of ∼7 seconds per GO term at this cell count, which users can scale according to the number of GO terms and cell counts in their own datasets. Covariate attention analysis requires a single forward pass through the model across all cells and is therefore bounded by inference time rather than the interpretability computation itself. Taken together, NetworkVI is usable on current research-grade hardware for typical existing datasets up to atlas-scale of millions of cells.

### Importance of the gene layer and the GO hierarchy

To assess the impact of the gene layer, the co-regulation prior information, and the GO layer, we performed ablation studies on the fully paired NeurIPS 2021 CITE-seq dataset. We tested the effect of nonsensical TAD information by replacing the TAD co-regulation networks prior knowledge with TAD, that have been shifted by the mean length of all TAD in this cell line. Second, we made the TAD information nonspecific by replacing the TAD co-regulation networks prior knowledge retrieved from the K562 cell line with TAD co-regulation networks prior knowledge from alternative cell lines (GM12878, MCF-7, T47D), STRING database (Szklarczyk *et al*. 2023) and the BIOGRID database (Oughtred *et al*. 2021), GRAND GRN (Ben Guebila *et al*. 2022), TFLink (Liska *et al*. 2022), hTFtarget (Zhang *et al*. 2020)). As an alternative to the GO biological process namespace, we evaluate the molecular function and cellular component namespaces. Additionally, we tested former releases of GO (2024-06-01, 2019-06-09, 2022-06-15) and pathway ontologies, namely Reactome (Milacic *et al*. 2024) and Pathway Commons (Cerami *et al*. 2011). For all reported percentage changes in GO importance between the full and ablated models, 95% confidence intervals were computed by bootstrapping the AUROC difference across cells, as described in the Gene and GO interpretability section.

### Gene and GO interpretability

The prior knowledge-driven architecture of NetworkVI allows for the inference of a metric representing the importance of specific components within an encoder, such as individual GO terms or genes, for the nonlinear model performance (see Supplementary Figure 14). For groups of nodes representing individual genes or GO terms, logistic regression analyses are conducted using the activations, as previously illustrated by (Ma *et al*. 2018) (see Figure 2c). The case labels can be defined based on patient- or cell-level phenotypes, for instance a particular cell type. Control labels may then be all alternative labels or a manual selection of specific subgroups. The importance of each gene and GO term is quantified by the area under the receiver operating characteristic curve (AUROC) from the logistic regression models fitted against a specific label, e.g. the cell type. Thus, in the absence of any true effects, we would expect the GO importance to be close to 0.5. In a multimodal setting, the GO importances are computed independently for each modality. This measure can also be employed for downstream analysis, such as assessing the similarities between distinct cell types, the effect of *in vitro* perturbations or specific disease cohorts. TAD importances can be computed analogously by concatenating the activations of the genes in a TAD and computing the AUROC in the same fashion. Notably, this is possible both with and without the TAD layer, because it only relies on gene-level activations and the Gene-to-TAD assignment. If the TAD-layer is present, gene representations after the TAD layer are used.

While GO importance allows for global comparisons of functional relevance across modalities and cell types, it also supports fine-grained interpretability at the level of individual genes. To this end, we leveraged the model’s structure to perform an *in silico* investigation of counterfactual scenarios. Specifically, cell type-specific marker genes are sequentially removed from a dataset by treating all values for these marker genes as dropouts and setting them to zero (see Supplementary Figure 14). The difference in GO term importance between the modified dataset (with removed marker genes) and the original dataset (with marker genes in place) is used to quantify the association of each marker gene with the respective GO terms. This approach is analogous to the image degradation task commonly used for quantitative evaluations of explainable models for images, where instead of patches we ‘degrade’ genes by setting their expression to zero (Schulz et al. 2020). In comparison to conventional gene-GO annotations, our approach provides a quantifiable, cell type-specific resolution of these associations. To analyze the transcriptional correlates of specific GO terms across individual cells, we projected the multi-dimensional activation space of a given GO term onto a one-dimensional axis using principal component analysis. Specifically, we fitted a one-dimensional PCA on the activations of all nodes corresponding to the GO term of interest across cells. The first principal component, capturing the dominant direction of variation, was then used to rank cells by their GO activation profile. This ranked activation space enables correlation analysis with gene expression levels and stratification of cells into quantiles for downstream transcriptional profiling. To assess the robustness of GO importance scores to training stochasticity and dataset size, NetworkVI was retrained under three variant conditions on the NeurIPS 2021 CITE-seq BMMC dataset: an alternative random seed, and two cell-subsampled variants retaining 75% and 50% of cells, respectively. GO importance AUROC scores were computed identically in all variants. Spearman rank correlations between per-GO-term AUROC importance scores of the default model and each variant were computed for the Naive CD20⁺ B IGKC⁺ cell-type comparison task. To verify the robustness of GO importance scores to training stochasticity and dataset size, we computed Spearman rank correlations of per-GO-term AUROC values for Naive CD20⁺ B IGKC⁺ cells between the full model and three variant conditions: an alternative random seed (ρ = 0.681, p = 1.98×10⁻²³), subsampling to 75% of cells (ρ = 0.456, p = 1.62×10⁻²²), and subsampling to 50% of cells (ρ = 0.413, p = 1.31×10⁻¹⁶). Seed stability exceeds subsampling stability, as expected: reseeding perturbs only weight initialisation and minibatch order, whereas reducing cell count also reduces the statistical power of the underlying logistic regression. These correlations are computed across all modelled GO terms; for GO terms with high importance in a given comparison, for the biologically relevant subset rank concordance may be higher. To quantify top-term membership stability rigorously and across all available cell types, we additionally computed an odds-ratio (OR) enrichment test for each model comparison. At each percentile threshold q, GO terms were independently classified as ‘top’ (AUROC ≥ q-th quantile of that model) or ‘bottom’, and a one-sided Fisher exact test assessed whether the top-tier sets identified by both models overlap more than expected by chance. This was conducted across 29 shared cell types for the 50%/75% subsampling comparison and 7 shared cell types for a seed 1 vs. seed 2 comparison at 100% of cells. Across all cell types for the 50%/75% subsampling comparison, the geometric mean OR was 16.0× at the top-5% threshold (∼90 GO terms), 9.6× at top 10%, and 6.2× at top 25%. An independent seed comparison on the full dataset yielded nearly identical enrichments (OR = 16.6× at top 5%, 9.5× at top 10%), confirming that top-tier identity is driven by biological signal rather than initialisation-specific convergence. While Spearman correlation within the top-percentile subgroup approaches zero, reflecting shifts in relative ordering among already-important terms, the composition of the top tier is highly reproducible across training variants.

To quantify uncertainty in GO importance and gene importance scores, we computed 95% bootstrap confidence intervals for all reported AUROC values. Specifically, for each GO term or gene, we drew 200 bootstrap samples with replacement from the set of cells used for logistic regression evaluation, requiring that both classes were represented in each sample. The AUROC was recomputed for each bootstrap replicate, and the 2.5th and 97.5th percentiles of the resulting distribution were taken as the lower and upper bounds of the 95% confidence interval, reported around the median. Bootstrap samples in which only one class was represented were discarded. This procedure was applied uniformly to all AUROC values reported in the main text and figures, including GO importance scores for individual cell types, substate-level GO importance scores, and TAD-level importance scores. TAD importance confidence intervals were computed analogously by concatenating gene-level activations within each TAD prior to bootstrapping. For percentage changes in GO or TAD importance between model variants (e.g., full model versus ablated model), confidence intervals were propagated by bootstrapping both models independently and computing the distribution of pairwise differences. To assess differences in predictive performance across groups, we computed bootstrapped AUROC distributions (n=200 resamples with replacement) for each group. Group-wise AUROCs are reported as median with 95% confidence intervals derived from the 2.5th and 97.5th percentiles of the bootstrap distribution. Differences in AUROC between groups were assessed using the two-sided DeLong test (DeLong *et al*. 1988), which accounts for the correlation structure of AUC estimates derived from overlapping samples. A significance threshold of α=0.05 was applied with Bonferroni correction for three pairwise comparisons. Spearman correlation coefficients reported in the text were accompanied by 95% confidence intervals computed via bootstrap resampling (10,000 iterations) by resampling the unit of observation with replacement: TADs for correlations between TAD-level quantities (TAD importance delta versus TAD size, Hi-C contact domain score, HVG normalized dispersion, and variance of TAD importance improvement), and cells for within-population gene expression correlations (e.g. IRF8 and BCL11A versus pDC activation space scores), for which 95% CIs were computed via Fisher’s z-transformation. Statistical significance of GO and TAD importance deltas calculated comparing the two model variants was assessed using two-sided Wilcoxon signed-rank tests across phenotypes, with 95% bootstrap confidence intervals computed by resampling phenotypes with replacement (10,000 iterations). To assess whether TAD importance deltas are associated with CpG island density, we computed the number of UCSC hg38 CpG islands (downloaded from the UCSC Table Browser, *cpgIslandExt* track) overlapping each TAD genomic locus, normalized by TAD span in kilobases. TAD genomic coordinates were derived by taking the union of Ensembl gene body coordinates (GRCh38) for all constituent genes, retrieved via BioMart. Overlap with CpG islands was computed using bedtools intersect (v2.30). The resulting CpG island density was correlated with mean TAD importance delta using Spearman correlation with 10,000-iteration bootstrap confidence intervals.

To assess whether TAD importance scores align with independent genomic regulatory signals, we computed Spearman correlations between TAD importance scores and ENCODE K562 ChIP-seq signals for 458 transcription factors and chromatin regulators from multiple laboratories (Full list of files in Supplementary Table 11), accessed as p-value signal bigWig tracks from the ENCODE portal (https://www.encodeproject.org). For each genomic marker, the mean bigWig signal was extracted across the TAD genomic locus using pyBigWig, and Spearman correlation was computed between this signal vector (across all TADs) and the TAD importance score vector, separately for each of the 44 cell types and for both the full model and the ablated baseline without the TAD co-regulation layer. To identify regulators showing the largest improvement in alignment between the full model and baseline, markers were ranked by mean change in Spearman correlation across cell types (model minus baseline). Statistical comparison between the full distributions of model and baseline correlations was performed using the Wilcoxon signed-rank test and the two-sample Kolmogorov-Smirnov test. All analyses used the expression modality encoder.

### Position-shifted TAD negative control

To generate a negative control for the TAD co-regulation layer that preserves overall genomic statistics while destroying biologically meaningful co-regulatory groupings, TAD boundary coordinates were shifted along the chromosome by a fixed offset equal to the mean TAD span across all modelled TADs in the K562 reference (ENCFF173VDJ). Genes were then re-assigned to shifted TAD groups using the same proximity-based procedure applied to the original boundaries. This procedure preserves the number of TADs, the distribution of TAD sizes, and the overall density of residual connections in the gene layer, while ensuring that co-localised gene pairs within the original TADs are no longer grouped together. The position-shifted model was trained and evaluated identically to the full K562 model.

### Global shuffle negative control

To assess whether GO importance scores reflect biologically correct gene–GO associations or architectural properties of the sparse encoder, we generated globally shuffled gene–GO annotation files. For each of the three human GAF annotation files (canonical, isoform, RNA), the GO term assignments were permuted uniformly at random across all annotated gene–GO pairs using a fixed random seed (seed = 42), preserving the total number of annotated pairs and the marginal distribution of GO term usage frequencies while destroying all biologically meaningful gene–GO associations. A NetworkVI model was trained and evaluated identically to the default model using the shuffled annotations in place of the true GO Biological Process annotations.

### K562 interaction shuffle negative control

To test whether the performance benefit of the TAD co-regulation layer requires biologically meaningful gene–gene co-localisation (as encoded in the K562 Hi-C TAD calls) or arises from architectural complexity alone, we generated a shuffled interaction control in which the K562 gene– gene interaction pairs (ENCFF173VDJ) were randomly permuted while preserving the number of connections per gene.

### Spike-in positive-control analysis

To evaluate whether NetworkVI’s gene–GO routing is biologically specific, we performed spike-in perturbation experiments on the NeurIPS 2021 CITE-seq BMMC dataset. For each of 17 biologically characterized genes spanning 6 functional groups, expression in all cells of a designated target cell type was multiplied by 5× in the raw count layer prior to inference, while all other cell types were unchanged. After encoding with the trained model, cell-type discriminative AUROCs were computed for each of the 1,791 GO Biological Process latent dimensions using one-vs-rest logistic regression. ΔAUROC was defined as the spiked AUROC minus the unperturbed baseline AUROC for each GO term. GO annotations for the spiked gene were propagated through the GO hierarchy using the true-path rule (each direct annotation implies all ancestor terms), following standard GO enrichment analysis conventions and correctly mapping gene-level annotations to the hierarchical level represented in the model. Enrichment of annotated GO terms in the distribution of |ΔAUROC| values was assessed independently for each experiment using a one-sided Mann-Whitney U test (annotated vs. all 1,791 GO terms). All 17 experiments reached statistical significance (p < 0.05; 16 of 17 at p < 0.001), spanning baseline expression levels from 0.0% to 49.0% of cells in the spiked cell type. Representative activated GO terms for selected genes are shown in Supplementary Figure 27.

### Differential GO term activity across assay conditions

To quantify the number of GO terms with statistically significant differential activity across the three experimental conditions (−IFNγ vs. control, +IFNγ vs. control, TIL coculture vs. control), we applied pairwise two-sample DeLong tests between all condition pairs for each GO term passing the pre-defined parent-term selection criteria. Because the three screens comprise non-overlapping sets of cells, the cross-covariance term of the standard DeLong variance estimator is zero, and the variance of the difference in AUROCs reduces to the sum of the individual variances: Var(AUC_i − AUC_j) = Var(AUC_i) + Var(AUC_j). Each per-condition variance was estimated from the stored 95% bootstrap CI half-width as SE = CI_halfwidth / 1.96, giving Var(AUC) = SE². A GO term was classified as differentially active if at least one of the three pairwise comparisons was significant after Bonferroni correction for three tests (corrected α = 0.05/3 ≈ 0.017, corresponding to |z| > 2.39, where z = (AUC_i − AUC_j) / √(SE_i² + SE_j²)). The total number of differentially active GO terms per perturbation gene was plotted as a bar chart (Figure 6d).

### GO-based substate identification

To identify GO terms with structured within-cell-type variation for a given cell type, all modeled GO terms were screened by applying HDBSCAN (minimum cluster size = 10) to the first two principal components of the multi-dimensional GO term activation vector for cells of that type. The silhouette score was computed for each GO term as a measure of cluster separation. GO terms were ranked separately by their GO importance score (AUROC for the given cell type versus all others) and their silhouette score. The GO term selected for detailed substate analysis was the one ranking highest on both criteria simultaneously. Substate labels were assigned based on differential expression between HDBSCAN-identified clusters: genes were retained if mean expression exceeded 0.1 in at least one cluster, and ranked by log2 fold change. The top-ranked gene in the pairwise comparison between the most transcriptionally active cluster and the quiescent cluster was used as the primary label for that substate, with additional co-expressed markers listed in the text. Substate robustness was confirmed by examining donor and batch distributions within each substate. As a baseline we implemented a standard workflow comprising Leiden clustering (resolutions 0.3–3.0), ORA/GSEA, GRNBoost2 (Moerman *et al*. 2019), and SCENIC (AUCell) (Van de Sande *et al*. 2020). GRN targets were aggregated for top-ranked transcription factors and subjected to GO enrichment. SCENIC regulon activity matrices were clustered using HDBSCAN. Agreement with NetworkVI substates was quantified using adjusted Rand index (ARI), and cluster separability using silhouette scores. GO recovery was assessed by overlap with ORA, GSEA, GRN-based, or SCENIC-derived enrichments. Analyses were performed per cell type; missing outputs were treated as non-detections.

To model *in vitro* perturbations, NetworkVI was trained on the unperturbed control dataset, and inference was run for both the perturbed and unperturbed data. GO importance scores for a specific cell type were then computed by assigning the perturbation status of cells as labels for each perturbed gene individually. The GO importances thus reflect how relevant each term is in distinguishing between perturbed and unperturbed cells based on representations learned on the unperturbed cells. As a baseline, we calculated GO enrichment using gene set enrichment analysis (GSEA), as implemented in the GSEApy package (Fang *et al*. 2023), following the method described by (Barbie *et al*. 2009). As an additional baseline, we calculated GO over-representation analysis (ORA) using the Enrichr interface implemented in the GSEApy package, applied to differentially expressed genes between CD58-perturbed and unperturbed cells within each screen condition, identified using Scanpy’s Wilcoxon rank-sum test (Benjamini-Hochberg FDR < 0.05, |log2FC| > 0.5).

### Covariate attention uncertainty quantification

To quantify uncertainty in covariate attention values, we computed 95% confidence intervals using a normal approximation. For each GO term or gene node, attention weights were collected across all cells assigned to the relevant label (e.g., cell type). The per-node attention matrix has shape (cells x outputs x covariates). This matrix was first flattened across the cells and outputs dimensions to yield a sample of attention values for each covariate. The mean attention and its standard error were then computed across this flattened sample, and the 95% confidence interval was constructed as mean ± z₀.₀₂₅ × SE, where z₀.₀₂₅ = 1.96. Group-level attention values, aggregating across covariate categories (e.g., all levels of a categorical covariate), were computed as the mean of the individual covariate attention values within the group, with confidence intervals propagated accordingly as the mean of the per-covariate lower and upper bounds. This procedure is applied uniformly to all attention values reported in the main text and figures, including cell-type-level and GO-term-level attention values for age, smoking status, BMI, and collection site covariates.

### Software Implementation and Dependencies

NetworkVI was developed for Python versions 3.9, 3.10 and 3.11, and implemented using the following key packages: flax, lightning (≥2.0), ml-collections (≥0.1.1), mudata (≥0.1.2), numpyro (≥0.12.1), rich (≥12.0.0), scikit-learn (≥0.21.2), sparse (≥0.14.0), torchmetrics (≥0.11.0), xarray (≥2023.2.0), pqdm, prettytable, goatools, tabulate, addict, and torch (version 2.2.0).

## Data availability

All datasets analyzed in this manuscript are public and can be downloaded through https://github.com/LArnoldt/NetworkVI_reproducibility. The DOGMA-seq dataset is available from GEO (GSE156478). We used the DOGMA-seq dataset as provided in the MultiVI zenodo (doi: 10.5281/zenodo.5762077). The NeurIPS 2021 CITE-seq and Multiome BMMC dataset is available from GEO (GSE194122). The CITE-seq dataset from (Hao *et al*. 2021) is available from GEO (GSE164378). The Perturb-CITE-seq dataset is available via zenodo (doi:10.5281/zenodo.13350497). The Human Lung Cell Atlas dataset is available via CELLXGENE (https://cellxgene.cziscience.com/e/9f222629-9e39-47d0-b83f-e08d610c7479.cxg/).

## Code availability

The code for NetworkVI is publicly available at https://github.com/LArnoldt/NetworkVI (Arnoldt *et al*. 2026). NetworkVI’s open-source implementation within the scvi-tools ecosystem ensures broad accessibility and reproducibility. Preprocessing scripts, the code to reproduce the results and the custom notebooks needed to generate the figures in this paper will be made available upon publication.

## Acknowledgements

We would like to acknowledge Carl Herrmann, Simon Anders, Leonhard Kohleick, Tillmann Rheude, Johannes Kraume and Vincent von Lipinski for helpful discussions and feedback. We thank the scverse community, in particular the developers and the maintainers of scvi-tools. We thank the participants of the studies used in this work for providing their data. L.A. was supported by the Helmholtz Association, as part of the joint research school Munich School for Data Science (MUDS). The study has been supported by the Deutsche Forschungsgemeinschaft (DFG, German Research Foundation) – Project-ID 437531118 – SFB 1470. The authors acknowledge the Scientific Computing of the IT Division at the Charité - Universitätsmedizin Berlin for providing computational resources that have contributed to the research results reported in this paper.

## Contributions

L.A., J.U., B.W., and R.E. conceived and designed the project. L.A., J.U., L.H. implemented models, conducted experiments, and performed data analysis. L.A., J.U., L.H. and N.I. conceptualized the gene-to-TAD strategy. K.N. supported the analysis. F.J.T. provided methodological support and contributed to the discussion of the results. L.A., J.U., N.I., B.W., and R.E. wrote and prepared the manuscript. All authors read, revised, and approved the manuscript.

## Ethics declarations

F.J.T. consults for Immunai Inc., Singularity Bio B.V., CytoReason Ltd, Cellarity, and has ownership interest in Dermagnostix GmbH and Cellarity.

