## Supplementary Information for "Biologically Informed Variational Inference Enables Interpretable Cell Phenotyping and Discovery"

### Supplements

#### Supplementary Note 1: Integration, query-to-reference mapping, and TAD co-regulation details

##### 1.1 Integration and query-to-reference mapping with NetworkVI

We evaluated integration quality on a trimodal DOGMA-seq dataset [1] and three additional bimodal datasets spanning diverse tissue types: the NeurIPS 2021 Multiome dataset and the CITE-seq BMMC dataset [2], and a CITE-seq PBMC dataset [3]. We benchmarked performance in both paired and mosaic integration scenarios. For mosaic settings, we simulated unpaired data by randomly omitting cells from each modality at controlled paired rates. In addition to NetworkVI, we included a range of state-of-the-art baselines representing diverse modelling paradigms. For both paired and mosaic settings, we evaluated MultiVI [4], MultiMIL [5], and MIDAS [6]. For paired integration only, we also considered totalVI [7], WNN [3, 8], and MOFA+ [9]. For each dataset and method, we performed independent hyperparameter optimization for all tasks. MIDAS was computationally prohibitive, requiring training times up to an order of magnitude longer than other methods, while WNN has no commonly tuned hyperparameters in practice.

On the trimodal DOGMA-seq dataset, NetworkVI achieved the highest scib scores in both paired and mosaic integration, outperforming other methods (Supplementary Figure 2a, 3). Its performance remained robust across paired rates, driven by strong preservation of biological variance. While MIDAS excelled at low paired rates (e.g., 20%), it did so by emphasizing modality mixing over biological fidelity. On bimodal datasets, NetworkVI consistently achieved state-of-the-art performance. For the Multiome BMMC dataset, it outperformed all methods on fully paired data, with MultiVI performing comparably at lower paired rates (Supplementary Figure 2b, 4). For CITE-seq datasets, MultiVI remained competitive, especially on the BMMC dataset, while NetworkVI showed superior or comparable performance across paired and unpaired settings (Figure 2c, Supplementary Figure 5–7). WNN showed strong biological conservation, particularly on the CITE-seq PBMC dataset [3], for which it was originally developed; however, its lack of batch correction results in poor batch correction subscores and thus lower overall scib scores. Overall, NetworkVI offered the best preservation of biological signal across integration scenarios.

In query-to-reference mapping, a query dataset is contextualized with a reference dataset, enabling downstream tasks such as cell-level label transfer. This allows users to utilize the resource-intensive annotation of previous datasets for new datasets, resulting in more comparable annotations. To this end, we adapt scArches [10] (see Methods), training MultiVI and NetworkVI in a cross-tissue query-to-reference mapping scenario on the entire CITE-seq BMMC dataset and using the CITE-seq dataset from [3] as the query. Cell labels are assigned to the query dataset by training a separate classifier on the resulting latent spaces. Before mapping, we harmonized cell type annotations between the two datasets, though some cell types were unique to each dataset and could not be mapped. NetworkVI improves the micro F1 score by 0.141 (0.871 vs. 0.730) for the bimodal query and by 0.189 (0.717 vs. 0.528) for the unimodal query with the transcriptome modality (Supplementary Figure 2d). Like MultiVI, NetworkVI accurately assigns most cells to their correct or closely related cell types. Incorrect mappings occurred more frequently for unimodal queries of the transcriptome and epitome modalities (Supplementary Figure 8a–c vs. 8d–f).

We additionally evaluated NetworkVI in two cross-batch mapping scenarios, where the method was tasked with inferring cell labels for one batch, using training data from other batches. NetworkVI (Supplementary Figure 9a) showed an average absolute improvement in the F1 score (micro) of 0.039 (0.770 vs. 0.731) and 0.077 (0.721 vs. 0.644) compared to MultiVI (Supplementary Figure 9b) on the CITE-seq BMMC dataset for bimodal and unimodal queries, respectively (Supplementary Figure 2d). NetworkVI and MultiVI reach the same F1 score for the cross-batch mapping scenario on the Multiome BMMC dataset (Supplementary Figure 2d, Supplementary Figure 10).

Imputing gene expression on the CITE-seq BMMC dataset with NetworkVI results in a mean per-gene Spearman correlation of $0.157$, where $\Delta r_{\text{NetworkVI}-\text{MultiVI}} = 0.08$ (Supplementary Figure 11a–f). Using denoised unnormalized gene expression (see Methods), the correlation increases to 0.313, where NetworkVI and MultiVI perform approximately equally. NetworkVI imputes more genes with Spearman correlation >0.8 compared to MultiVI (81 vs. 77 genes; Supplementary Figure 11g–h), a 5% increase that holds regardless of whether denoised or raw gene expression data is used, with the additionally recovered genes involved in immune system function, cell cycle regulation, metabolism, and transcriptional control. Furthermore, on the Multiome BMMC dataset, NetworkVI improves on MultiVI's ATAC imputation performance, reaching an average precision of $0.264$ with $\Delta r_{\text{NetworkVI}-\text{MultiVI}} = 0.05$ (Supplementary Figure 12).

##### 1.2 Prior knowledge source comparison

NetworkVI was compared across a comprehensive range of alternative prior knowledge sources on the NeurIPS 2021 CITE-seq BMMC dataset, with performance measured as ΔscIB relative to the default configuration (K562 TADs + GO Biological Process, scIB = 0.576; Figure 2b).

Among GO namespace alternatives, GO Molecular Function (GO MF; 227 modelled terms) reaches near-parity in integration performance, with 92.25% of terms achieving AUROC > 0.55 in the cell-type discrimination task, confirming it as a viable alternative when molecular-function-level interpretation is preferred. GO Cellular Component (GO CC; 677 terms) incurs a larger integration penalty, consistent with compartment annotations being less discriminative for haematopoietic cell identity than process annotations. Earlier GO releases (2014, 2019, 2022) perform increasingly worse than the 2024 release, reflecting growth in gene–GO annotations for immunity- and haematopoiesis-relevant biological processes over time.

Pathway ontologies show only modest integration penalties while retaining strong interpretability: Reactome [11] ($\Delta$scIB $= -0.017$) retains 88.57% of all 1,320 modelled terms with AUROC > 0.55; Pathway Commons [12] ($\Delta$scIB $= -0.014$) retains 81.92% of all 896 modelled terms. These results confirm that the GO importance framework extends to denser, less strictly hierarchical networks, and pathway-level alternatives remain viable when pathway-level annotation is the scientific goal.

For the gene co-regulation layer, tissue-mismatched TAD maps from breast cancer cell lines (T47D, MCF-7) are more detrimental than the lymphoid GM12878 map, consistent with partial TAD boundary conservation across haematopoietic lineages. Non-specific gene-gene interaction databases (BioGRID [13], GRAND GRN [14], TFLink [15], hTFtarget [16], STRING [17]) incur larger penalties than any ontology alternative, reflecting the absence of lineage-specific co-regulatory structure. A position-shifted K562 negative control, in which TAD boundaries are displaced by the mean TAD span, destroying co-regulatory groupings while preserving overall genomic statistics (see Methods), likewise reduces integration performance, confirming that genuine co-regulatory structure, not architectural sparsity, drives the TAD benefit.

The global shuffle control, in which all gene-GO assignments are randomly permuted while preserving annotation statistics (see Methods), serves as the key negative control for annotation content. Despite a modest integration penalty ($\Delta$scIB $= -0.008$), the Spearman correlation between GO importance scores from the shuffled and true model is only $\rho = 0.116$ (95% CI: $\approx \pm 0.030$; $p = 4.7 \times 10^{-6}$), computed for naive CD20$^{+}$ B IGKC$^{+}$ cells across all 1,791 modelled GO terms. This remaining correlation is easily explained by the fact that more abstract GO terms generally have a larger GO importance because their receptive field is larger, simply giving them information about more transcripts. This remains true in the shuffled case because the structure of the GO is retained, only the specific gene-GO assignments are randomised. This confirms that interpretive content derives from biologically correct annotations rather than from encoder architecture. Central interpretability results, discussed in the following, do not reproduce under the shuffled condition (Supplementary Figure 13).

##### 1.3 Generalisation to non-haematopoietic tissues

To assess generalisation beyond haematopoietic contexts, we benchmarked NetworkVI against MultiVI on three tissues from the Tabula Sapiens atlas [18], using lineage-matched TAD references for each tissue. NetworkVI matches MultiVI integration performance within run-to-run variability across all three tissues (liver: 0.563 vs. 0.566; lung: 0.584 vs. 0.592; large intestine: 0.586 vs. 0.576), confirming that the biologically structured encoder does not degrade performance outside immune contexts.

#### Supplementary Note 2: GO interpretability: extended results

##### 2.1 GO importance stability

To assess the robustness of GO importance scores to training stochasticity and dataset size, NetworkVI was retrained under three variant conditions on the NeurIPS 2021 CITE-seq BMMC dataset: an alternative random seed, and two cell-subsampled variants retaining 75% and 50% of cells, respectively. Spearman rank correlations of per-GO-term AUROC importance scores for the Naive CD20$^{+}$ B IGKC$^{+}$ comparison are: seed 1 vs. seed 0, $\rho = 0.681$ ($p = 1.98 \times 10^{-23}$); 75% subsample vs. full, $\rho = 0.456$ ($p = 1.62 \times 10^{-22}$); 50% subsample vs. full, $\rho = 0.413$ ($p = 1.31 \times 10^{-16}$). Seed stability exceeds subsampling stability, as expected: reseeding perturbs only weight initialisation and minibatch order, whereas reducing cell count also reduces the statistical power of the logistic regression underlying each GO importance estimate.

##### 2.2 Cell type similarity inference

We further explored the potential of GO importance values in distinguishing between various cell types and states for each modality, and their relevance for cross-modality comparisons (Supplementary Figure 15). To achieve this, we calculated Spearman correlation coefficients for all GO terms within each encoder and those present in the GO graph of the transcriptome and epitope modalities across each encoder for selected cell types (Supplementary Figure 14a,

16). As anticipated, we observed high correlations within (sub-)cell types, including subtypes of naive CD20$^+$ B cells, plasmablasts, plasma cells, progenitor cells, erythroid cells, and CD8$^+$ T cells. Notably, the correlation patterns for GO interpretability derived from the transcriptome modality and those from the epitope modality showed similarities, especially for the highest correlations. However, the correlation between GO importance values for the same cell types across the two modalities, represented by different encoders, was low, indicating that the encoders learn modality-specific GO representations.

We also calculate the Spearman correlation matrix of GO importances for substates within the CD8$^+$ T cell subpopulation (Supplementary Figure 16b). For CD8$^+$ T CD57$^+$ CD45RA$^+$/CD45RO$^+$ cells (terminally differentiated, senescent effector T cells or memory cells), we observe high importance for GO terms such as *regulation of programmed cell death* (GO:0043067), indicative of heterogeneous cytotoxic activity [19]. For CD8$^+$ T TIGIT$^+$ CD45RA$^+$/CD45RO$^+$ cells (regulated or exhausted CD8$^+$ T cells), high importances were found for *leukocyte activation involved in inflammatory response* (GO:0002269) and *B cell antigen processing and presentation* (GO:0002450), both shown to be downregulated in CD8$^+$ TIGIT$^+$ T cells [20]. Notably, high importance for an 'activation' term does not imply activation in the cell population, but rather that the genes associated with that activation are of high importance for cell identity, which can also be the case when the activation behaviour is inhibited.

While cell type similarity can also be inferred from node importances in a fully-connected encoder like MultiVI (Supplementary Figure 16c), the mean GO importance AUROC across cell types is 0.035 lower compared to NetworkVI (0.753). This difference is particularly pronounced for less frequent cell types (plasma cells, plasmablasts, and progenitor cells), which display less distinct similarity profiles in MultiVI. Additionally, the node importances of an MLP are not interpretable on a per-node level since there is no correspondence of the node to a specific biological concept.

##### 2.3 Sensitivity analysis: full marker gene results

For the in silico sensitivity analysis, cell-type-specific marker genes are sequentially set to zero across all cells, and the resulting changes in GO importance are recorded (Supplementary Figure 14b, Supplementary Figure 17a–e). This approach is analogous to the image degradation task used for quantitative evaluation of explainable models [21].

**CD19 (B-cell marker).** The absence of CD19 most significantly increased the GO importance for *regulation of sequestering of calcium ion* (GO:0051282; $\Delta$AUROC = 0.32, 95% CI: 0.30–0.34) and *leukocyte activation in an immune response* (GO:0002366; 0.21, 95% CI: 0.19–0.23) in B cells, followed by a decrease in *mononuclear cell proliferation* (GO:0032943; $-0.17$, 95% CI: 0.15–0.19). In lymphocyte progenitors, the largest decrease was observed for *regulation of localization* (GO:0032879; $-0.23$, 95% CI: $-0.19$ to $-0.23$) and in one of its child terms, *regulation of transport* (GO:0051049; $-0.16$, 95% CI: $-0.14$ to $-0.18$). Several calcium sequestration-related GO terms share the parent GO term *localization* (GO:0051179) with the identified terms. Calcium sequestration has been shown to differ between lymphocyte progenitors and differentiated B cells and in the context of B cell fate specification [22, 23].

NetworkVI does not prioritize all upstream terms equally, as their relevance depends on their utility in determining cell identity. The GO term *adaptive immune memory response* (GO:0090716) shows lower importance specifically in the CD4$^+$ T cell subset CD314$^+$ CD45RA$^+$ (0.17, 95% CI: 0.15–0.19), a population characterized by the CD314 marker, an activating receptor enabling cytotoxic functions [24]. Setting CD19 to zero in erythroblasts, normoblasts, proerythroblasts, and reticulocytes resulted in increases for *cell activation in immune responses* (GO:0002263; $\sim$0.21) and *leukocyte activation in immune response* (GO:0002366; $\sim$0.12), diverging from trends in B cells, likely reflecting the differentiation of these progenitors from lymphocyte progenitors.

**CD4, CD8A, and CD38.** Removing CD4, CD8A, or CD38 caused the most substantial GO importance changes in terms directly linked to each respective marker gene or its parent GO terms (Supplementary Figure 17b–d).

**IRF8 (pDC master regulator).** Setting IRF8 expression to zero placed *plasmacytoid dendritic cell activation* (GO:0002270) among the top 2% of GO terms with the largest importance change, confirming IRF8 as a primary driver of this term's discriminative power in pDCs, with no comparable effect in other cell types (Supplementary Figure 17e). This is consistent with IRF8's established role in pDC identity [25]. Broad changes in general immune GO terms reflect expected information propagation through the GO graph.

##### 2.4 Monocyte substate biological interpretation

The ICAM1-inflammatory state is present in both CD14$^+$ and CD16$^+$ monocytes (Figure 3e) and likely reflects a stimulus-induced, reversible activation program. ICAM1 facilitates endothelial adhesion critical for MHC–TCR interactions [26].

While the $CD14^{+}$ transitional-inflammatory subgroup showed slight enrichment for FCGR3A, the $CD16^{+}$ counterpart did not re-express CD14. Conversely, within the $CD16^{+}$ ICAM1-inflammatory subgroup, we observed low-level CD14 expression. These asymmetries suggest that the transitional-inflammatory state marks a gradual shift originating in $CD14^{+}$ monocytes and progressing toward $CD16^{+}$ identity, whereas the ICAM1-inflammatory state reflects a shared, stimulus-driven activation program adopted by both monocyte subsets. The presence of CD14 in inflammatory $CD16^{+}$ monocytes may reflect incomplete lineage transition or re-expression under inflammatory cues, consistent with findings that $CD14^{+}CD16^{+}$ bone marrow monocytes occupy an intermediate state in CD14/CD16 maturation [27].

Standard Louvain clustering approaches on the NetworkVI latent space did not recapitulate these states, as marker gene expression remained broadly distributed across clusters (Supplementary Figure 18d, 18e). We further benchmarked against a standard workflow including ORA/GSEA, GRNBoost2, and SCENIC (see Methods). None of these approaches recovered the identified functional programs, including GO:0002457, nor the corresponding substates. GRNBoost2 identified canonical immune regulators (e.g., JUN/FOS, CEBP), but the resulting programs were broad and did not resolve the antigen-presentation axis. SCENIC regulon activity failed to separate ICAM1-inflammatory and transitional-inflammatory substates (ARI $\approx 0$), indicating that these states are not captured at the level of regulon activity or transcription factor-centric programs.

A control model with shuffled gene–GO assignments did not reproduce the substates (Supplementary Figure 13a). As an additional stability check, we retrained NetworkVI under an independent random seed and reran HDBSCAN clustering on the GO:0002457 activation space. The second model independently recovers three substates whose cell-type assignments overlap 80-90% with those of the primary model, confirming that the substate decomposition is a stable property of the learned GO activation space rather than an artifact of weight initialization.

##### 2.5 NK substate biological interpretation

Via the described HDBSCAN clustering screening procedure we identified GO:0002519 (*natural killer cell tolerance induction*. This term has the highest silhouette score among NK-specific GO terms. HDBSCAN on the GO:0002519 activation space resolved four substates, Transitional NK (1%), Active NK I (12%), Active NK II (6%), and Mature NK (78%), labeled by differential expression and cross-referenced with [28] (Figure 3f-h). Transitional NK and Active NK I occupy an intermediate CD56bright-CD56dim position, co-expressing NCAM1 (CD56), TIGIT, and PTPN6 (hyporesponsive NK markers; [29]), and HAVCR2 (TIM-3) and FCGR3A, consistent with activated rather than exhausted states [30]. Active NK II is distinguished by SELL expression and Mature NK by high cytotoxic effector gene expression characteristic of fully differentiated CD56dim cells. Our substate classification captures tolerance-induction activity invisible to transcriptional clustering, complementing and extending the subsets described by [28].

##### 2.6 TAD importance gene-gene pairs weights

NetworkVI incorporates a TAD-informed co-regulation layer implemented as sparse residual connections among genomically co-localized genes; the learned residual weights are directly interpretable. Across the dataset, gene pairs assigned the highest residual connection weights frequently exhibit strong functional relationships. For instance, CD4-LAG3 reflects a known immune interaction in which LAG3 serves as an immune checkpoint receptor competing with CD4 for MHC class II binding [31], while H2BC9-H3C6 represents co-regulated histone genes [32]. Another example is CCR10-ENSG00000267042, where the latter is an antisense transcript to CCR10, though its regulatory role remains uncharacterized. Beyond these cases, a substantial number of high-weighted pairs include at least one gene lacking functional annotation.

##### 2.7 TAD importance: alternative TAD source comparison

Gene-level and TAD-group-level importance deltas were compared across four alternative connectivity sources relative to the K562 baseline. Tissue-mismatched TAD maps from T47D and MCF7 reduce mean gene importance by 1.4% (95% CI: 1.1-1.7%) and 0.9% (95% CI: 0.6-1.2%) and TAD-group importance by 2.5% (95% CI: 2.2-2.8%) and 1.9% (95% CI: 1.6-2.2%) respectively, confirming that biologically appropriate co-regulatory structure, not architectural complexity, drives the observed deltas. The lineage-matched GM12878 lymphoblastoid map yields a modest positive improvement (gene importance: +1.3%, 95% CI: 1.1–1.5%; TAD-group importance: +1.9%, 95% CI: 1.7–2.1%), consistent with partial TAD boundary conservation across haematopoietic lineages [33]. The position-shifted K562 control shows no meaningful change in either metric (both <0.03%), confirming that the deltas require biologically coherent co-regulatory groupings rather than architectural sparsity alone.

##### 2.8 TAD importance: confounder analysis and ChIP-seq validation

To assess whether TAD importance deltas reflect genuine co-regulatory signal rather than confounding technical or structural features, we tested four potential sources of bias (Supplementary Figure 19b–e). None showed a significant association with TAD importance deltas: TAD size ($\rho = 0.01$, $p = 0.76$); Hi-C contact domain score ($\rho = 0.002$, $p = 0.96$); HVG normalised dispersion ($\rho = 0.016$, $p = 0.66$); and CpG island density ($\rho = -0.069$, $p = 0.065$). The near-significant trend for CpG density is not unexpected, as active TADs tend to enrich for CpG islands; however, with less than 10% of variance explained, we do not expect substantial confounding.

Notably, the average improvement in TAD-level importance across all cell types is positively correlated with the variance in TAD importance improvement (Spearman's $\rho = 0.49$, 95% CI: 0.413–0.556, $p = 2.8 \times 10^{-44}$), further indicating that the model leverages TAD information in a cell type–specific manner (Supplementary Figure 20).

To provide orthogonal validation that the TAD co-regulation layer captures biologically meaningful regulatory signal, we correlated TAD importance scores from both the full model and the ablated baseline against ENCODE K562 ChIP-seq fold-change over control signals for 458 transcription factors and chromatin regulators across all 44 cell types (Supplementary Figure 21; full file list in Supplementary Table 12). The full model shows consistently higher positive correlations with the chromatin binding landscape compared to the ablated baseline (mean Spearman $\rho = 0.022$ vs $-0.017$; Wilcoxon $p < 10^{-300}$). The factors with the largest correlation deltas (full model minus baseline) include the cohesin subunit RAD21, among the strongest predictors of TAD organisation [34]; the PRC2 component SUZ12; and POLR3G, forming a functionally coherent group of domain-level regulators. We caution that this alignment does not establish causality, as these factors bind broadly.

##### 2.9 TAD-Driven Co-Regulation Uncovers lncRNA Co-Regulatory Activity

To further explore the role of the co-regulation layer, we examined two TADs from the top 10% (Supplementary Figure 18) showing the largest increases in TAD-level importance (Figure 4d). To provide broader context, Supplementary Figure 22 shows the TAD importance delta profiles across all 44 cell types for the ten TADs with the largest mean importance deltas. The majority of top-delta TADs independently show established co-regulatory patterns (e.g. the glycophorin cluster (GYPA/GYPE/GYPB), PPP2R5D/PTCRA, AREG/EREG) , while KIF14/CAMSAP2, the TAD with the highest mean delta, shows a broad gain across nearly all cell types rather than a lineage-specific pattern, suggesting that some top-delta TADs reflect general co-regulation. This motivated the selection of TADs with cell-type-specific profiles for detailed mechanistic interpretation: one containing exclusively protein-coding genes with partial GO annotation (TAD 1: FGD2/PI16/C6orf89) and one containing a single annotated gene alongside two lncRNAs (TAD 2: ERAP1/lncRNAs).

The first TAD, encompassing FGD2, PI16, and C6orf89, shows substantial increases in TAD importance for CD14+ monocytes, erythroblasts, and normoblasts. Individually, these genes exhibit modest importance shifts ( 0.03-0.06; e.g. FGD2: AUROC 0.900, 95% CI 0.889-0.909; PI16: AUROC 0.907, 95% CI 0.889-0.924; C6orf89: AUROC 0.964, 95% CI 0.949-0.976 in CD14+ monocytes), but their combined co-regulatory effect within the TAD contributes a substantial gain ( 0.36-0.37). While FGD2 and PI16 are not typically linked to erythroid lineages, C6orf89 displays robust expression in progenitor cells. Its expression profile in erythroblasts and normoblasts (Figure 4e, left) suggests a potential regulatory role, warranting further investigation. Notably, the co-regulation layer leads to 2-5-fold increase in the gene importance of FGD2 and PI16, but not of C6orf89, further underscoring TAD-driven synergy. Two variants within C6orf89 (rs113960910-T at 6:36931829 and rs138549196-A at 6:36930976) have been associated with PI16 protein levels in blood in a UK Biobank GWAS (GCST90470229, GWAS Catalog; [35]).

A second illustrative example involves a sparsely annotated TAD containing only one GO-annotated protein-coding gene, ERAP1, alongside two lncRNAs: ENSG00000247121 (antisense to LNPEP and ERAP2, which are excluded from the dataset) and ENSG00000248734 (Figure 4e, right). Unlike the first example, gene expression provides limited cues about the lncRNAs' relative contributions (Figure 4e, right); the TAD layer is the only path by which they enter the GO graph.

Despite only minor changes in gene importance ( 0.01-0.02 in monocytes), the TAD-level importance increases 20-fold in both CD14+ and CD16+ monocytes, indicating that the three genes act non-additively. The residual connection weights in the co-regulation layer between ENSG00000247121 and the other two genes are stronger across all cell types than the connection between the latter pair, suggesting that ENSG00000247121 plays a dominant role in conveying regulatory information in this context (Figure 4f). This interpretation is further supported by a 1.4-fold and 1.5-fold enrichment of IRF1 and IRF2 ChIP-seq signal, respectively, at the ENSG00000247121 locus, as measured by ENCODE fold-change over control BigWig tracks (K562 cells, ENCFF843BFO and ENCFF102RGH). These transcription factors are known regulators of ERAP1 and ERAP2 expression during IFN-$\gamma$ signaling [36].

To further dissect the functional relevance of these lncRNAs, we performed a gene-level sensitivity analysis as previously applied for GO importances. For each gene in the TAD, we artificially set its expression to zero across all cells and measured the resulting change in TAD importance. In CD16+ monocytes, TAD importance deltas were abolished when any single gene was removed, indicating a non-additive interaction of all three genes. However, in CD14+ monocytes, the TAD-level importance remained largely stable following dropout of ERAP1 or ENSG00000247121 but decreased substantially (TAD importance increase of $0.05 \pm 0.042$ instead of $0.43 \pm 0.042$) when ENSG00000248734 was dropped out. This suggests that, while largely unannotated, ENSG00000248734 plays a disproportionately central regulatory role within this TAD in CD14+ monocytes, but not in CD16+ monocytes.

##### 2.10 TAD importance: negative controls using TADs with largest mean importance decreases

To assess whether the co-regulation layer selectively captures biologically coherent signal rather than uniformly inflating TAD importances, we examined the distribution of TAD importance changes across all 737 modelled TADs and identified the ten TADs with the largest mean importance decreases as negative controls (Supplementary Figure 23). Biologically, we would expect TADs containing functionally unrelated or co-localization-only gene pairs to show no gain or a decrease, which is what we observe. Negative-delta TADs include functionally unrelated pairs whose co-localization does not reflect co-regulation in hematopoietic cells (ATOSA/MYO5A; MFAP3/SAP30L-AS1), housekeeping gene combinations (TRIM52/LINC01962; CAMK1/BRPF1), and one biologically informative case: DYRK2/IFNG-AS1, where despite the immune relevance of the IFNG locus, the antisense lncRNA IFNG-AS1 lacks discriminative co-regulatory signal because IFNG itself is absent from the dataset. This contrasts directly with the ERAP1/lncRNA TAD, where the annotated anchor gene is present and lncRNA importance propagates through it. The fact that importance decreases for these biologically incoherent TADs, rather than remaining at baseline, confirms that the model is not inflating all TAD importances indiscriminately, but responding to the presence or absence of genuine co-regulatory context.

##### 2.11 Covariate attention: additional cell types and covariates

In contrast to the age-associated patterns observed in lymphocytic effector cells, no such associations were found for myeloid, plasmacytoid, or B-cell lineages. For these cell types, high attention values were observed for GO terms such as *upregulation of detection of biotic stimulus* (GO:0009595) and *appendage morphogenesis* (GO:0035107), which have been associated with immunosenescence processes.

Additionally, GO terms such as *protein metabolic process* (GO:0019538), *cell development* (GO:0048468), *cellular process* (GO:0009987), *platelet activation* (GO:0030168), and *biological regulation* (GO:0065007) displayed higher attention values for smoking status and BMI, in line with their physiological relevance (Supplementary Figure 24).

#### Supplementary Note 3: GO importance analysis for additional perturbation genes in the Perturb-CITE-seq dataset

To more comprehensively compare NetworkVIs interpretability with existing approaches, we compare it against GSEA and ORA baselines for additional six perturbation genes measured in [37] beyond CD58 (discussed in the main text): CD59, ACSL3, DNMT1, ILF2, CDK6, and MYC. All six were identified in [37] as modulators of the immune cancer resistance (ICR) signature or as direct immune-evasion candidates. GO importance scores represent AUROC values comparing perturbed versus unperturbed cells within each screen condition ($-$IFN$\gamma$, $+$IFN$\gamma$, and TIL coculture); maximum 95% CI width across all displayed values is 0.03.

A key observation consistent across all six genes is that GSEA and ORA identify few or no significant GO terms for the TIL coculture condition, the biologically most relevant screen for immune evasion, reflecting the limited statistical power of threshold-based enrichment methods at approximately 100 cells per perturbation guide. Where GSEA terms do reach significance, they predominantly do so at relaxed FDR thresholds ($FDR < 0.25$) and without screen specificity, appearing with comparable enrichment scores across all three conditions. NetworkVI, by contrast, detects a larger number of GO terms with $AUROC > 0.53$ and for CDK6 and CD59 concentrates antigen presentation and T cell activation terms specifically in the TIL coculture condition.

**CD59.** CD59 (protectin) is a GPI-anchored membrane protein that inhibits membrane attack complex formation; its overexpression in tumours suppresses complement-dependent cytotoxicity and CD8$^+$ T cell activation as shown for lung cancer [38]. CD59 co-occurred with CD58 as an enriched perturbation in the viability screen of [37]. NetworkVI identifies elevated importance of exogenous antigen processing and presentation, T cell selection, and leukocyte activation terms in the TIL coculture condition (Supplementary Figure 25a), consistent with this co-occurrence. GSEA

(Supplementary Figure 25b) captures humoral immune response and cell adhesion terms broadly across conditions without clear TIL coculture specificity.

**ACSL3.** ACSL3 activates long-chain fatty acids for incorporation into membrane phospholipids and lipid droplets, and is overexpressed in melanoma where it confers ferroptosis resistance via oleic acid uptake [39]. ACSL3 knockout represses the ICR signature [37]. NetworkVI GO importance is broadly distributed at low magnitude across metabolic and general immune terms without clear screen specificity (Supplementary Figure 25c), as is GSEA (Supplementary Figure 25d). Both methods reflect the indirect, metabolically mediated contribution of ACSL3 to immune resistance.

**DNMT1.** DNMT1 maintains DNA methylation patterns and is upregulated by MYC, suppressing the cGAS-STING innate immune sensing pathway through promoter hypermethylation as shown for triple-negative breast cancer [29]. DNMT1 knockout represses the ICR signature [37]. NetworkVI identifies modest importance for leukocyte cell-cell adhesion and somatic diversification of immune receptors terms, with slight TIL coculture enrichment (Supplementary Figure 25e). GSEA (Supplementary Figure 25f) identifies T cell activation and antigen receptor signaling terms without clear screen specificity.

**ILF2.** ILF2 (NF45) is a multifunctional nuclear protein involved in RNA splicing, DNA damage repair, and miRNA biogenesis; its expression is elevated in metastatic melanoma [40]. ILF2 knockout represses the ICR signature [37]. NetworkVI GO importance is low-magnitude and broadly distributed without screen-specific enrichment (Supplementary Figure 25g). GSEA (Supplementary Figure 25h) shows broader immune term coverage predominantly in the $+$IFN$\gamma$ condition.

**CDK6.** CDK6 promotes G1/S cell cycle transition and is a component of the ICR resistance program [41]; CDK4/6 inhibition reverses this program and enhances T cell-mediated killing in patient-derived melanoma models. CDK6 has been directly linked to repression of antigen-presentation genes including MHC class I components. CDK6 knockout represses the ICR signature [37]. NetworkVI identifies elevated importance for T cell selection, peptide antigen processing and presentation via MHC class I, exogenous antigen processing and presentation, and adaptive immune response terms, concentrated in the TIL coculture condition (Supplementary Figure 25i), consistent with its documented role [41]. GSEA (Supplementary Figure 25j) captures overlapping terms but distributes them across all three conditions. ORA (Supplementary Figure 25k) identifies overlapping terms but limited to the $+$IFN$\gamma$ screen.

**MYC.** MYC knockout represses the ICR resistance program in melanoma cells, consistent with the association between MYC upregulation and increased resistance to immunotherapy [37, 41]. NetworkVI GO importance is broadly distributed at low magnitude without clear screen-specific enrichment (Supplementary Figure 25l). GSEA (Supplementary Figure 25m) and ORA (Supplementary Figure 25n) likewise identify diverse immune and non-immune terms across all conditions without screen specificity, consistent with the pleiotropic transcriptional effects of MYC.

**Conclusion.** NetworkVI and GSEA identify broadly consistent biological themes across all perturbation genes. Systematic differences appear in three respects. First, NetworkVI provides greater screen specificity for CDK6 and CD59, concentrating antigen presentation and T cell selection terms in the TIL coculture condition where GSEA distributes these more evenly. Second, GSEA returns a higher proportion of non-specific terms (wound healing, DNA damage, metabolic processes) alongside immune-relevant ones. Third, for perturbations with diffuse immune effects (MYC, ACSL3, ILF2, DNMT1), neither method yields a sharply interpretable signal; NetworkVI reflects this transparently through AUROC values near the random baseline rather than returning false-positive enrichments. The low absolute AUROC values ($\sim$0.505–0.57) across all genes are expected given the modest transcriptional effect sizes of individual CRISPR perturbations at $\sim$100 cells per guide; bootstrap CI widths should be examined alongside point estimates before interpreting differences as biologically meaningful.

#### Supplementary Figures

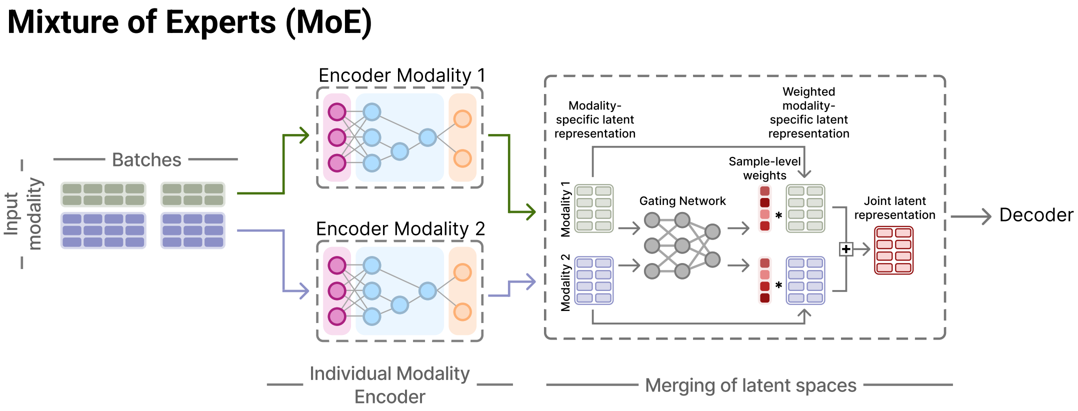


**Figure S1: Mixture of Experts framework in NetworkVI.**

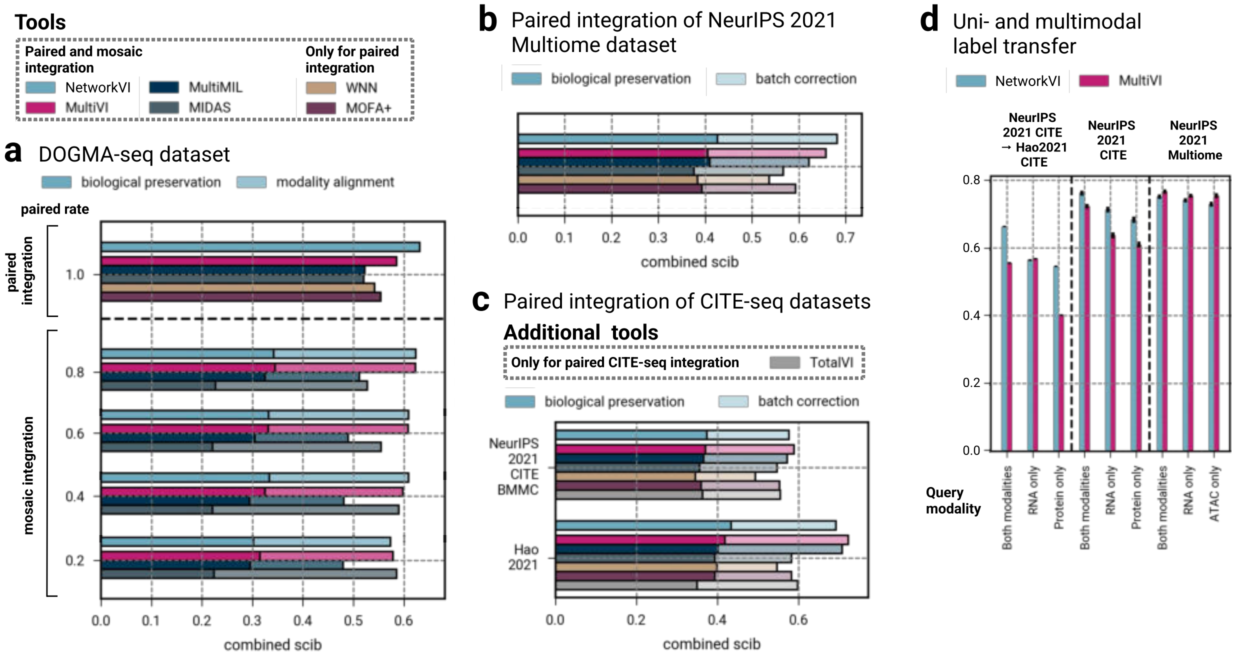


**Figure S2: NetworkVI achieves state-of-the-art performance for integration and query-to-reference mapping. a** For the DOGMA-seq dataset, NetworkVI demonstrates superior performance in both paired and mosaic integration scenarios, achieving the highest combined scib scores across 50 hyperparameter runs compared to other integration methods. The scib score consists of several metrics measuring the preservation of biological variance and the mix of modalities since no batch information is available for the DOGMA-seq dataset. NetworkVI performs on the level of state-of-the-art models for paired and mosaic integration of **b** Multiome and **c** CITE-seq datasets and maintains the best preservation of biological variance across the datasets. Concurrently, the other models outperform NetworkVI in modality mixing. **d** NetworkVI improves the cell label transfer across batches for the BMMC datasets and cross-tissue for the CITE-seq dataset by (Hao et al. 2021) for multimodal and unimodal queries. The error bars indicate the 95% confidence intervals via bootstrapping with 200 samples.

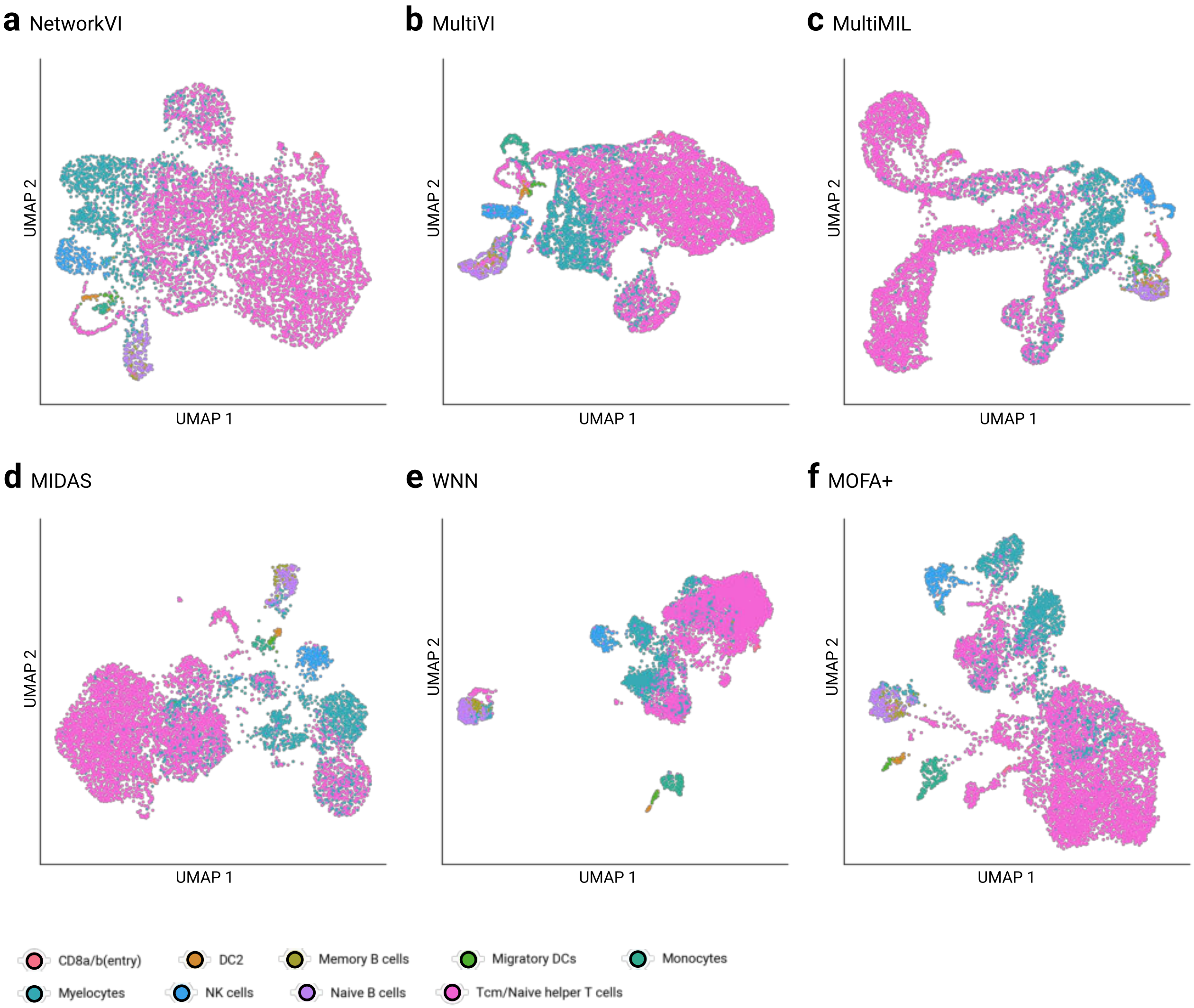


**Figure S3:** UMAPs of latent representations for the DOGMA-seq dataset generated by **a** NetworkVI **b** MultiVI **c** MultiMIL **d** MIDAS **e** WNN **f** MOFA+ for fully paired data using default values for tools.

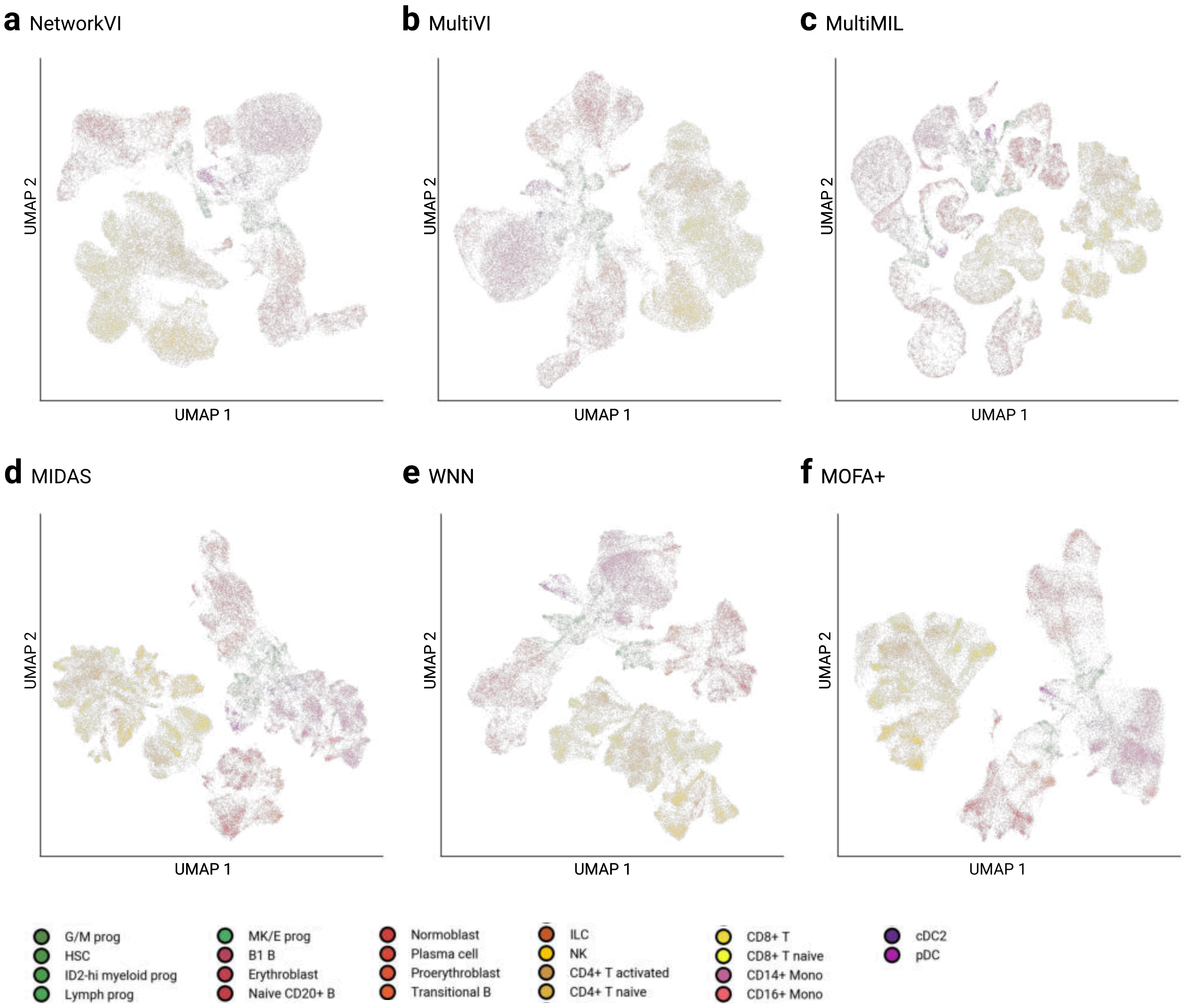


**Figure S4:** UMAPs of latent representations for the Multiome BMMC dataset generated by **a** NetworkVI **b** MultiVI **c** MultiMIL **d** MIDAS **e** WNN **f** MOFA+ **g** TotalVI for fully paired data using default parameters.

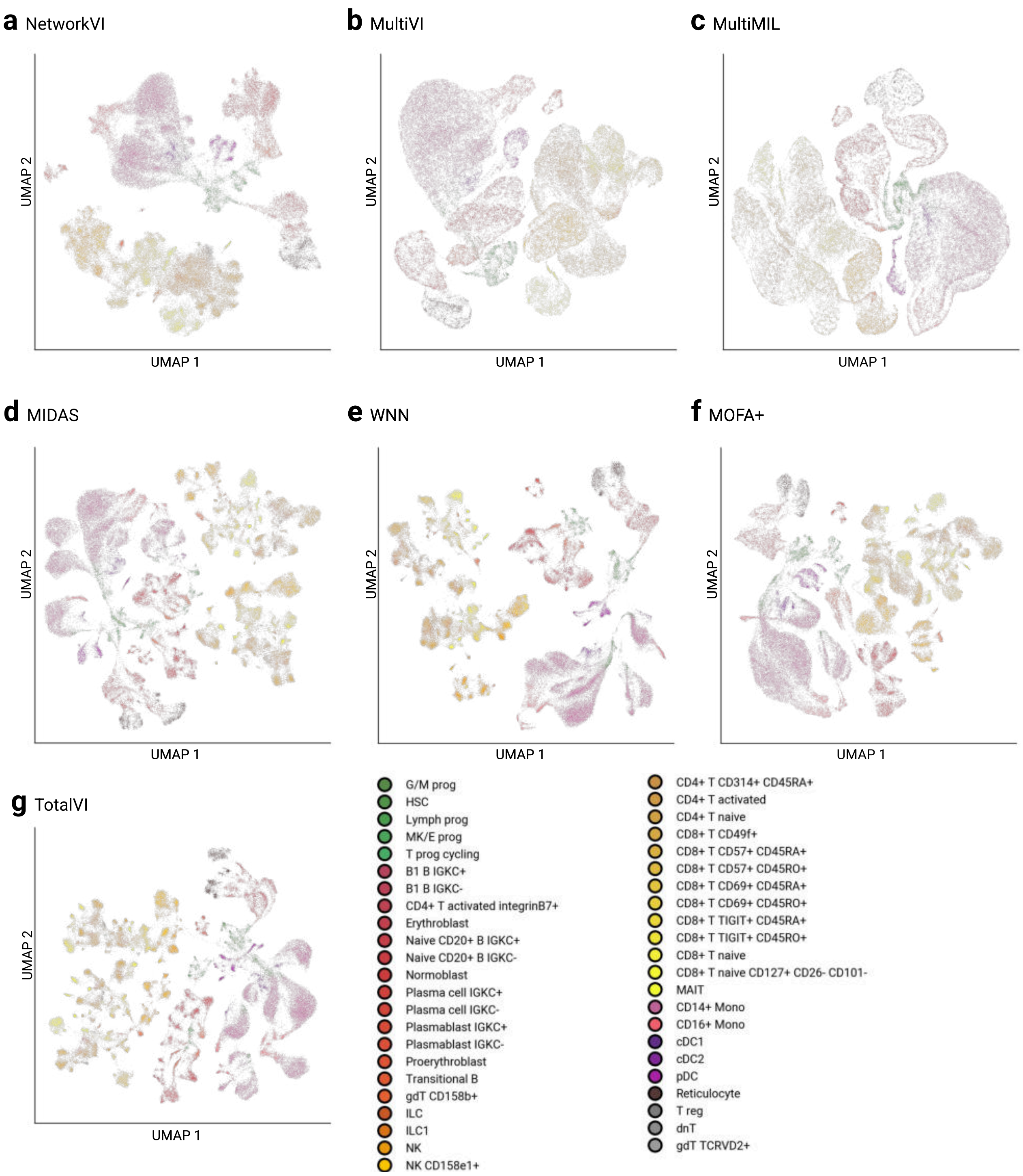


**Figure S5:** UMAPs of latent representations for the CITE- seq BMMC dataset generated by **a** NetworkVI **b** MultiVI **c** MultiMIL **d** MIDAS **e** WNN **f** MOFA+ **g** TotalVI for fully paired data using default parameters.

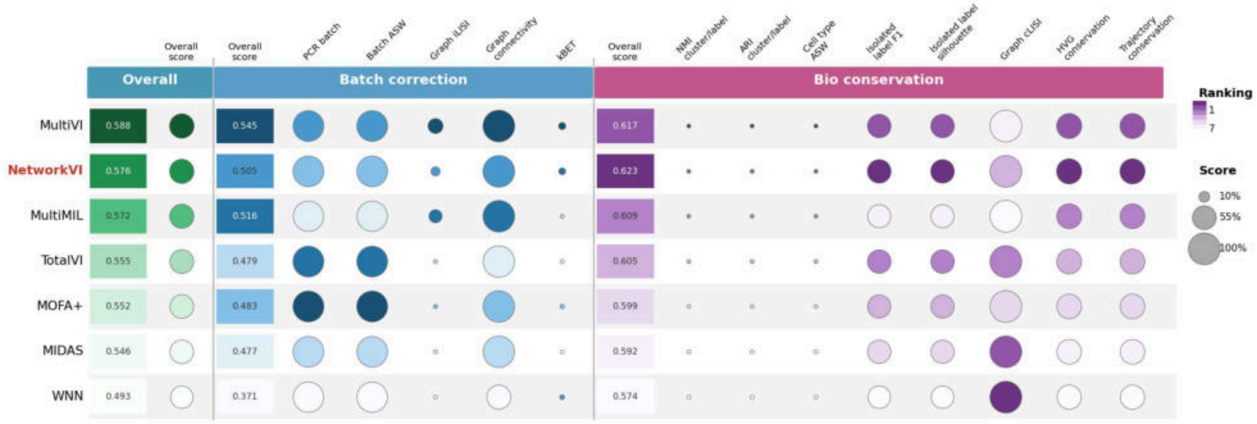


**Figure S6: scib scores for fully paired integration of the NeurIPS 2021 CITE-seq dataset.** Full breakdown of all individual scib sub-metrics across batch correction (PCR batch, Batch ASW, Graph iLISI, Graph connectivity, kBET) and biological conservation (NMI cluster/label, ARI cluster/label, Cell type ASW, Isolated label F1, Isolated label silhouette, Graph cLISI, HVG conservation, Trajectory conservation) dimensions, following the display format of Luecken et al. (2022). Circle size reflects the relative score (10-100%) and colour indicates rank among benchmarked methods (dark purple = rank 1, light = rank 7). Aggregate overall scores are shown for both batch correction and biological conservation separately. NetworkVI (highlighted) ranks among the top 4 methods across all batch correction metrics and achieves the highest score for 4 out of 5 biological conservation metrics, with the exception of Graph cLISI, where TotalVI, MIDAS, and WNN perform more strongly.

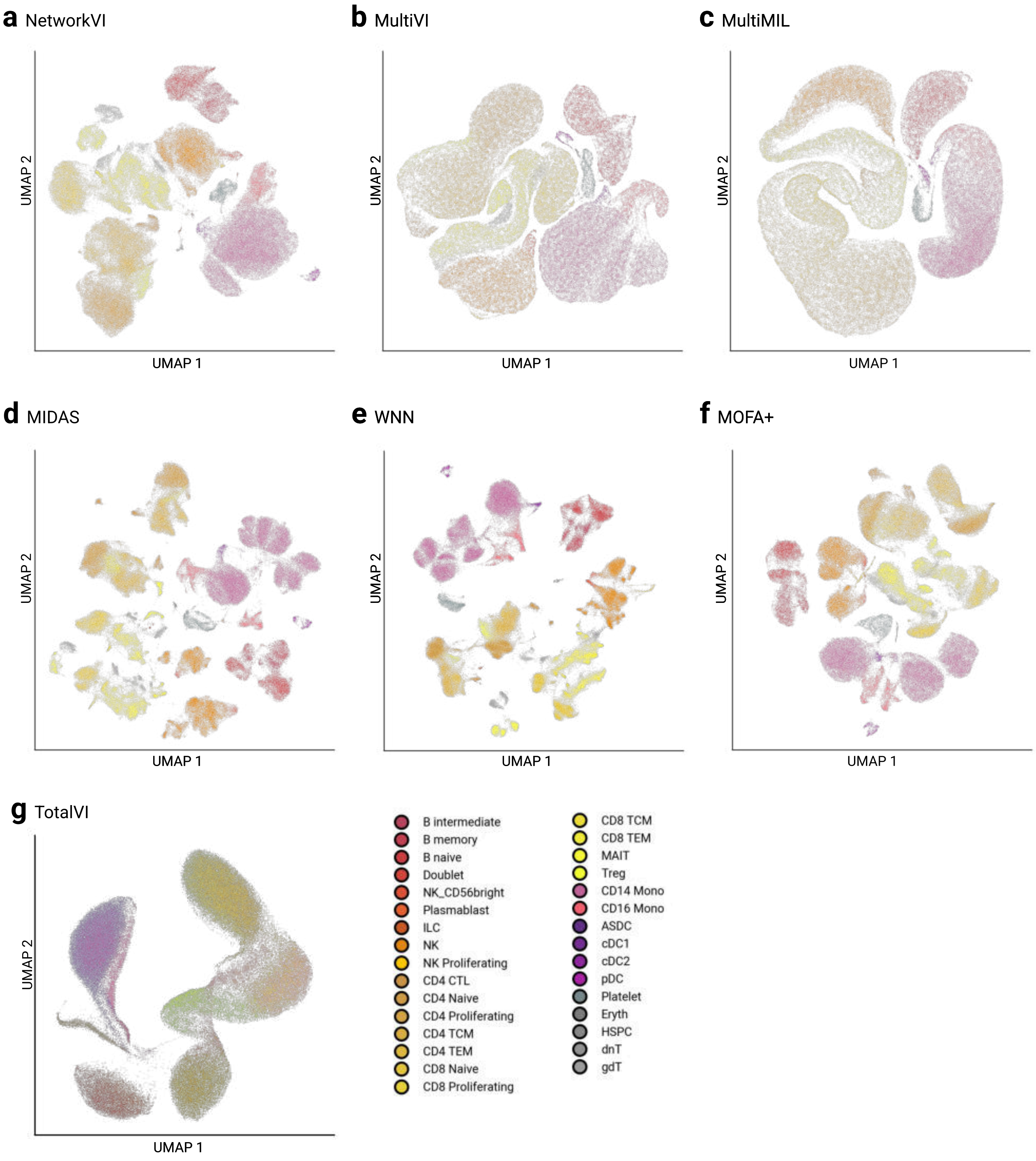


**Figure S7:** UMAPs of latent representations for the [3] dataset generated by **a** NetworkVI **b** MultiVI **c** MultiMIL **d** MIDAS **e** WNN **f** MOFA+ **g** TotalVI for fully paired data using default parameters.

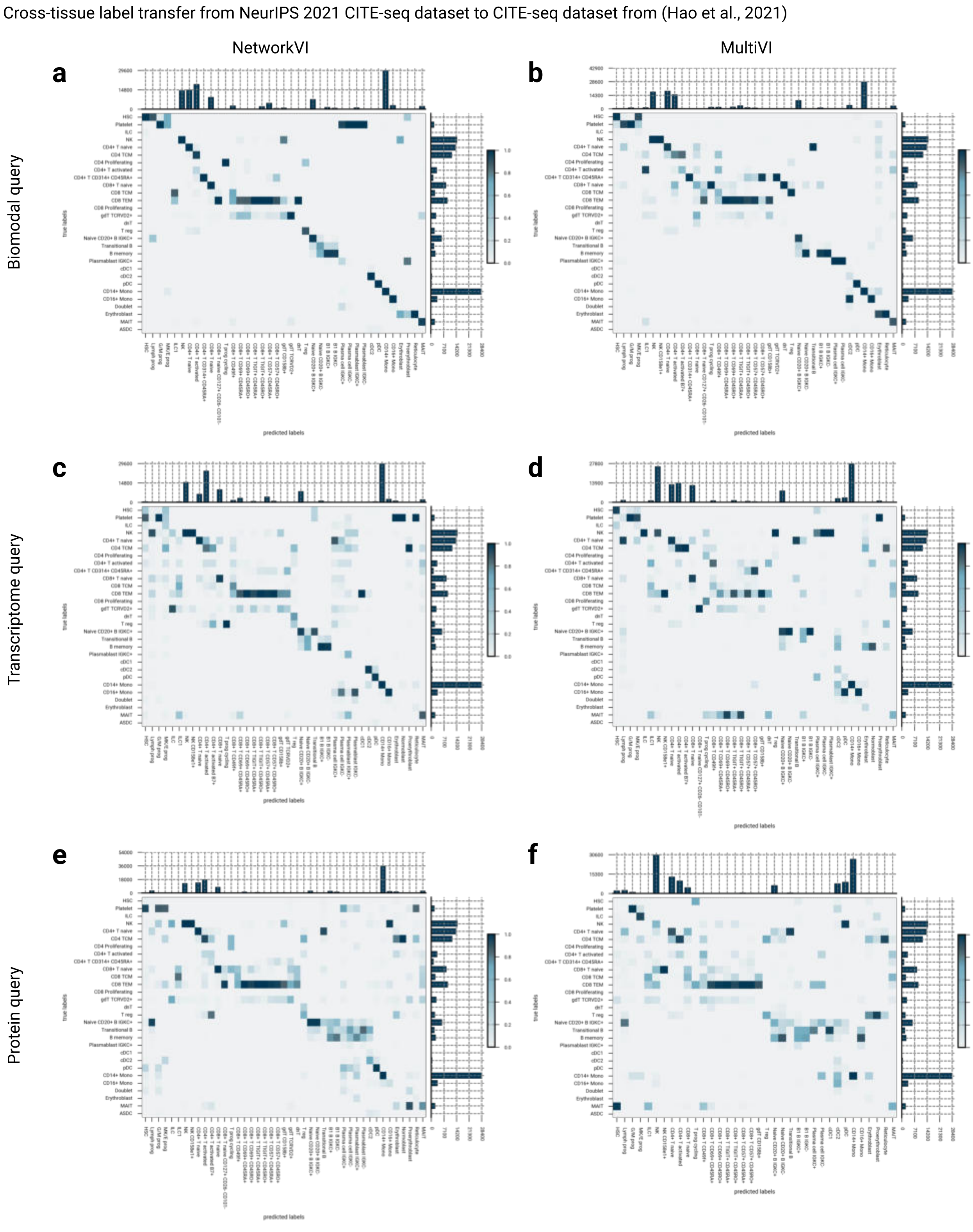


**Figure S8: NetworkVI enhances label transfer via reference-to-query mapping and a simple classifier.** Confusion matrix of NetworkVI (left, a, c, e) / MultiVI (right b, d, f) cross-tissue label transfer from the CITE-seq BMMC dataset to the [3] CITE-seq dataset performing a **a/ d** bimodal query, **b/ e** transcriptome modality query, **c/ f** epitope modality query. Misclassifications often can be traced back to closely related cell types; however in unimodal queries, the share of misclassifications to less related cells increases.

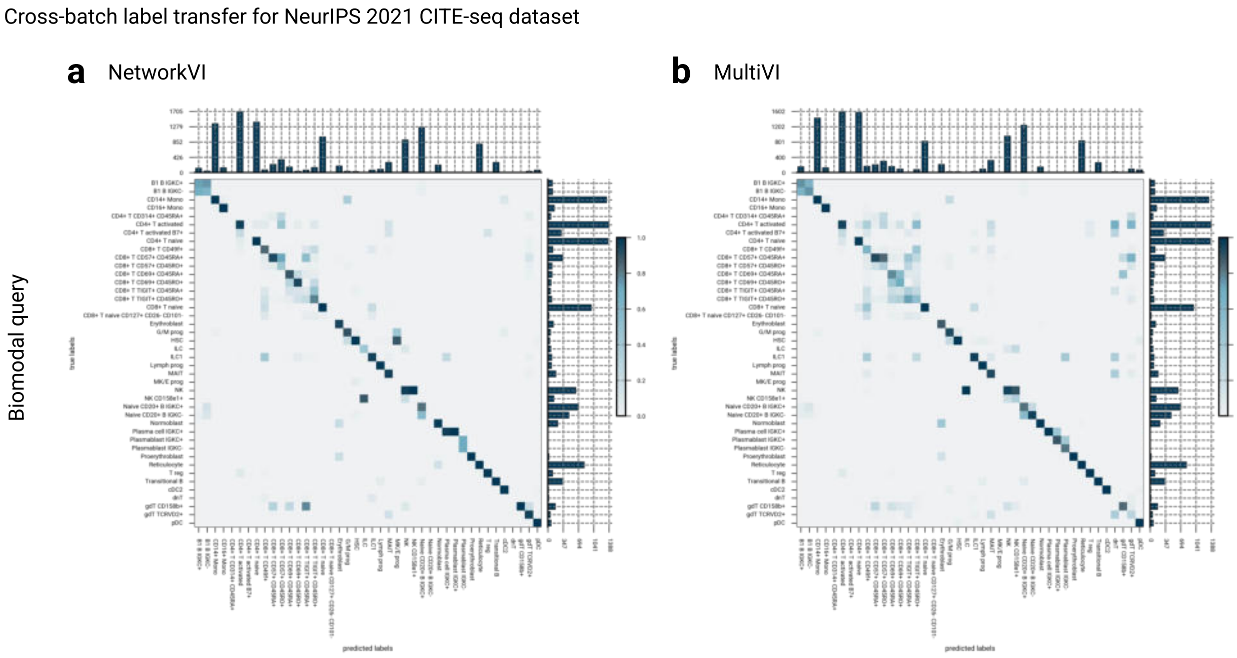


**Figure S9: NetworkVI enhances label transfer via reference-to-query mapping and a simple classifier.** Confusion matrix of **a** NetworkVI and **b** MultiVI label transfer on the CITE-seq BMMC dataset performing a bimodal query.

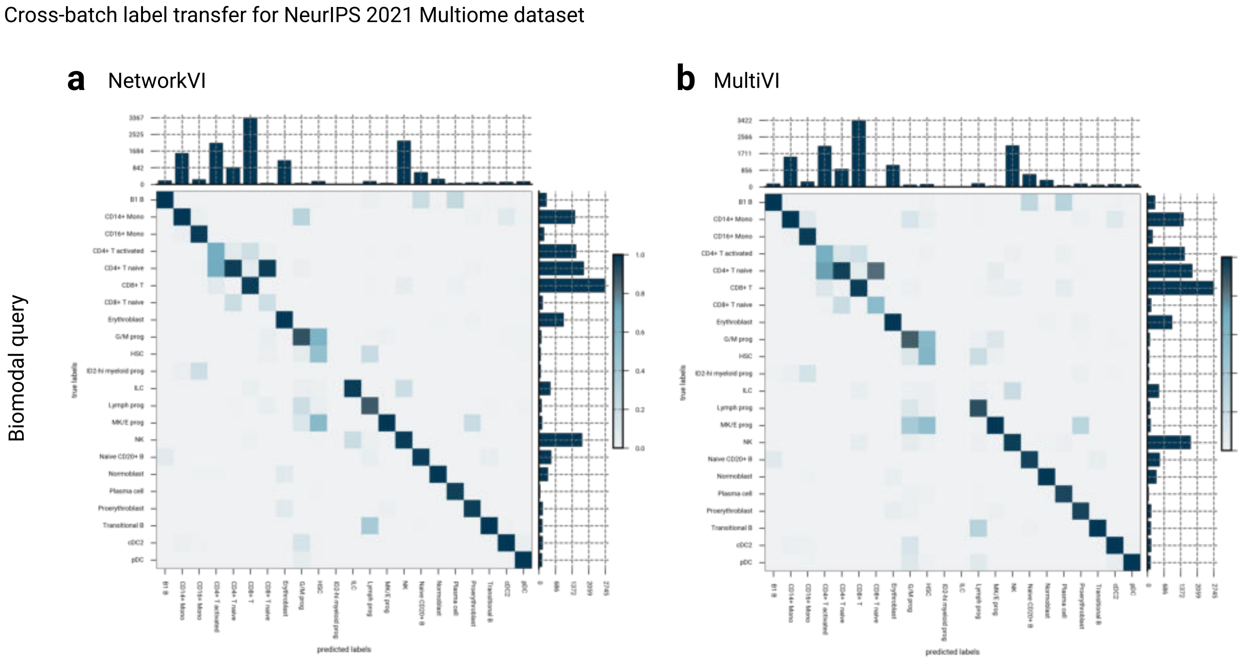


**Figure S10: NetworkVI enhances label transfer via reference-to-query mapping and a simple classifier.** Confusion matrix of **a** NetworkVI and **b** MultiVI label transfer on the Multiome BMMC dataset performing a bimodal query.

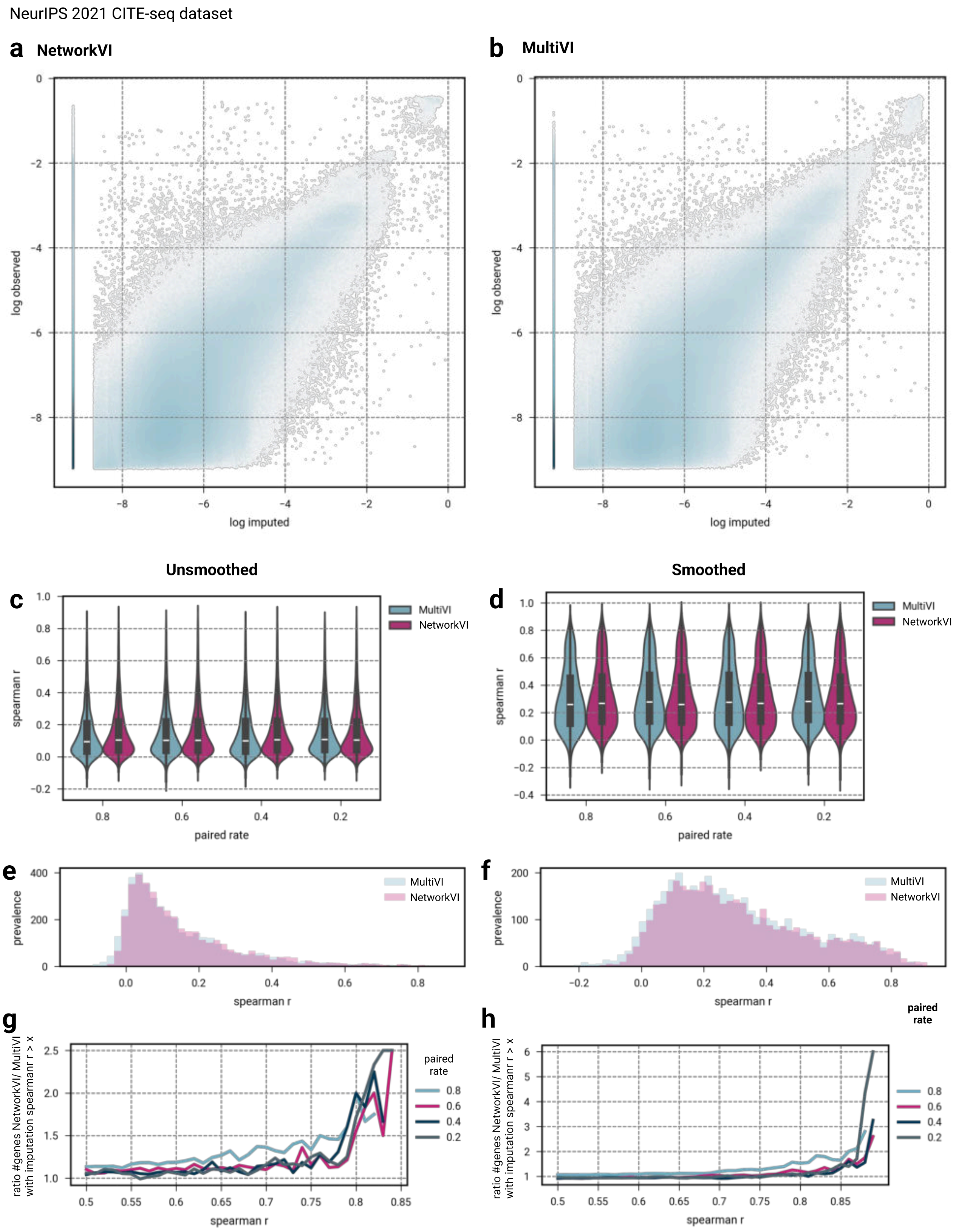


**Figure S11:** *See caption on next page.*

**Figure S11: NetworkVI enhances transcriptome imputation compared to MultiVI.** Hexagonal binning plots illustrating the imputed gene expression relative to the observed baseline for **a** MultiVI and **b** NetworkVI, utilizing the CITE-seq BMMC dataset with 40% of the dataset paired. **c** NetworkVI consistently outperforms MultiVI in per-gene expression imputation across pairing rates. **d** When assessing gene expression imputation on denoised un-normalized gene expression, NetworkVI and MultiVI perform equally across pairing rates. Histogram of per-gene spearman correlation between **e** raw or **f** denoised un-normalized gene expression data and imputed gene expression for a paired rate of 0.2. NetworkVI predicts a larger proportion of genes with a Spearman correlation coefficient exceeding 0.5, assessed on **g** raw and **h** denoised un-normalized gene expression data, across on different pairing rates. Here we show the ratio of the number of genes imputed at a given spearman r, so 1 would mean equal performance and larger values indicate NetworkVI outperforming MultiVI.

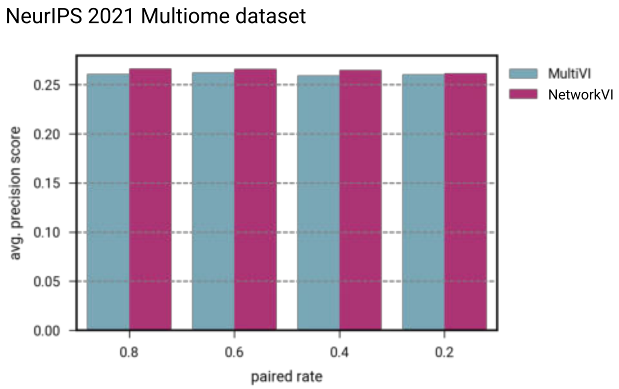


**Figure S12: NetworkVI enhances chromatin accessibility imputation compared to MultiVI.** The imputation performance is measured via the average precision for binary imputation of ATAC peaks within the Multiome BMMC dataset.

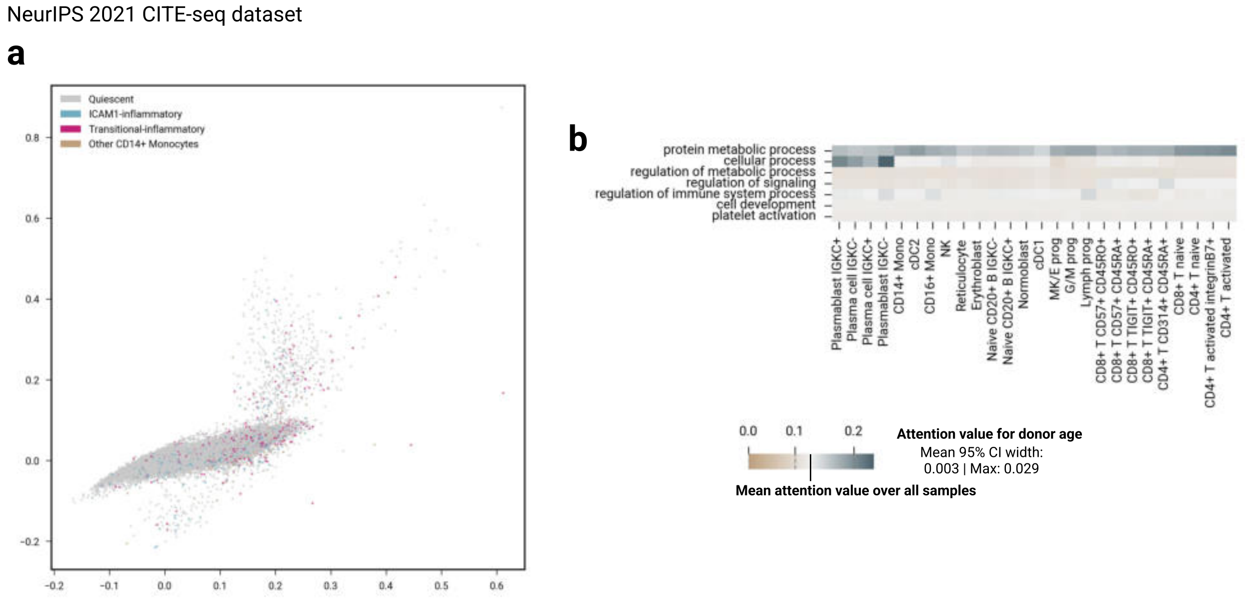


**Figure S13: Central NetworkVI interpretability results do not reproduce under global gene-GO shuffle negative control.** GO importance scores for Naive CD20$^+$ B IGKC$^+$ cells from the globally shuffled model (random permutation of all gene-GO assignments; see Methods, "Global shuffle negative control") does not recover the biologically expected pattern of high importance for humoral immune response terms and low importance for pDC activation terms. **a** GO activation space of T cell antigen processing and presentation (GO:0002457) in CD14$^+$ monocytes under the shuffled model. The three substates identified in Figure 3a (quiescent, ICAM1-inflammatory, transitional-inflammatory) are not recoverable from the shuffled activation space. **b** Covariate attention values for donor age under the shuffled model, shown using the same cell types and GO terms as Figure 5b. The age-associated attention pattern in lymphocytic effector cells is substantially attenuated relative to the true model (Spearman $r = 0.22$ between shuffled and true attention values for age; compare with $r = 0.61$ for the same seed under the true model, as reported in the main text).

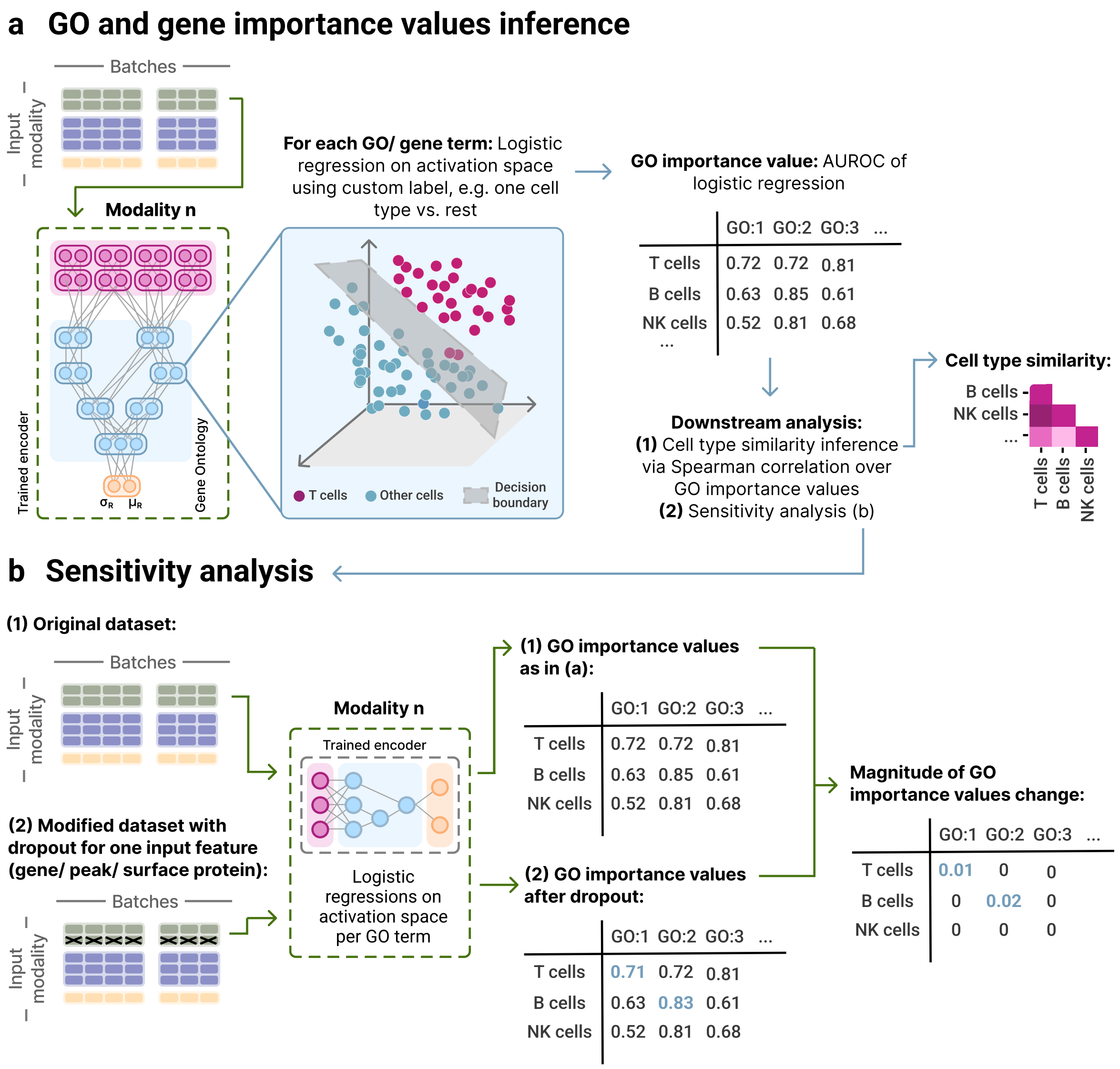


**Figure S14:** Flowchart of **a** the inference of GO and gene importance values and **b** performing a sensitivity analysis.

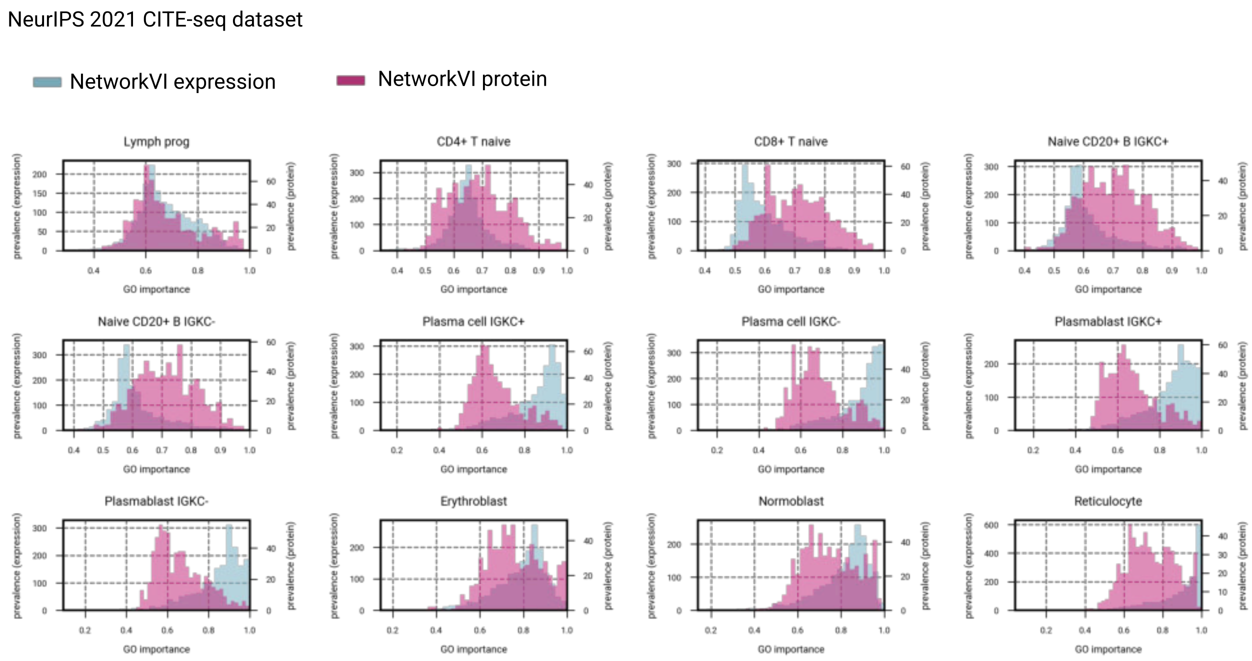


**Figure S15: Distributions of node importance values across cell types in CITE-seq BMMC dataset for GO terms in expression and protein encoder in NetworkVI.**

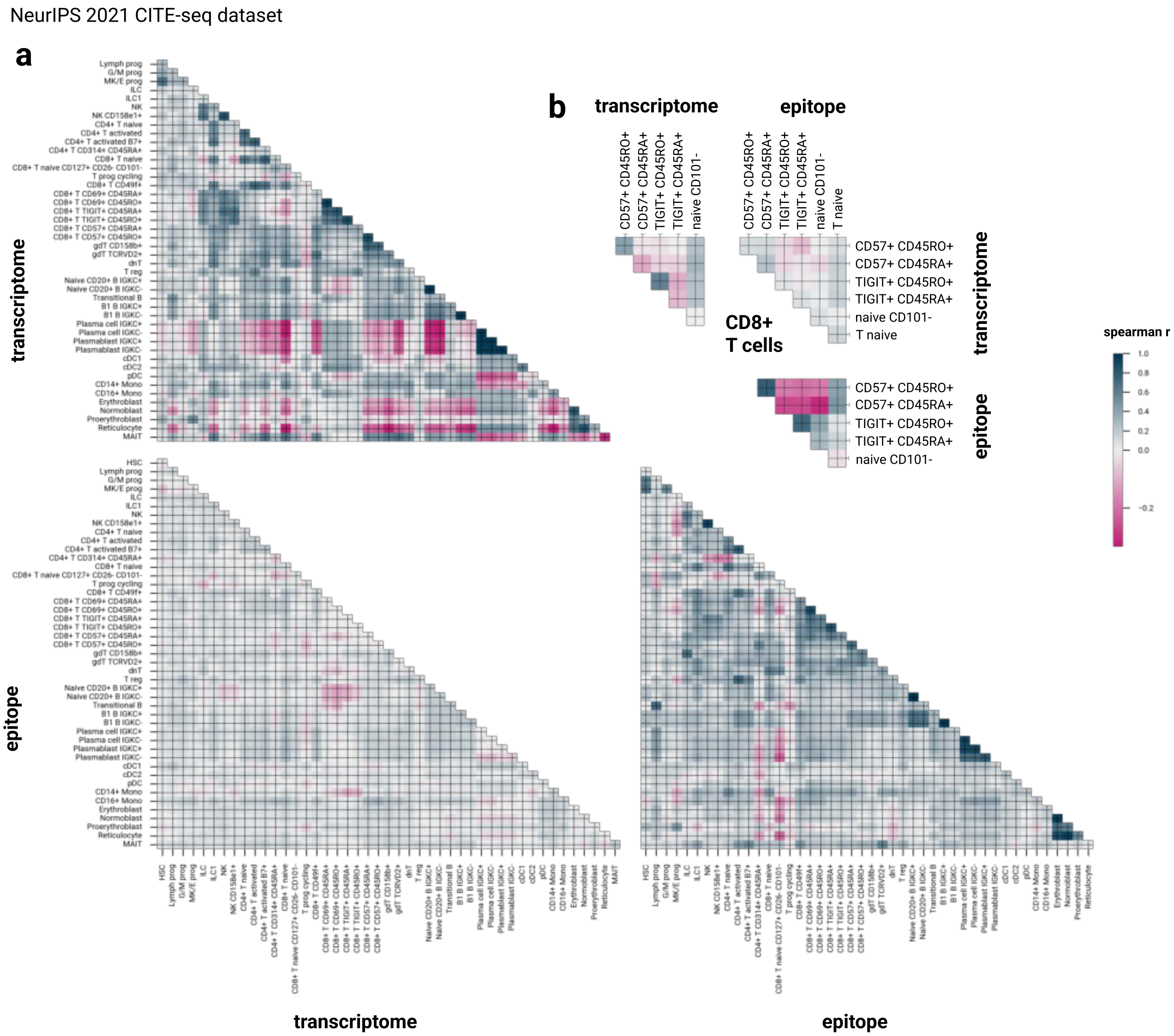


**Figure S16:** *See caption on next page.*

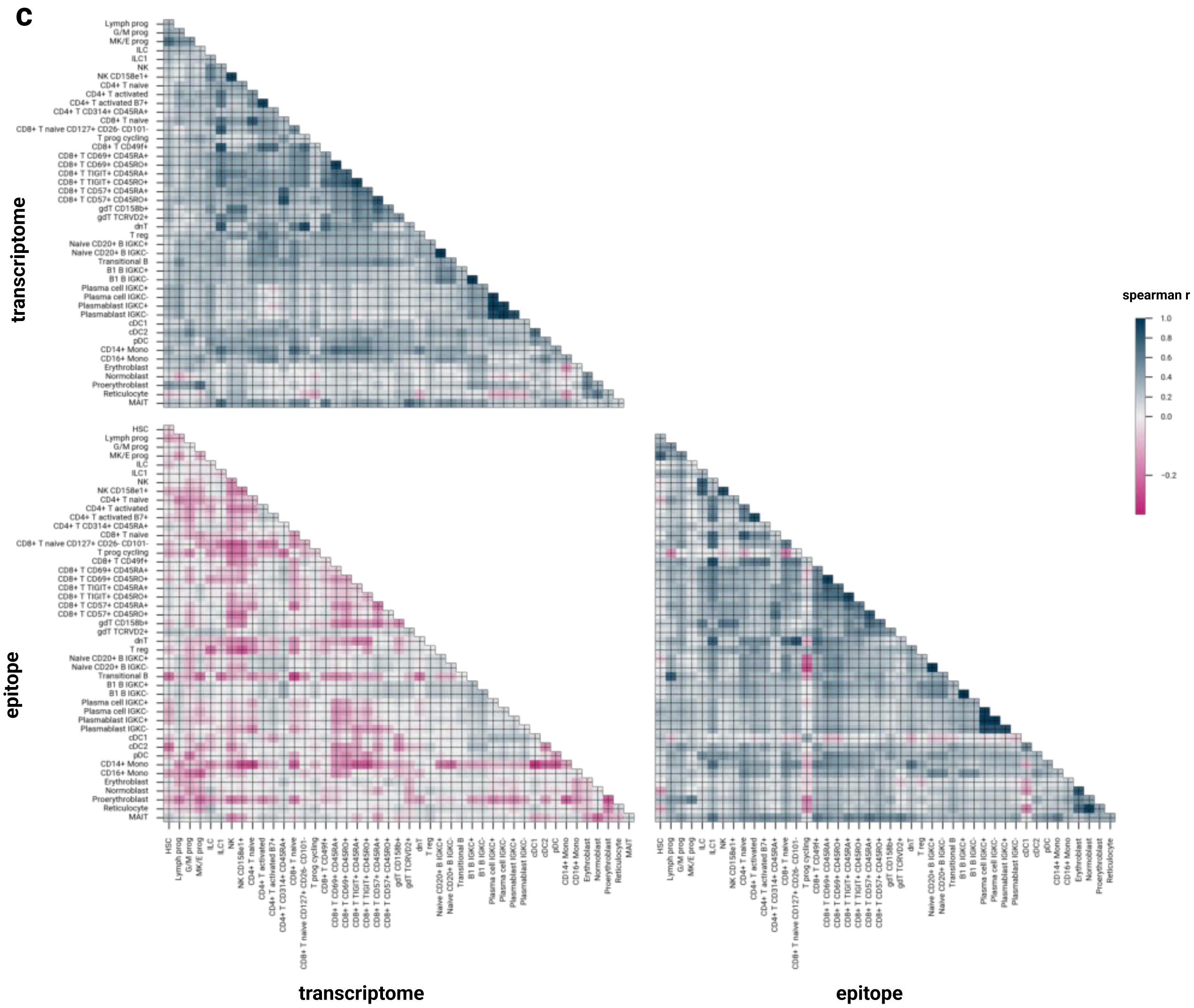


**Figure S16:** Using per-cell type GO importances, a Spearman correlation coefficient matrix measuring the cell type similarity can be inferred, indicating **a** for NetworkVI high correlations among sub-cell types, for instance, between different CD8+ T cells. Progenitor cells, plasmablasts, plasma cells, and erythroid cells show a distinct profile, as expected. **b** The Spearman correlation coefficient matrix calculated on CD8+ T cell subset demonstrates the ability of NetworkVI to explain distinct cell states. For instance, terminally differentiated CD8+ T CD57+ CD45RA+/CD45RO+ cells exhibit unique behavior for GO terms, such as *regulation of programmed cell death* (GO:0043067), indicative of heterogeneous cytotoxic activity across T cells. **c** Technically, a cell similarity matrix can be also inferred from importance values calculated for the fully-connected nodes in MultiVI. While the overall cell similarity aligns well to NetworkVI, the distinct profile of specific cell types, for instance the plasma cells and plasmablasts, is not captured. Also the encoders differ more strongly in their interpretability.

NeurIPS 2021 CITE-seq dataset **(GO upstream of dropout gene are bold.)**

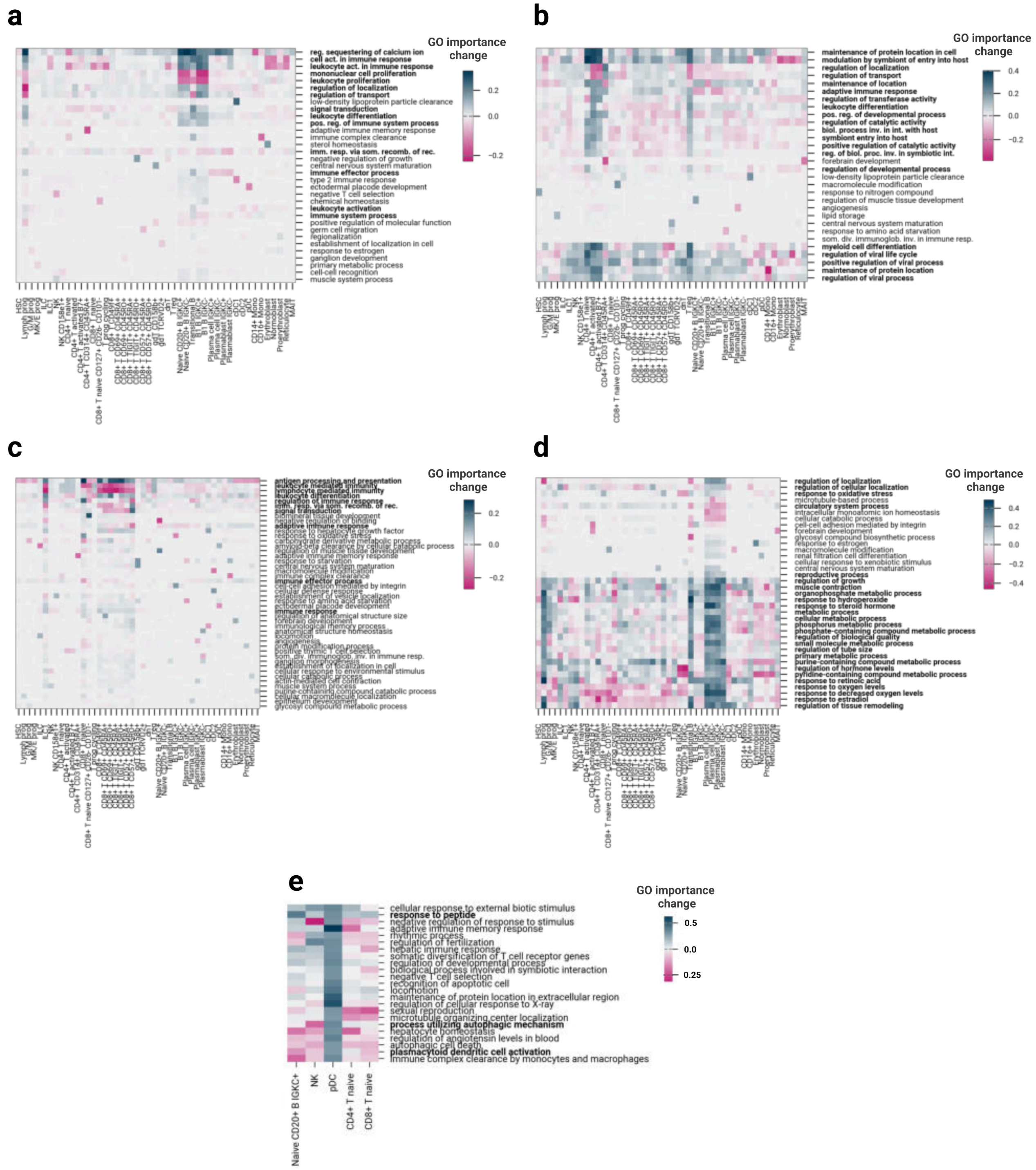


**Figure S17:** Sensitivity analysis on NetworkVI removing marker genes, **a** CD19 (Full data Figure 2e.) **b** CD4, **c** CD8A and **d** CD38 from the dataset. **e** Sensitivity analysis for IRF8 across selected cell types. GO importance changes are shown for all modelled GO terms after setting IRF8 expression to zero. GO terms are filtered to exclude those downstream of developmental process (GO:0032502), metabolic process (GO:0008152), cell proliferation (GO:0008283), and cell motility (GO:0048870) to focus on functionally interpretable immune-relevant terms. Plasmacytoid dendritic cell activation (GO:0002270) ranks among the top 2% of GO terms with the largest importance change specifically in pDCs, with no comparable signal observed in other cell types, confirming that the sensitivity framework correctly identifies IRF8 as a cell-type-specific driver of this GO term consistent with its known role in pDC identity. Broad changes in general immune GO terms reflect expected information propagation through the GO graph.

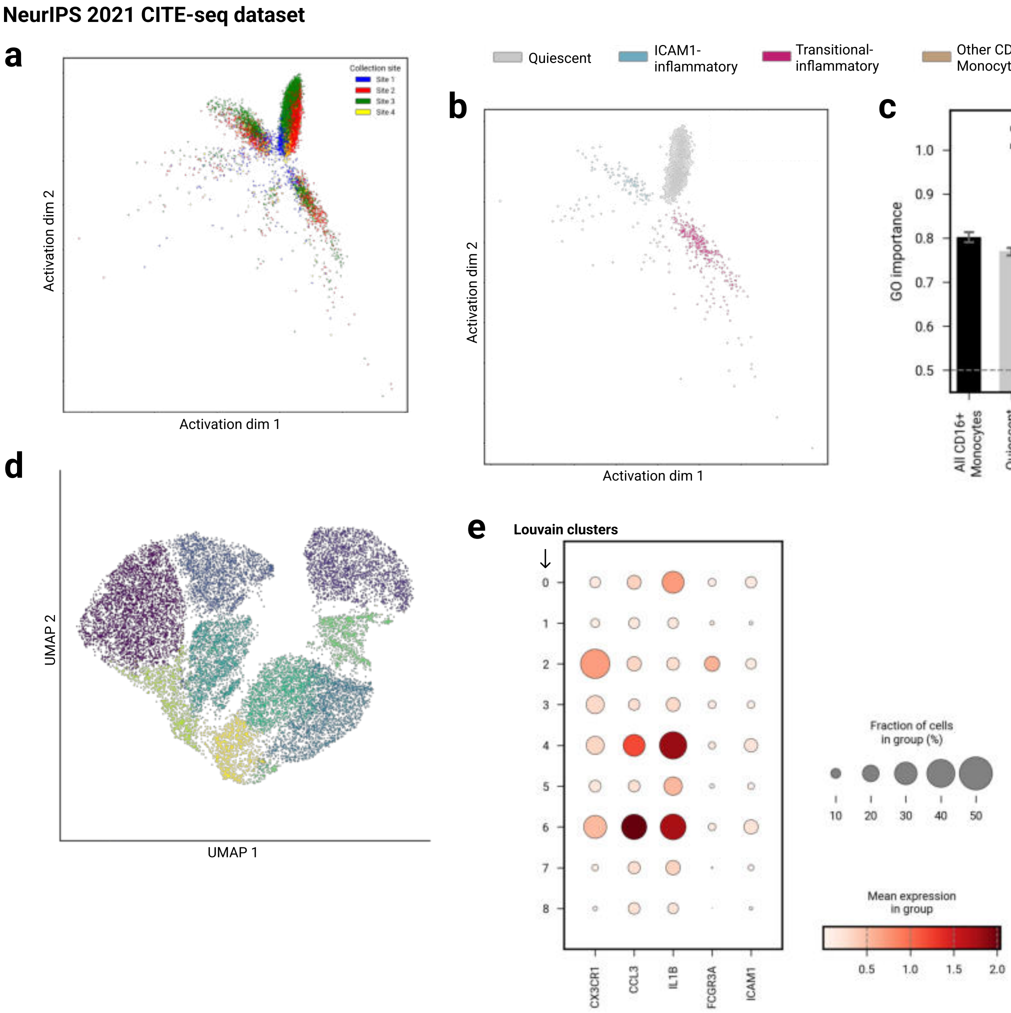


**Figure S18:** **a** GO activation projection (GO:0002457, *T cell antigen processing and presentation*) of CD14+ Monocytes, colored by collection site. **b** GO activation projection (GO:0002457, *T cell antigen processing and presentation*) of CD16+ Monocytes, colored by substates. **c** GO importances for subsets of CD16+ Moncoytes. Bootstrapped AUROC distributions (n=200 resamples) for each group. Bars indicate median AUROC across bootstrapped samples; error bars denote 95% bootstrap confidence intervals. Statistical comparisons between groups were performed using two-sided DeLong test on bootstrap distributions (***=p<0.001). **d** UMAP of transcriptional latent space (NetworkVI) of CD14+ Monocytes showing no spatial segregation of substates, colored by Louvain clusters. **e** Dot plot of substate-defining genes for Louvain clusters.

NeurIPS 2021 CITE-seq dataset

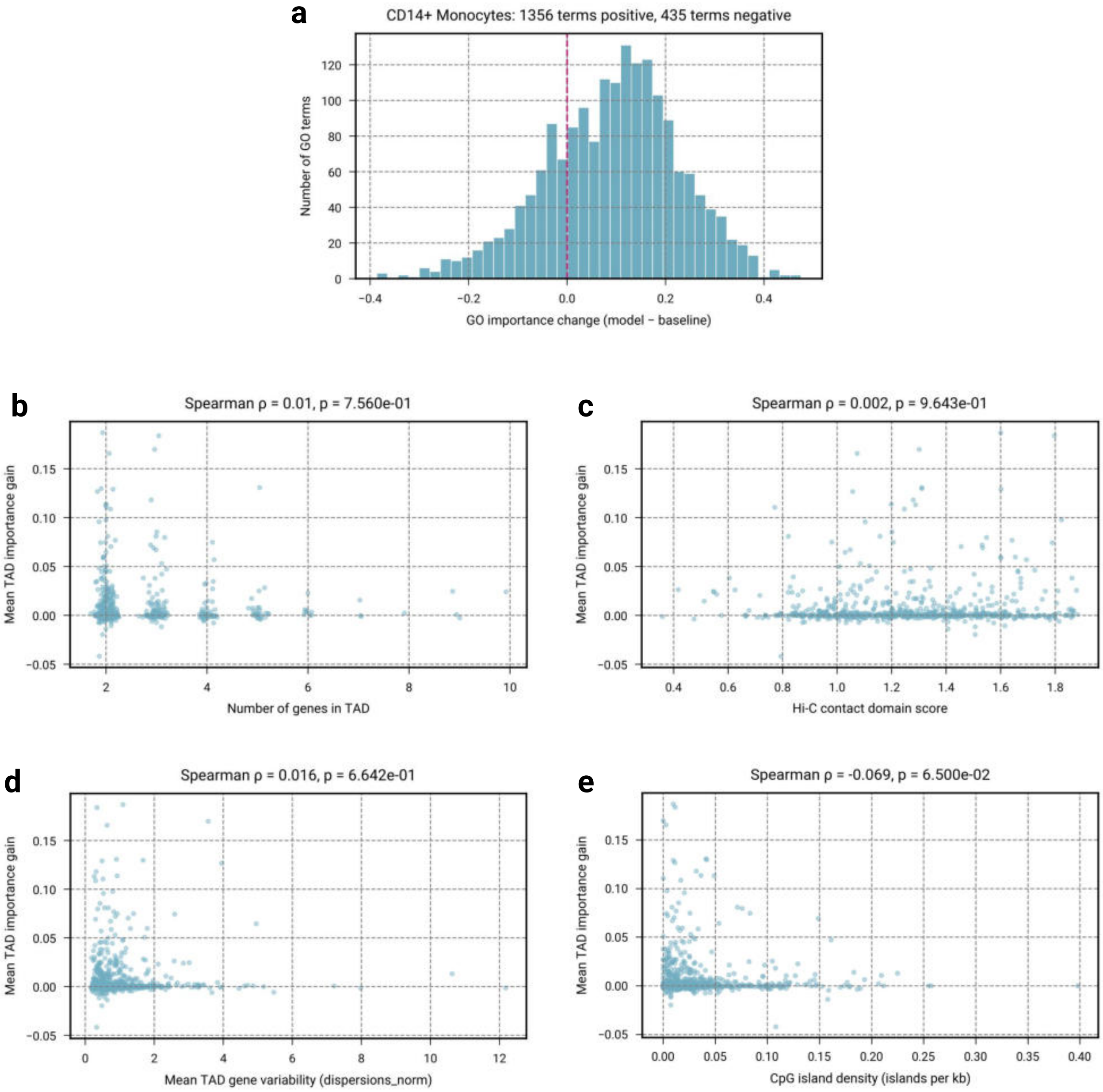


**Figure S19:** **a** Distribution of GO term importance changes upon addition of the TAD co-regulation layer for CD14+ Monocytes. Each bar represents the number of GO terms with a given change in AUROC importance (model minus baseline). Of 1,791 GO terms, 1,356 (75.7%) show a positive importance change, indicating that the TAD co-regulation layer broadly improves functional signal capture rather than benefiting a small subset of terms. The dashed pink line marks zero change. Mean TAD importance gain (model minus baseline AUROC, averaged across all 44 cell types) is independent of **b** TAD size (Spearman $\rho = 0.01$, 95% CI: $[-0.064, 0.085]$, $p = 0.76$, indicating that the observed importance gains are not a counting artifact of larger TADs containing more genes.), **c** Hi-C contact domain score (Spearman $\rho = 0.002$, 95% CI: $[-0.073, 0.075]$, $p = 0.96$, indicating that gains are not driven by TADs with the strongest chromatin insulation boundaries.), **d** constituent gene expression variability ( Spearman $\rho = 0.016$, 95% CI: $[-0.059, 0.087]$, $p = 0.66$, indicating that the co-regulation layer does not simply amplify already-variable genes but captures genuine co-regulatory context; constituent gene expression variabilit is represented by normalized dispersion of genes.), and **e** (Spearman $\rho = -0.069$, 95% CI: $[-0.141, 0.003]$, $p = 0.065$, indicating no significant association between local CpG density and TAD importance gain. CpG island density is calculated per TAD genomic locus (number of overlapping UCSC hg38 CpG islands normalized by TAD span in kilobases, $n = 722$ TADs).). Each point represents one TAD ($n = 722$).

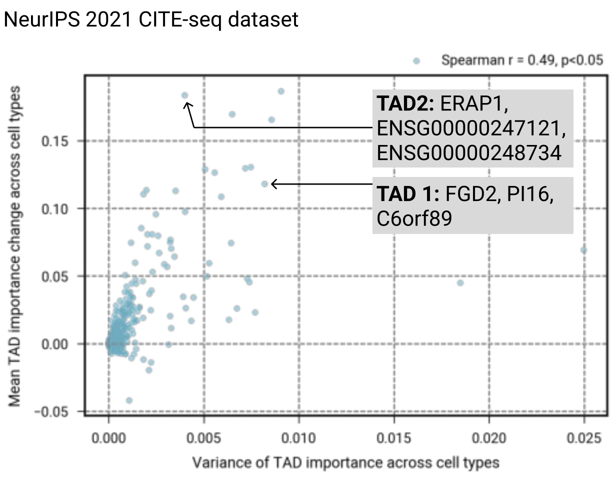


**Figure S20: Correlation between average and variance of TAD importance improvement across cell types suggests lineage-specific utilization of TAD-informed co-regulation.**

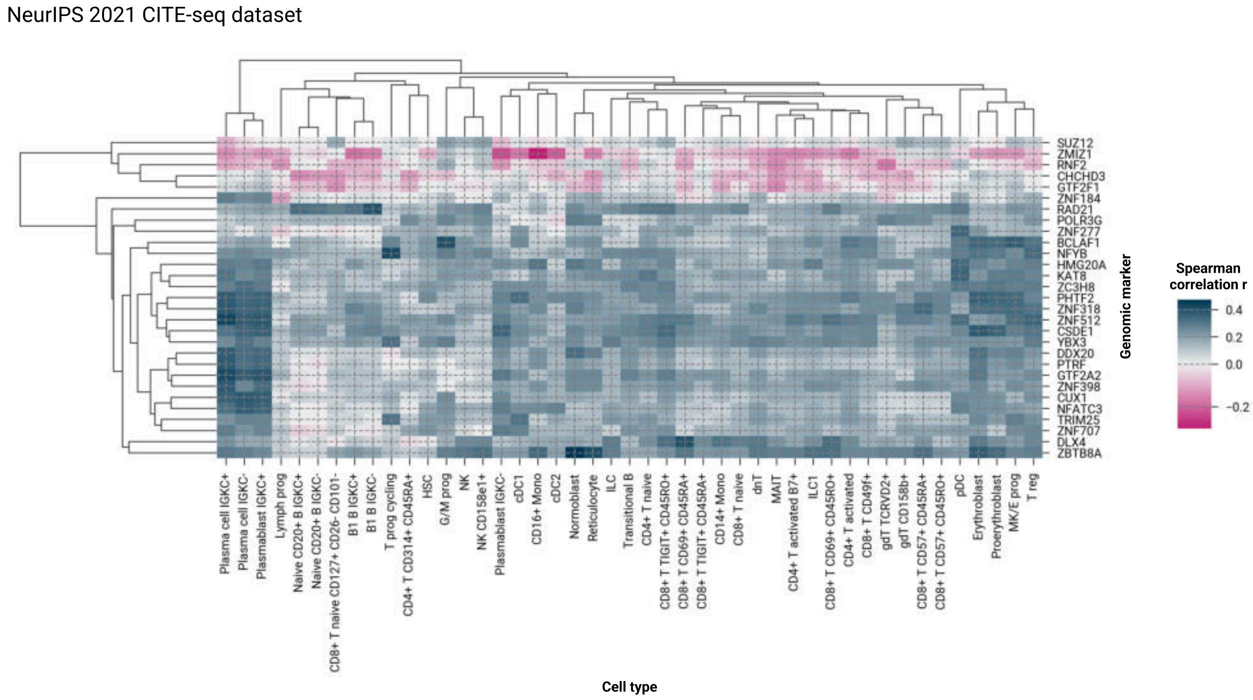


**Figure S21:** Correlation between TAD-level importance scores and genomic marker signals across transcription factors and chromatin regulators. Heatmap showing Spearman correlation coefficients between TAD importance scores (full NetworkVI model) and ChIP-seq signal p-value signals from ENCODE K562 bigWig tracks (Michael Snyder laboratory), across all 44 cell types (columns) and selected genomic markers (rows). Positive correlations (blue) indicate that TADs with higher importance scores in a given cell type tend to be bound by the corresponding factor. Hierarchical clustering is applied to both rows and columns. The full model (TAD co-regulation layer present) shows systematically higher correlations with genomic marker signals than the ablated baseline (Wilcoxon signed-rank $p < 10^{-300}$, rank-biserial $r = -0.110$).

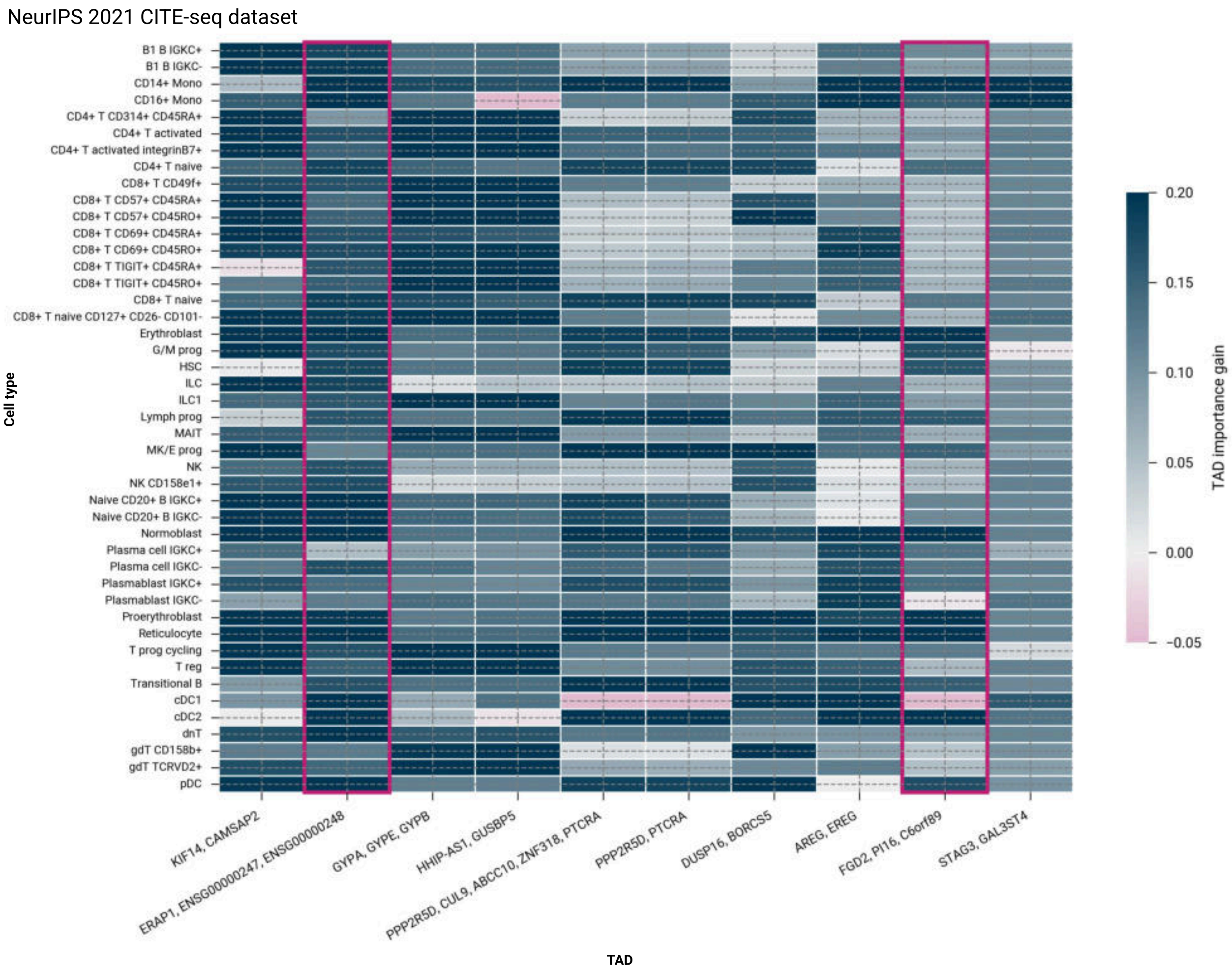


**Figure S22:** TAD importance gain profiles for the ten TADs with the largest mean importance gains across all 44 cell types. Each row represents a cell type and each column a TAD, with color encoding the difference in TAD-level AUROC importance between the full NetworkVI model and the ablated variant lacking the TAD co-regulation layer. The two TADs selected as representative examples in the main text (ERAP1/lncRNAs, rank 2; FGD2/PI16/C6orf89, rank 9) are highlighted with pink borders. Several top-gain TADs show lineage-specific patterns consistent with known biology: the glycophorin cluster (GYPA/GYPE/GYPB) gains importance selectively in erythroid populations; PPP2R5D/PTCRA gains importance in lymphoid lineages consistent with PTCRA's role in pre-T cell receptor signaling. KIF14/CAMSAP2 (rank 1) shows a broad gain across cell types reflecting general proliferation-associated co-regulation rather than lineage-specific biology.

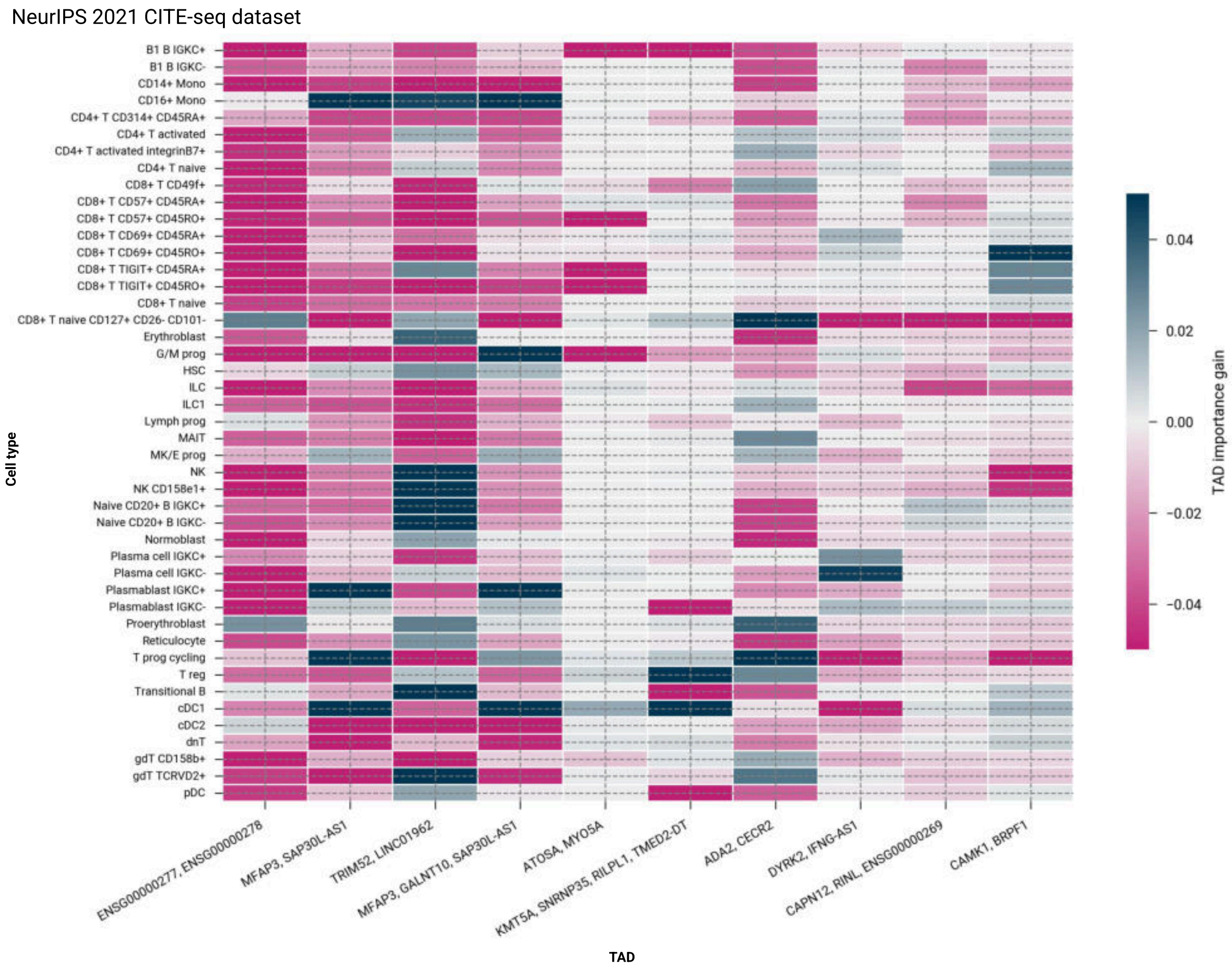


**Figure S23:** TAD importance gain profiles for the ten TADs with the largest mean importance decreases across all 44 cell types, serving as negative controls. TADs shown include functionally unrelated gene pairs (ATOSA/MYO5A; MFAP3/SAP30L-AS1), housekeeping gene combinations (TRIM52/LINC01962; CAMK1/BRPF1), and DYRK2/IFNG-AS1, an informative negative control in which the antisense lncRNA to the immune-relevant IFNG locus fails to provide discriminative co-regulatory signal in the absence of IFNG itself from the dataset. The absence of importance gain confirms that the model learns context-dependent co-regulatory signals rather than uniformly inflating TAD importances.

NeurIPS 2021 CITE-seq dataset

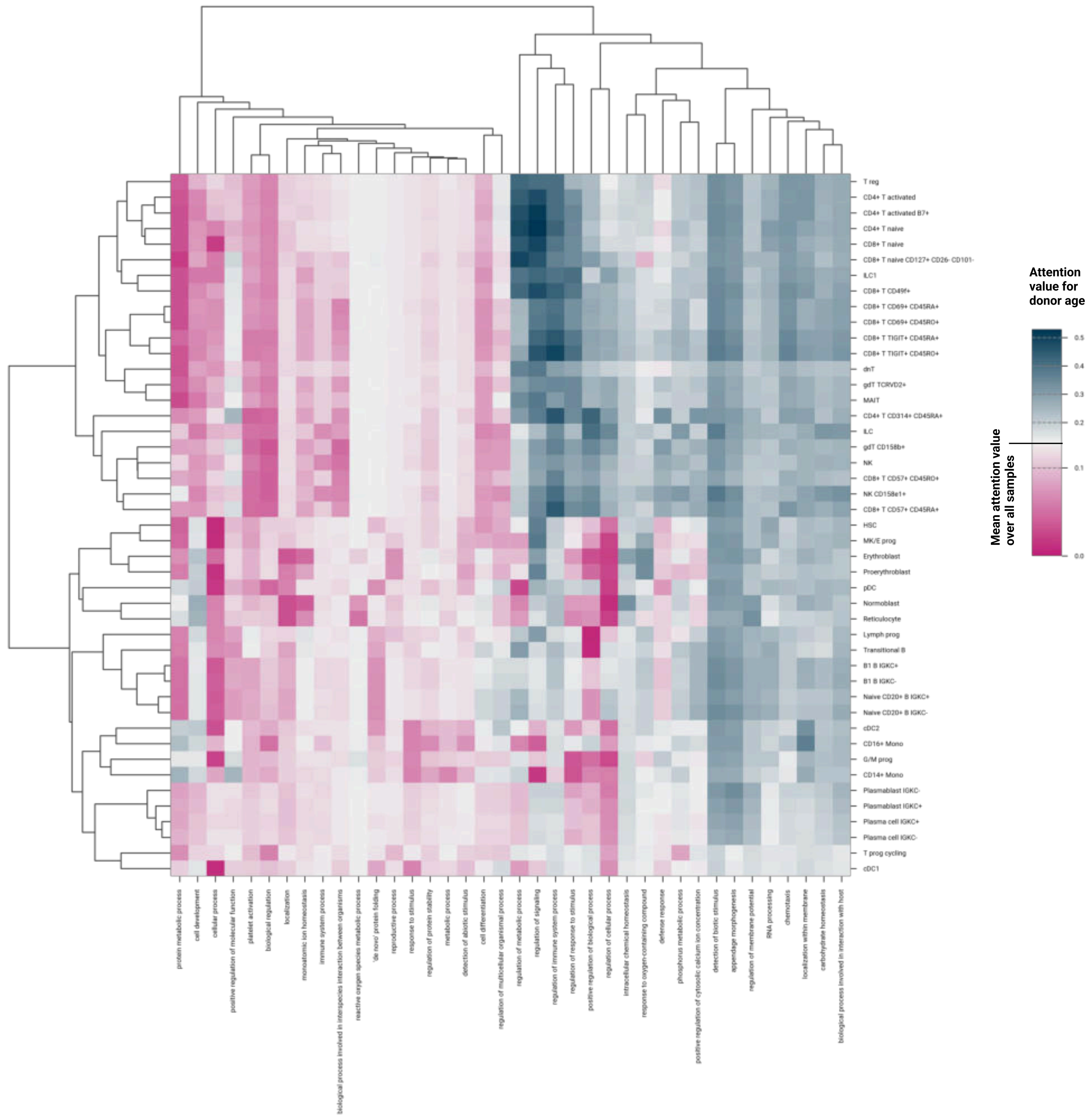


**Figure S24: GO term-specific covariate attention values indicate the association of T cells with aging-specific GO terms.**

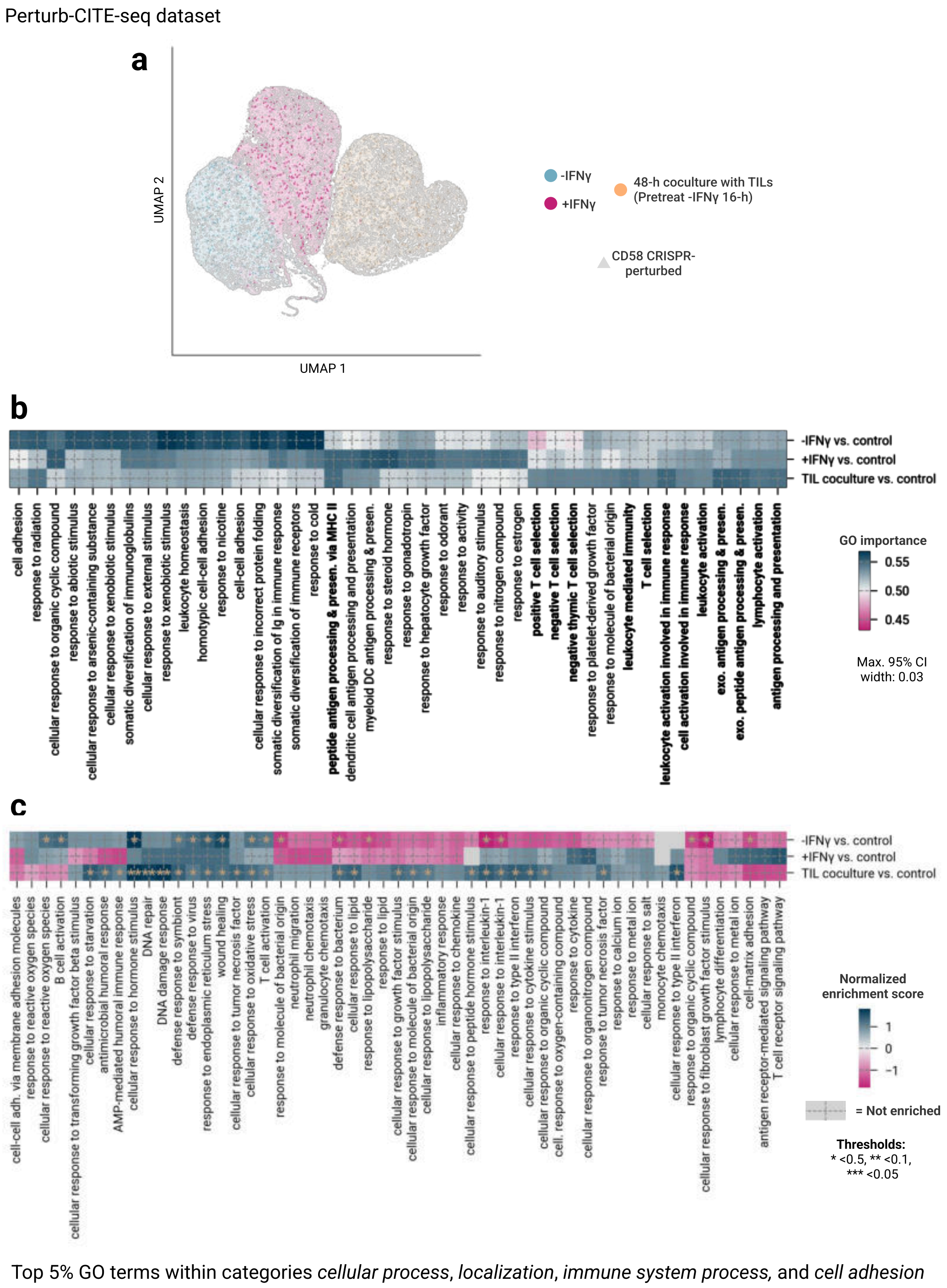


**Figure S25:** **a** Latent space generated using MultiVI of unperturbed and CD58 CRISPR-perturbed cells (trained only on unperturbed cells). **b** TNetworkVI GO importance scores and **c** GSEA normalized enrichment scores (NES) for CD58-perturbed cells in -IFN$\gamma$, +IFN$\gamma$ and TIL coculture screens (top 5% GO terms within the categories *cellular process*, *localization*, *immune system process*, and *cell adhesion*).

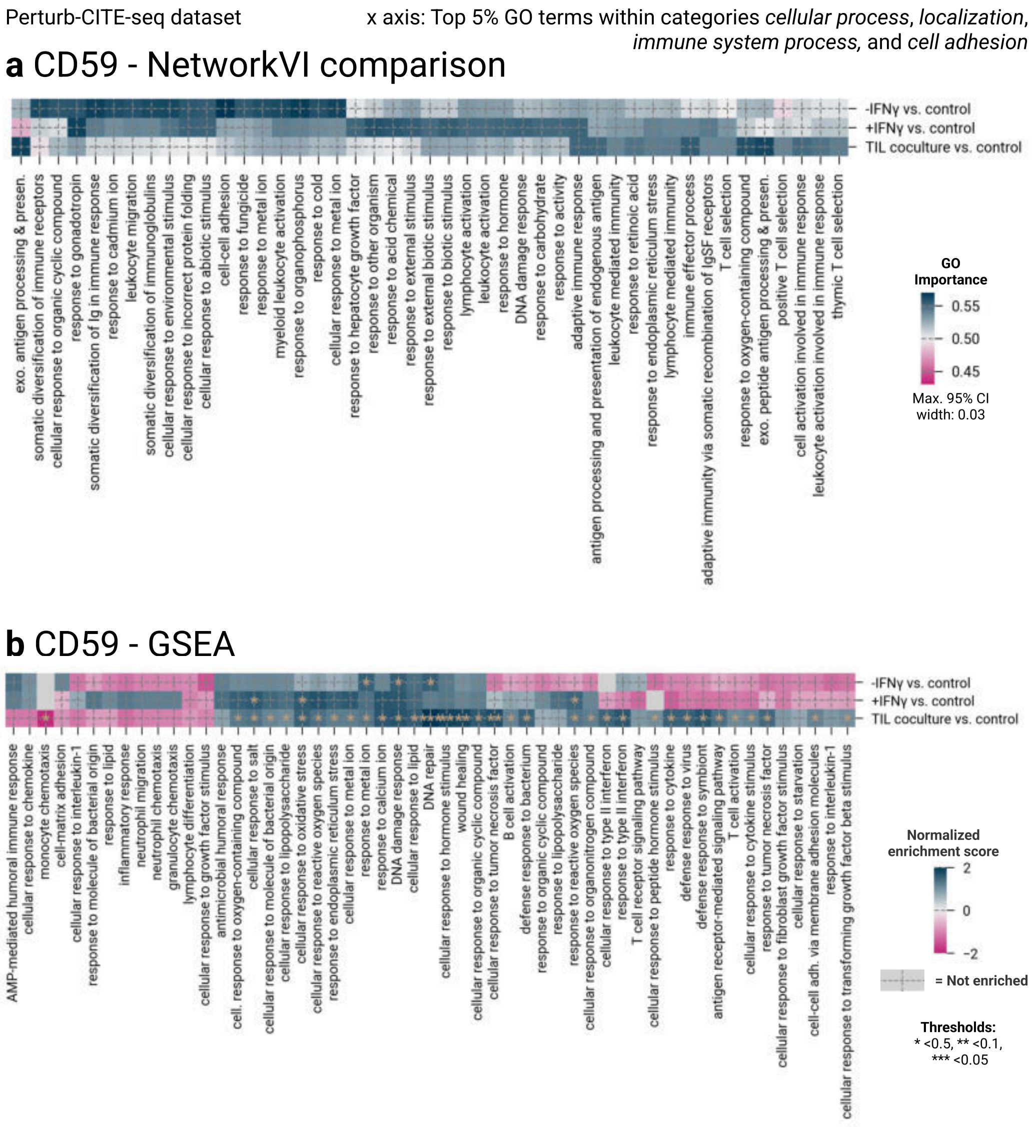


**Figure S26:** *See caption at the end of the figure.*

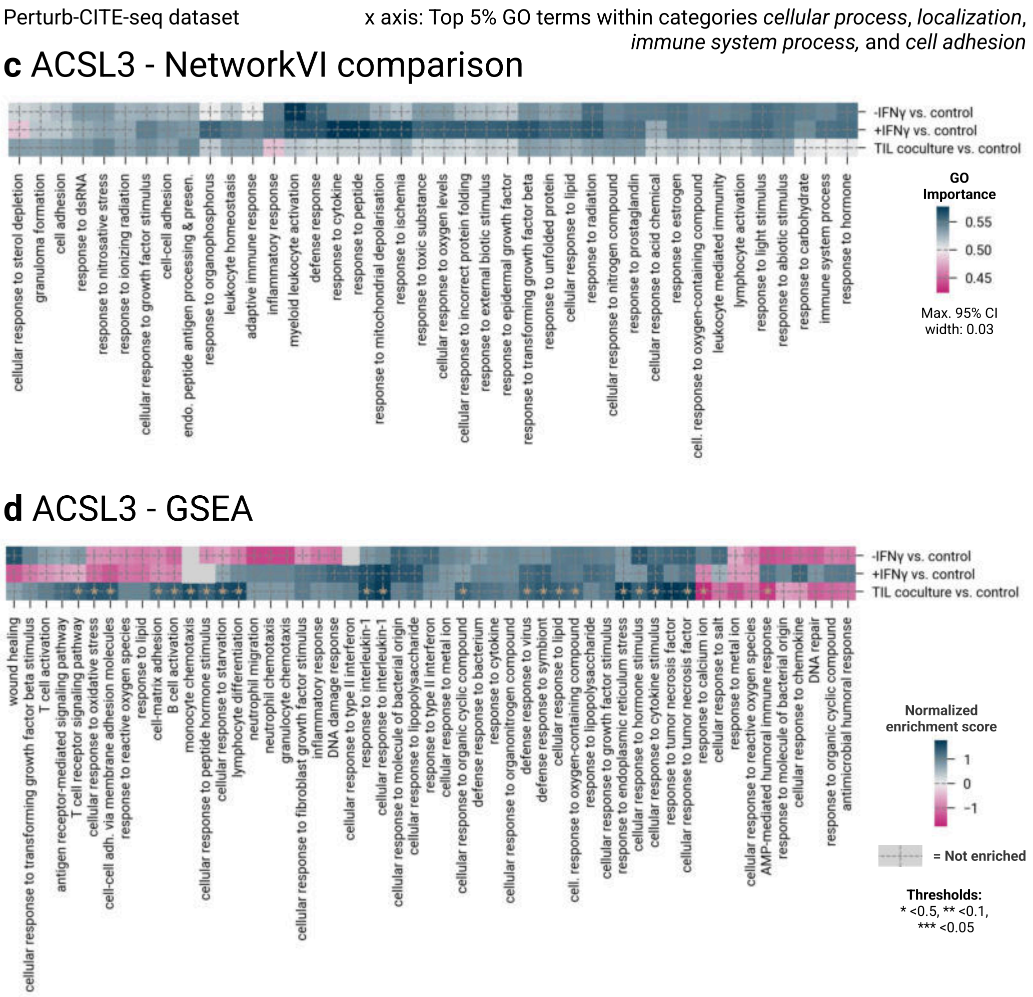


**Figure S26:** *See caption at the end of the figure.*

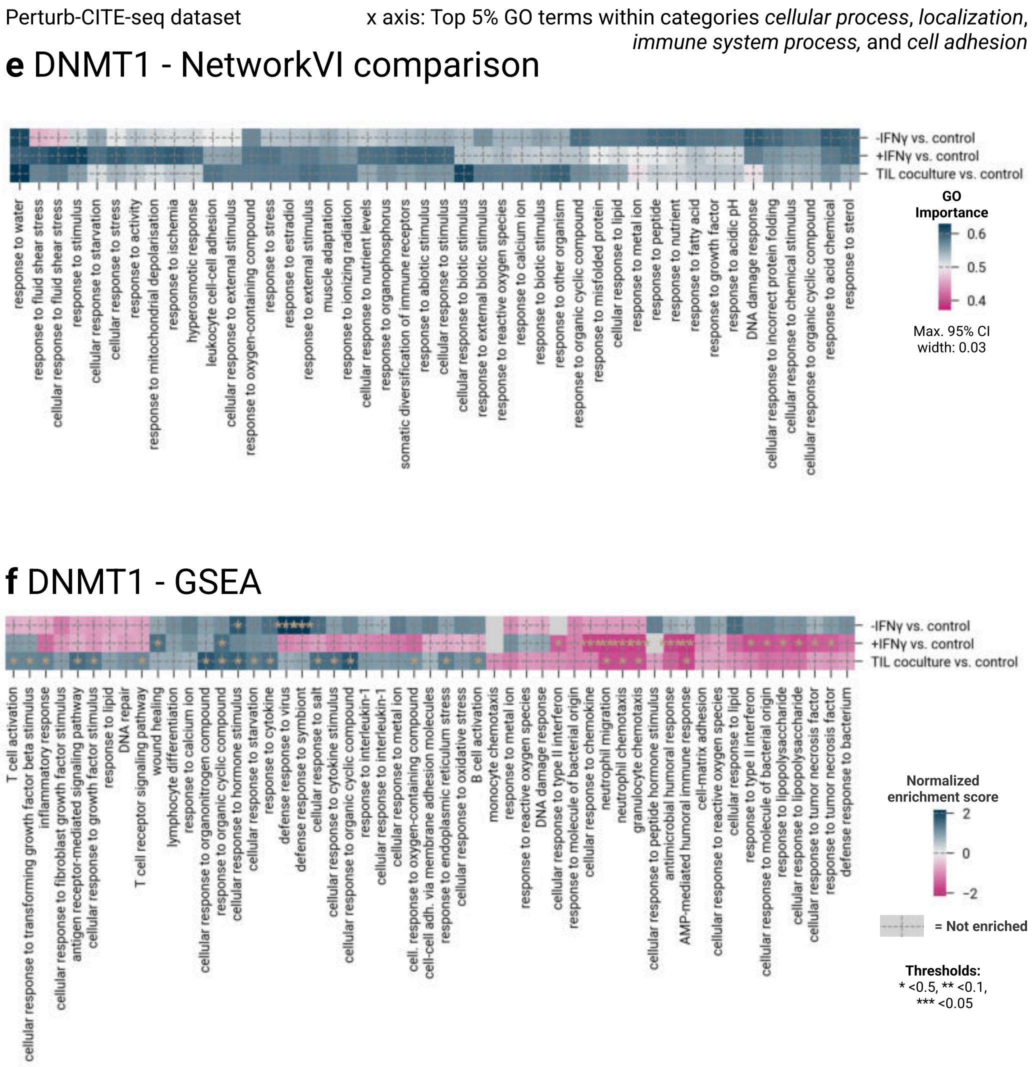


**Figure S26:** *See caption at the end of the figure.*

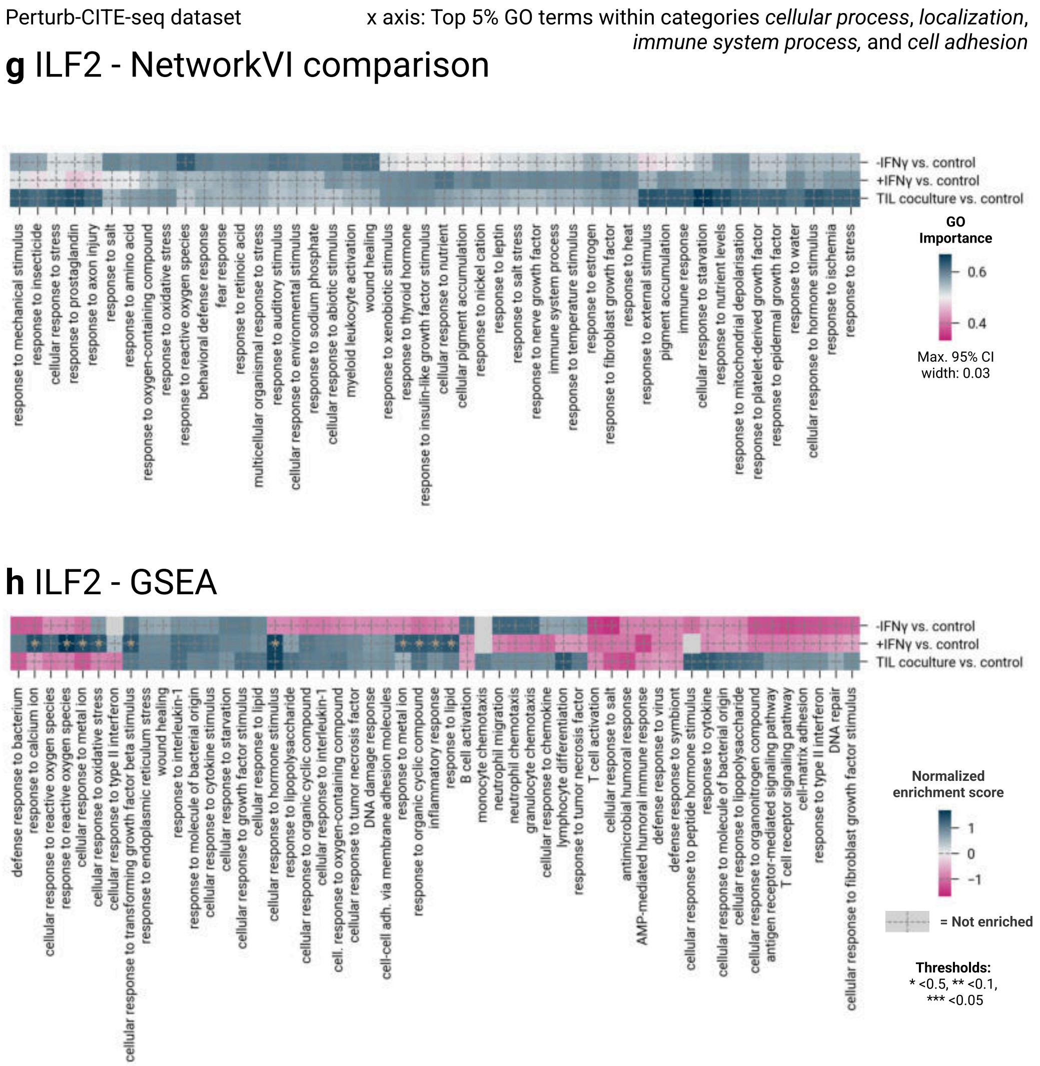


**Figure S26:** *See caption at the end of the figure.*

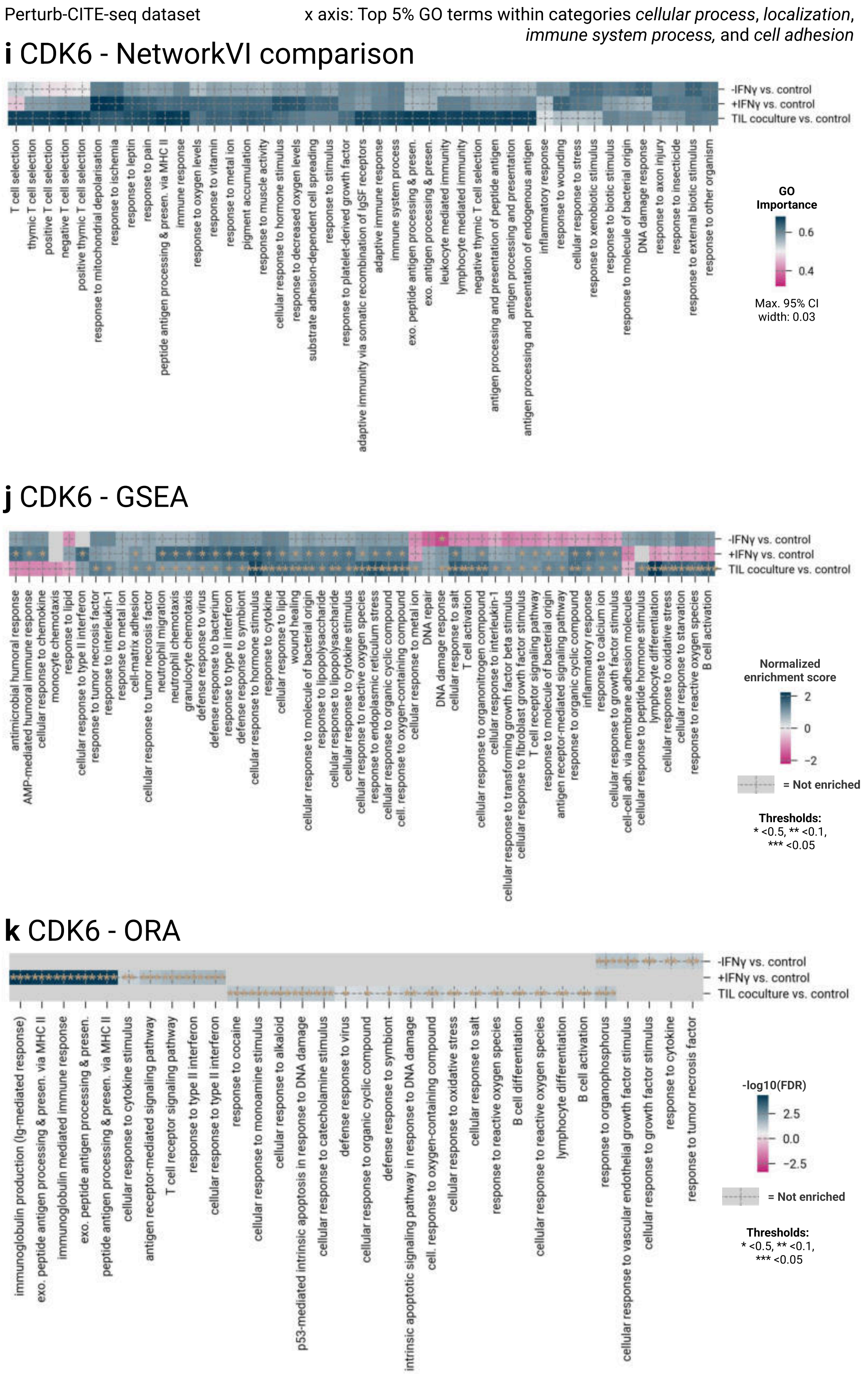


**Figure S26:** *See caption at the end of the figure.*

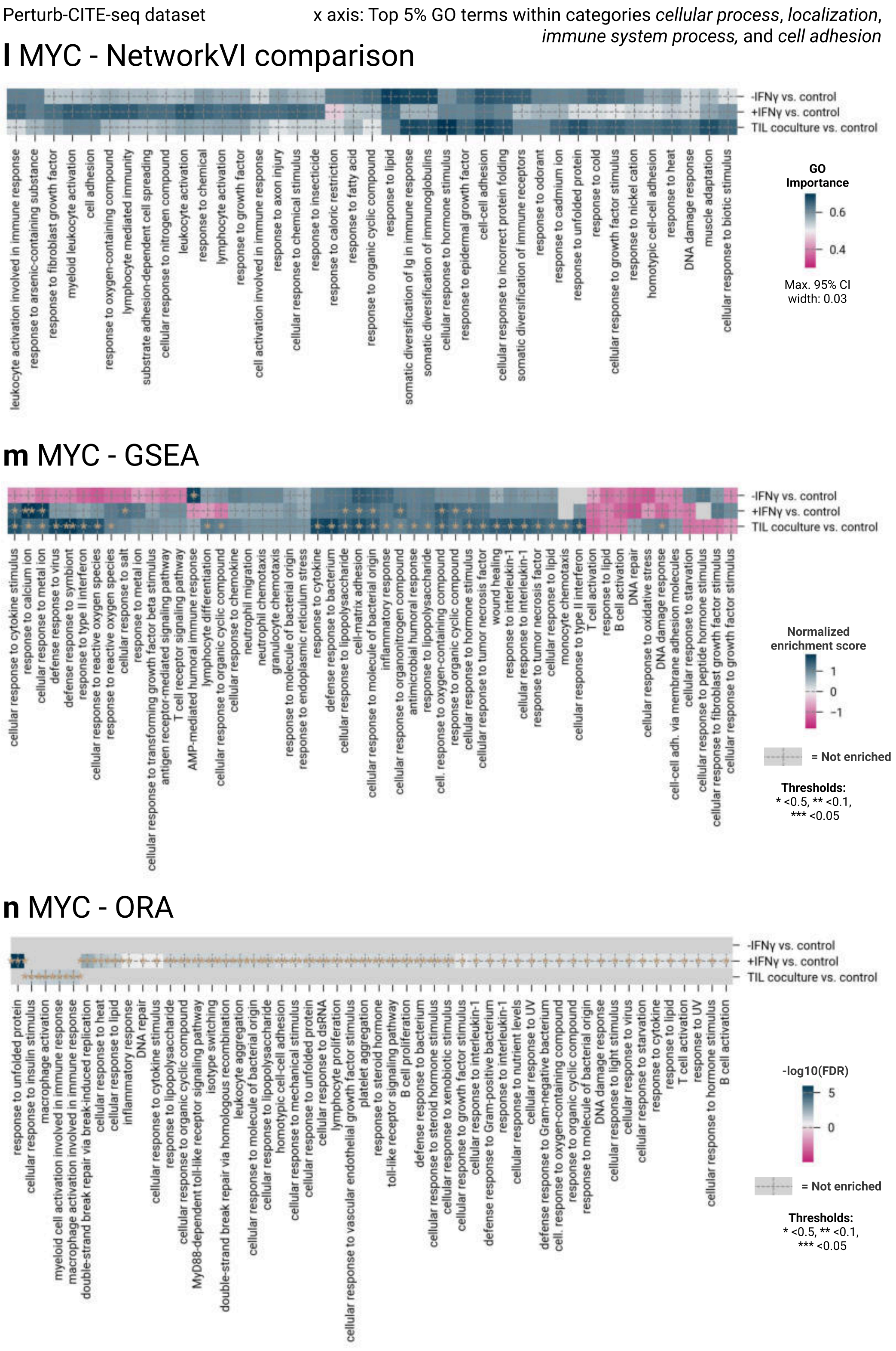


**Figure S26:** *See caption at the end of the figure.*

**Figure S26:** GO importance analysis/ GSEA and ORA (differentially expressed genes, Wilcoxon test, Benjamini-Hochberg FDR < 0.05, |logFC| > 0.5) analysis of CD59, ACSL3, DNMT1, ILF2, CDK6 and MYC perturbations in the [37] dataset. (Also see Figure 6d; Supplementary Note3), applying pairwise z-tests between conditions for each modelled GO term. NetworkVI and GSEA identify broadly consistent biological themes across all six additional genes. CDK6 shows the clearest screen-specific signal, with antigen presentation and T cell selection terms concentrated in the TIL coculture condition in NetworkVI, consistent with its documented role in repressing MHC class I genes [41], whereas GSEA distributes these terms more evenly across conditions and returns additional non-specific terms, including wound healing and DNA damage processes. For ICR-repressing genes with more distributed transcriptional effects (MYC, ACSL3, ILF2, and DNMT1) neither method yields a sharply localised signal; NetworkVI reflects this transparently through AUROC values near the random baseline rather than returning false-positive enrichments. Together, these results confirm that the pattern observed for CD58 is representative of the dataset-wide effect size rather than a selected outlier.

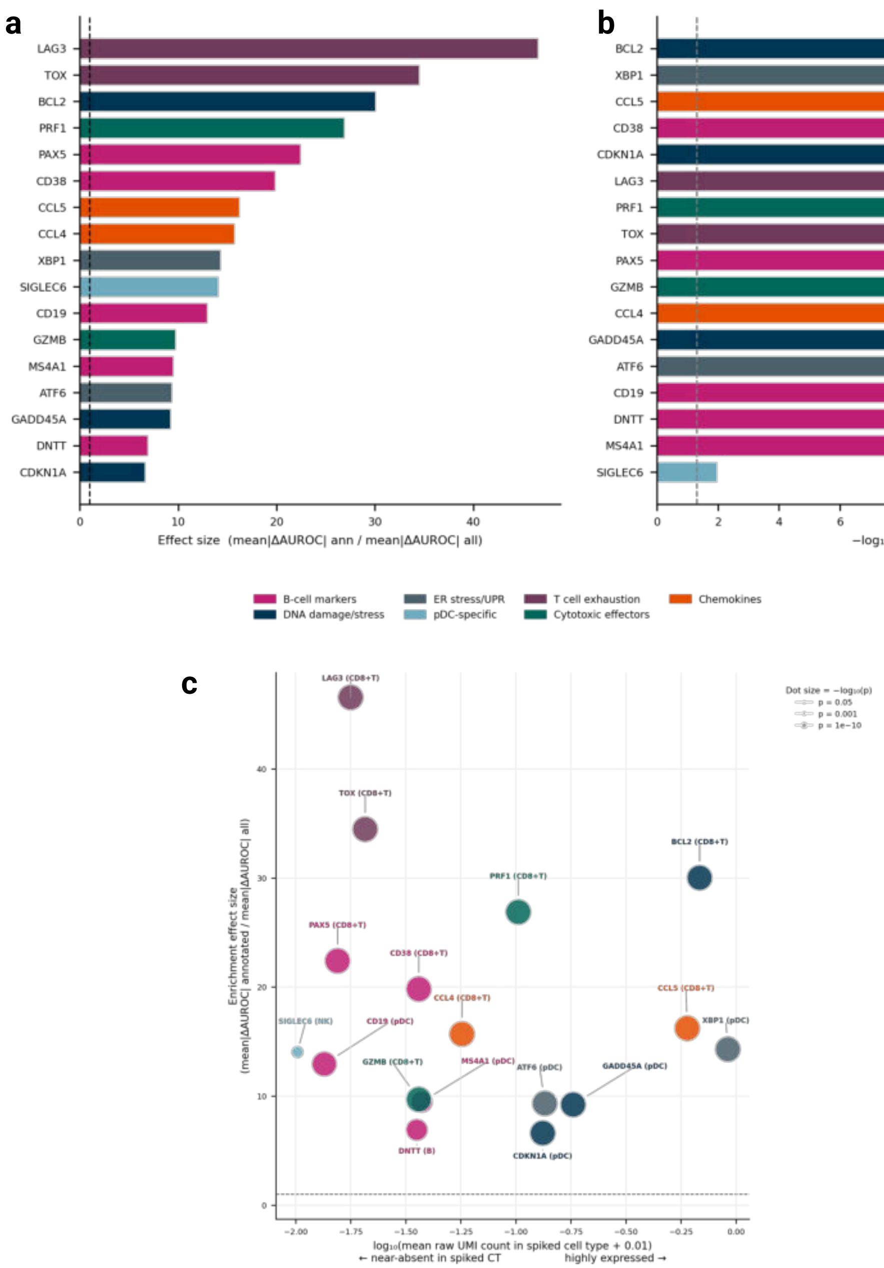


**Figure S27:** *See caption at the end of the figure.*

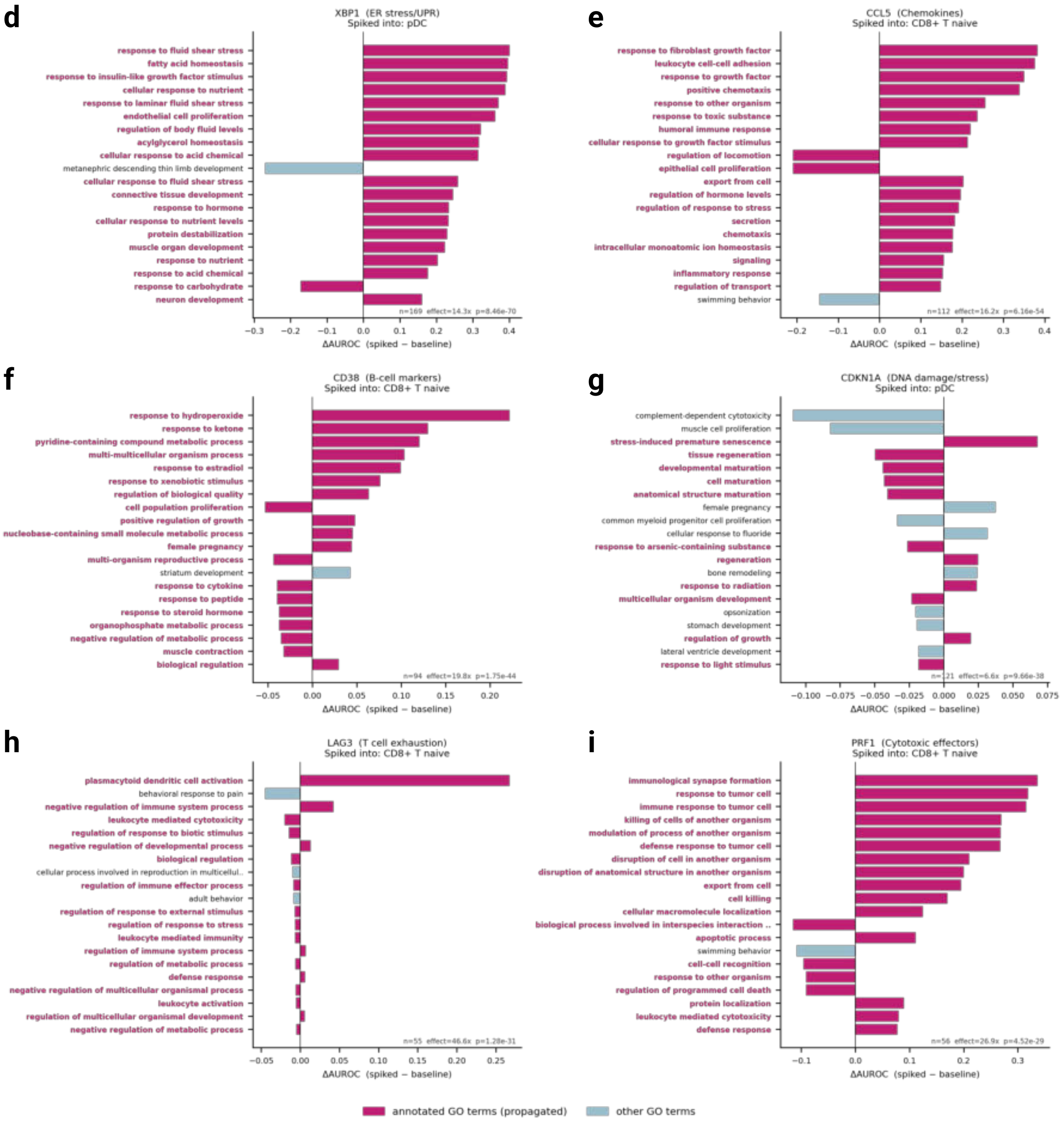


**Figure S27: Spike-in positive-control validation of NetworkVI gene-GO annotation routing.** **a** Enrichment effect sizes (mean |ΔAUROC| annotated GO terms / mean |ΔAUROC| all GO terms) for all 17 spike-in experiments, ranked by effect size. Each bar represents one gene spiked into its designated target cell type. Colors indicate biological group. The dashed vertical line marks an effect size of 1.0 (no enrichment relative to background). **b** Statistical significance ($-\log_{10}$ p-value, one-sided Mann-Whitney U test, annotated terms vs. all 1,791 GO terms) for all 17 experiments. The dashed vertical line marks p = 0.05. All 17 experiments reach significance; 16 of 17 reach p < 0.001. **c** Expression level in the spiked cell type (x-axis: $\log_{10}$(mean raw UMI count + 0.01)) versus enrichment effect size (y-axis). Dot size is proportional to $-\log_{10}$(p-value). No positive correlation between baseline expression and enrichment strength is observed: genes near-absent in the spiked cell type (e.g. LAG3: 0.7%, effect = 46.6x) show the strongest enrichment when the spike-in introduces a novel routing signal, while constitutively expressed genes (e.g. XBP1: 49.0%, effect = 14.3x) show smaller incremental shifts because their GO routing is already established at baseline. **d-i** Ranked ΔAUROC bar charts for six representative experiments, one per biological group (cytotoxic effectors: PRF1; chemokines: CCL5; DNA damage/senescence: CDKN1A; ER stress/UPR: XBP1; B cell markers: CD38; T cell exhaustion: LAG3). Each bar represents one of the model's 1,791 GO Biological Process latent dimensions, ranked by absolute ΔAUROC in the spiked cell type. GO terms annotated to the spiked gene (with ancestor propagation) are highlighted in pink; all other GO terms in light blue. Bold pink tick labels identify annotated GO terms. X-axis scaling is adaptive.

#### Supplementary Tables

**Table S1: Effect of stacking multiple TAD co-regulation layers on combined scib integration score.** Each entry reports the combined scib score on the NeurIPS 2021 CITE-seq BMMC dataset for one-, two-, and three-layer configurations of the gene co-regulation layer, across six gene-gene interaction sources. For the lineage-matched K562 TAD map, adding a second or third residual layer consistently degrades performance: the only additional connections introduced by deeper TAD layers would link genes whose TADs overlap at boundaries, pairs with no established direct co-regulatory relationship, providing misleading signal. The position-shifted K562 control shows no systematic degradation, consistent with the absence of biologically coherent co-regulatory structure to be disrupted. STRING similarly suffers with additional layers, reinforcing that dense, undirected, non-specific interaction networks do not benefit from deeper aggregation. By contrast, the TF-target interaction sources (TFLink, hTFtarget) and the GRAND K562 gene regulatory network show partial benefit from a second layer: these databases contain far fewer and more specific connections than STRING, and second-order TF-target-target paths can capture genuine indirect regulatory relationships. The number of co-regulation layers is exposed as a user-configurable parameter to allow dataset- and source-specific tuning.

| Layers | K562 | K562 Shifted | STRING | GRAND K562 | TFLink | hTFtarget |
|---|---|---|---|---|---|---|
| Single gene layer | 0.576 | 0.537 | 0.566 | 0.512 | 0.489 | 0.486 |
| Two gene layers | 0.498 | 0.544 | 0.556 | 0.558 | 0.556 | 0.563 |
| Three gene layers | 0.484 | 0.506 | 0.469 | 0.568 | 0.470 | 0.551 |

**Table S2: List of contact domain ENCODE datasets based on Hi-C for gene-gene interactions modeling.**

| Experiment | Bed file | Cell line | Datasets used for in study |
|---|---|---|---|
| ENCSR410MDC | ENCFF203AKP | GM12878 | DOGMA-seq [1] and CITE-seq dataset by [3] |
| ENCSR545YBD | ENCFF173VDJ | K562 | NeurIPS 2021 CITE and Multiome BMMC [2] |
| ENCSR660LPJ | ENCFF164AGX | MCF7 | Ablation in CITE-seq dataset by [3] |
| ENCSR549MGQ | ENCFF804SET | T47D | Ablation in CITE-seq dataset by [3] |
| ENCSR779QHO | ENCFF223RVL | ICR-80 | Lung dataset Tabula Sapiens by [18] |
| ENCSR549VZO | ENCFF549OBE | HepG2 | Liver dataset Tabula Sapiens by [18] |
| ENCSR543LYI | ENCFF958WAA | HCT116 | Large intestine dataset Tabula Sapiens by [18] |

**Table S3: Quantities of interactions between GO Layers in encoder for gene expression modality in DOGMA-seq dataset.** $gene_n$, $go_n$ is the number of neurons per gene or GO term. The number of features is 4000 genes mapping to 1749 GO terms with 664 genes mapping to none of the GO terms in the GO graph.

| | | | | |
|---|---|---|---|---|
| **GO Layer 2** (148*$go_n$) | 11 | | | |
| **GO Layer 3** (571*$go_n$) | 15 | 74 | | |
| **GO Layer 4** (1010*$go_n$) | 161 | 641 | 1248 | |
| **Gene Layer** (4000*$gene_n$) | 237 | 2745 | 9067 | 41040 |
| | **GO Layer 1** (19*$go_n$) | **GO Layer 2** (148*$go_n$) | **GO Layer 3** (571*$go_n$) | **GO Layer 4** (1010*$go_n$) |

**Table S4: Quantities of interactions between GO Layers in encoder for chromatin accessibility modality in DOGMA-seq dataset.** $gene_n$, $go_n$ is the number of neurons per gene or GO term. The number of features is 20000 peaks mapping to 2063 GO terms with 7467 peaks mapping to none of the GO terms in the GO graph.

| | **GO Layer 1** (19*$go_n$) | **GO Layer 2** (176*$go_n$) | **GO Layer 3** (674*$go_n$) | **GO Layer 4** (1193*$go_n$) |
|---|---|---|---|---|
| **GO Layer 2** (176*$go_n$) | 13 | | | |
| **GO Layer 3** (674*$go_n$) | 16 | 92 | | |
| **GO Layer 4** (1193*$go_n$) | 189 | 766 | 1474 | |
| **Gene Layer** (15745*$gene_n$) | 424 | 5855 | 17608 | 91117 |

**Table S5: Quantities of interactions between GO Layers in encoder for surface proteins modality in DOGMA-seq dataset.** $gene_n$, $go_n$ is the number of neurons per gene or GO term. The number of features is 210 surface proteins mapping to 841 GO terms with 9 surface proteins mapping to none of the GO terms in the GO graph.

| | **GO Layer 1** (19*$go_n$) | **GO Layer 2** (97*$go_n$) | **GO Layer 3** (260*$go_n$) | **GO Layer 4** (464*$go_n$) |
|---|---|---|---|---|
| **GO Layer 2** (97*$go_n$) | 9 | | | |
| **GO Layer 3** (260*$go_n$) | 10 | 37 | | |
| **GO Layer 4** (464*$go_n$) | 109 | 298 | 569 | |
| **Gene Layer** (192*$gene_n$) | 41 | 246 | 902 | 3803 |

**Table S6: Hyperparameters used in ax hyperparameter search for NetworkVI for dataset benchmarking and default parameter inference.**

| **Hyperparameter** | **Hyperparameter Range** | **Default Value** |
|---|---|---|
| Learning Rate (`lr`) | [1e-6, 1e-5, 1e-4, 1e-3] | 1e-4 |
| Modality Penalty (`modality_penalty`) | ["Jeffreys", "MMD"] | Jeffreys |
| Number of Encoder Layers (`n_layers_encoder`) | [2, 3, 4, 5] | 2 |
| Number of Decoder Layers (`n_layers_decoder`) | [2, 3, 4, 5] | 2 |
| Number of Hidden Units (`n_hidden`) | [128, 256, 384, 512, 640, 768, "Auto-inferred"] | Auto-inferred |
| Gene Likelihood (`gene_likelihood`) | ["zinb", "nb", "poisson"] | zinb |
| Dropout Rate (`dropout_rate`) | [0.0, 0.1, 0.2, 0.3] | 0.1 |
| Activation Function (`activation_fn`) | ["relu", "leaky_relu", "swish"] | relu |
| Standard GO Size (`standard_go_size`) | [1, 2, 3, 4, 5, 6] | 2 |
| Standard Gene Size (`standard_gene_size`) | [1, 2, 3, 4, 5, 6] | 5 |

**Table S7: Ablation of GO hierarchy hyperparameters on the NeurIPS 2021 CITE-seq BMMC dataset (fully paired).** Each row corresponds to a single-parameter sweep in which one architectural hyperparameter was varied while the others were held at their default values (`go_depth` = 4, `standard_go_size` = 2, `standard_gene_size` = 5). The combined scib score is computed as $0.6 \times$ biological variance conservation $+ 0.4 \times$ batch correction, following the weighting used throughout this study. The default configuration (`go_depth` = 4, `standard_go_size` = 2, `standard_gene_size` = 5) achieves a combined scib score of **0.567**, which equals or exceeds all single-parameter alternatives and was therefore adopted as the default.

| Configuration | GO hierarchy depth | GO node size | Gene node size |
|---|---|---|---|
| *Varying GO hierarchy depth (standard_go_size = 2, standard_gene_size = 5)* | | | |
| depth = 3 | 3 | 2 | 5 |
| depth = 4 **(default)** | **4** | **2** | **5** |
| depth = 5 | 5 | 2 | 5 |
| depth = 6 | 6 | 2 | 5 |
| *Varying GO node size (go_depth = 4, standard_gene_size = 5)* | | | |
| go_size = 2 **(default)** | 4 | **2** | 5 |
| go_size = 3 | 4 | 3 | 5 |
| go_size = 4 | 4 | 4 | 5 |
| go_size = 5 | 4 | 5 | 5 |
| go_size = 6 | 4 | 6 | 5 |
| *Varying gene node size (go_depth = 4, standard_go_size = 2)* | | | |
| gene_size = 2 | 4 | 2 | 2 |
| gene_size = 3 | 4 | 2 | 3 |
| gene_size = 4 | 4 | 2 | 4 |
| gene_size = 5 **(default)** | 4 | 2 | **5** |
| gene_size = 6 | 4 | 2 | 6 |

| Configuration | Combined scib score |
|---|---|
| *Varying GO hierarchy depth* | |
| depth = 3 | 0.526 |
| depth = 4 **(default)** | **0.567** |
| depth = 5 | 0.535 |
| depth = 6 | 0.499 |
| *Varying GO node size* | |
| go_size = 2 **(default)** | **0.567** |
| go_size = 3 | 0.567 |
| go_size = 4 | 0.567 |
| go_size = 5 | 0.487 |
| go_size = 6 | 0.555 |
| *Varying gene node size* | |
| gene_size = 2 | 0.508 |
| gene_size = 3 | 0.473 |
| gene_size = 4 | 0.493 |
| gene_size = 5 **(default)** | **0.567** |
| gene_size = 6 | 0.518 |

**Table S8: Hyperparameters used in ax hyperparameter search for MultiVI for DOGMA-seq dataset benchmarking.**

| Hyperparameter | Hyperparameter Range |
|---|---|
| Learning Rate (`lr`) | [1e-6, 1e-5, 1e-4, 1e-3] |
| Modality Penalty (`modality_penalty`) | ["Jeffreys", "MMD"] |
| Number of Encoder Layers (`n_layers_encoder`) | [2, 3, 4, 5] |
| Number of Decoder Layers (`n_layers_decoder`) | [2, 3, 4, 5] |
| Number of Hidden Units (`n_hidden`) | [128, 256, 384, 512, 640, 768, "Auto-inferred"] |
| Gene Likelihood (`gene_likelihood`) | ["zinb", "nb", "poisson"] |
| Dropout Rate (`dropout_rate`) | [0.0, 0.1, 0.2, 0.3] |
| Activation Function (`activation_fn`) | ["relu", "leaky_relu", "swish"] |

**Table S9: Hyperparameters used in ax hyperparameter search for MultiMIL for DOGMA-seq dataset benchmarking.**

| Hyperparameter | Hyperparameter Range |
|---|---|
| Learning Rate (`lr`) | [1e-6, 1e-5, 1e-4, 1e-3] |
| Dropout Rate (`dropout`) | [0.0, 0.1, 0.2, 0.3] |
| Number of Encoder Layers (`n_layers_encoders`) | [2, 3, 4, 5, "Auto-inferred"] |
| Number of Decoder Layers (`n_layers_decoders`) | [2, 3, 4, 5, "Auto-inferred"] |
| Number of Hidden Units in Encoders (`n_hidden_encoders`) | [128, 256, 384, 512, "Auto-inferred"] |
| Number of Hidden Units in Decoders (`n_hidden_decoders`) | [128, 256, 384, 512, "Auto-inferred"] |
| Activation Function (`activation`) | ["leaky_relu", "tanh"] |

**Table S10: Hyperparameters, ranges, and default values for TotalVI for DOGMA-seq dataset benchmarking.**

| Hyperparameter | Range | Default Value |
|---|---|---|
| Learning Rate (`lr`) | [1e-6, 1e-5, 1e-4, 1e-3] | 0.0001 |
| Number of Encoder Layers (`n_layers_encoder`) | [2, 3, 4, 5] | 2 |
| Number of Decoder Layers (`n_layers_decoder`) | [1, 2, 3, 4, 5] | 1 |
| Number of Hidden Units (`n_hidden`) | [128, 256, 384, 512] | 256 |
| Gene Likelihood (`gene_likelihood`) | ["zinb", "nb"] | "nb" |
| Dropout Rate for Encoder (`dropout_rate_encoder`) | [0.0, 0.1, 0.2, 0.3] | 0.2 |
| Dropout Rate for Decoder (`dropout_rate_decoder`) | [0.0, 0.1, 0.2, 0.3] | 0.2 |
| Activation Function (`activation_fn`) | ["relu", "leaky_relu", "swish"] | "relu" |

**Table S11: List of ENCODE p-value bigWig files** (Criteria: K562 cell line, generated by the Synder lab, aligned to GRCh38, labeled as released, one of the categories "transcription factors", "chromatin remodelers", "RNA-binding proteins", "histones", "cohesins", "cofactors", "DNA repair proteins", and "components of the DNA replication machinery".)

| | | | | |
|---|---|---|---|---|
| ENCFF596FVK | ENCFF215TUE | ENCFF704HVH | ENCFF037QZG | ENCFF914ZIB |
| ENCFF335JJU | ENCFF952ZRK | ENCFF581LZY | ENCFF691QRL | ENCFF143PFU |
| ENCFF135DIS | ENCFF671NDS | ENCFF331URE | ENCFF113JYQ | ENCFF198WJK |
| ENCFF448BAC | ENCFF326DQL | ENCFF258ASA | ENCFF461WJA | ENCFF955QWG |
| ENCFF365EJL | ENCFF032ORM | ENCFF760GUS | ENCFF806AVD | ENCFF259JPA |
| ENCFF240UFL | ENCFF934VTV | ENCFF657ROM | ENCFF213RKS | ENCFF706ROY |
| ENCFF612JUY | ENCFF843DKK | ENCFF774SCK | ENCFF273RVE | ENCFF516AAU |
| ENCFF070PUZ | ENCFF337XFG | ENCFF497LNO | ENCFF244MOS | ENCFF384BTU |
| ENCFF551KHO | ENCFF474CNJ | ENCFF764MXC | ENCFF128CFI | ENCFF974SRQ |
| ENCFF227WQI | ENCFF466GFK | ENCFF016WRT | ENCFF274ZIX | ENCFF386AWZ |
| ENCFF574RND | ENCFF995HPN | ENCFF503UFO | ENCFF541CKQ | ENCFF130GMP |
| ENCFF712SIB | ENCFF208ICL | ENCFF164TRU | ENCFF832GNG | ENCFF404COW |
| ENCFF564JOD | ENCFF747YVI | ENCFF655ZSU | ENCFF768NOJ | ENCFF808EXL |
| ENCFF671HXG | ENCFF462EMG | ENCFF288LPW | ENCFF921ISO | ENCFF260GXL |
| ENCFF370VNR | ENCFF638ZCD | ENCFF157RTJ | ENCFF033EVS | ENCFF004ZMA |
| ENCFF103NZN | ENCFF155FOH | ENCFF347WBJ | ENCFF845HTM | ENCFF395BVM |
| ENCFF454NLI | ENCFF514TNL | ENCFF970PUH | ENCFF037VAZ | ENCFF407OAJ |
| ENCFF342DXD | ENCFF336UPT | ENCFF835GXP | ENCFF140BXD | ENCFF800QUT |
| ENCFF773GNR | ENCFF425DBM | ENCFF817NXE | ENCFF933LNW | ENCFF416PCN |
| ENCFF875QZF | ENCFF118RGH | ENCFF448GFY | ENCFF336PZX | ENCFF818MPI |
| ENCFF288PSZ | ENCFF722KTI | ENCFF928BKT | ENCFF432OUZ | ENCFF721HET |
| ENCFF715HYZ | ENCFF440CYL | ENCFF405ELK | ENCFF780JRJ | ENCFF601NPD |
| ENCFF124PXB | ENCFF489SEA | ENCFF636FAG | ENCFF916NBA | ENCFF442IDG |
| ENCFF582NAG | ENCFF462UOM | ENCFF498WCM | ENCFF107GXG | ENCFF805EHS |
| ENCFF272LUB | ENCFF935PTG | ENCFF832TMB | ENCFF112BYB | ENCFF511RUY |
| ENCFF676UBU | ENCFF148GVM | ENCFF200NGJ | ENCFF375KUG | ENCFF941JUQ |
| ENCFF243OUW | ENCFF082KZY | ENCFF699KLZ | ENCFF398LBP | ENCFF614TYG |
| ENCFF143RUK | ENCFF346YIF | ENCFF517EKS | ENCFF544TIU | ENCFF735YUX |
| ENCFF041TQQ | ENCFF483LYL | ENCFF847HGM | ENCFF836FEQ | ENCFF766VME |
| ENCFF160ZNA | ENCFF930IRA | ENCFF198SGR | ENCFF280QPM | ENCFF701JND |
| ENCFF864PWN | ENCFF155XQP | ENCFF756UVW | ENCFF643SJZ | ENCFF529QQI |
| ENCFF104GDH | ENCFF191CES | ENCFF114SLT | ENCFF419IBT | ENCFF431SAX |
| ENCFF106EDU | ENCFF127RII | ENCFF253NTK | ENCFF830KGJ | ENCFF190FPU |
| ENCFF530IWB | ENCFF519ZFG | ENCFF610LRU | ENCFF772GDY | ENCFF970SIA |
| ENCFF293XFC | ENCFF283CLF | ENCFF501PQF | ENCFF085WAW | ENCFF293QAG |
| ENCFF158VSI | ENCFF263XFB | ENCFF675MND | ENCFF191VQV | ENCFF022EWV |
| ENCFF371VGO | ENCFF263YZZ | ENCFF913ZJJ | ENCFF463IKW | ENCFF908UVY |
| ENCFF949AOV | ENCFF262TFX | ENCFF373JER | ENCFF918KYJ | ENCFF596CNE |
| ENCFF199FSE | ENCFF934YZP | ENCFF750TBY | ENCFF122XON | ENCFF743DXN |
| ENCFF232TBZ | ENCFF203IUO | ENCFF405DRC | ENCFF563WCK | ENCFF685GZT |
| ENCFF211RPU | ENCFF223CKV | ENCFF770WTY | ENCFF021HPB | ENCFF316PSV |
| ENCFF560VXL | ENCFF723VHQ | ENCFF810ZPC | ENCFF026NAA | ENCFF475BKW |
| ENCFF567RJL | ENCFF209IFC | ENCFF427YFE | ENCFF360BJK | ENCFF630LBW |
| ENCFF842WAK | ENCFF244ZVM | ENCFF723UZM | ENCFF013FZZ | ENCFF794REJ |
| ENCFF206NPF | ENCFF652ZLV | ENCFF855QPP | ENCFF854EIV | ENCFF145GDZ |
| ENCFF253PNH | ENCFF064SRL | ENCFF131CUG | ENCFF584OLL | ENCFF796VGZ |
| ENCFF110ZZE | ENCFF043WON | ENCFF408KTS | ENCFF692MSP | ENCFF846LZT |
| ENCFF442MRI | ENCFF397QRA | ENCFF503FPF | ENCFF548JIB | ENCFF478OGJ |
| ENCFF250RIG | ENCFF666PCT | ENCFF266ORM | ENCFF386SIX | ENCFF316KJE |
| ENCFF675JPF | ENCFF286IOU | ENCFF521YNL | ENCFF191RLR | ENCFF163YNA |
| ENCFF985YBO | ENCFF162QJJ | ENCFF408DGX | ENCFF740VPW | ENCFF127BNU |
| ENCFF670OSB | ENCFF879EOK | ENCFF396SRD | ENCFF585IAR | ENCFF098FKU |

**Table S11: List of ENCODE p-value bigWig files**.

| | | | | |
|---|---|---|---|---|
| ENCFF167PVP | ENCFF235XFA | ENCFF607UFI | ENCFF631RZW | ENCFF200DLV |
| ENCFF274UBU | ENCFF508RXH | ENCFF615VLU | ENCFF563XMY | ENCFF713BBY |
| ENCFF125SPZ | ENCFF169WSC | ENCFF271CFD | ENCFF551YRA | ENCFF684UYV |
| ENCFF971IQQ | ENCFF062OZT | ENCFF495YNJ | ENCFF770ATI | ENCFF913TWG |
| ENCFF402ULS | ENCFF003NIX | ENCFF705BCZ | ENCFF377DDT | ENCFF674URK |
| ENCFF495KOS | ENCFF405BON | ENCFF856CAV | ENCFF223RME | ENCFF650DYQ |
| ENCFF036TCU | ENCFF988HVV | ENCFF512NFV | ENCFF454RXI | ENCFF979OJR |
| ENCFF629KBJ | ENCFF768QRC | ENCFF132HZU | ENCFF438XUP | ENCFF600BUX |
| ENCFF060GMB | ENCFF854BXW | ENCFF306XYI | ENCFF843WTJ | ENCFF816TPI |
| ENCFF682XOQ | ENCFF145FWH | ENCFF064AVI | ENCFF552HWW | ENCFF264SMF |
| ENCFF494PCP | ENCFF733QSR | ENCFF473UBB | ENCFF798IJB | ENCFF834QAV |
| ENCFF780PZN | ENCFF199WRR | ENCFF573MHC | ENCFF331LJE | ENCFF598RUP |
| ENCFF269VGE | ENCFF547AGX | ENCFF690ABT | ENCFF703CTG | ENCFF152HYG |
| ENCFF313CYB | ENCFF403AZP | ENCFF226VDW | ENCFF430RCW | ENCFF854POL |
| ENCFF129BBR | ENCFF364OGY | ENCFF655SMR | ENCFF366SMI | ENCFF794BGU |
| ENCFF420CLL | ENCFF106ZRO | ENCFF744MAU | ENCFF230DRI | ENCFF358XWW |
| ENCFF511ZRC | ENCFF325ACI | ENCFF843FWI | ENCFF304UOP | ENCFF483RAR |
| ENCFF039GGQ | ENCFF807KVX | ENCFF431FTN | ENCFF565PQQ | ENCFF550YIC |
| ENCFF676NJU | ENCFF594LZQ | ENCFF389PED | ENCFF830VNI | ENCFF265WDL |
| ENCFF332INK | ENCFF422IWO | ENCFF547OVU | ENCFF184MJA | ENCFF034GGJ |
| ENCFF897MKD | ENCFF831QAC | ENCFF865XSE | ENCFF183GAS | |
| ENCFF346NDO | ENCFF163JPZ | ENCFF391FZA | ENCFF974KEW | |

**Supplementary Data**

**Supplementary Data 1 - 8:** scIB scores for paired and mosaic integration with default and hyper scib scores for the (1, 2) DOGMA-seq dataset, (3, 4) NeurIPS 2021 CITE BMMC dataset, (5, 6) NeurIPS 2021 Multiome BMMC dataset and (7, 8) Hao et al. CITE dataset.

**Supplementary Data 9:** GO importance scores (unfiltered) for CD58 perturbed cells in -IFN$\gamma$, +IFN$\gamma$ and TIL coculture screens.
